# Mixture Density Regression reveals frequent recent adaptation in the human genome

**DOI:** 10.1101/2021.12.20.473463

**Authors:** Diego F. Salazar-Tortosa, Yi-Fei Huang, David Enard

## Abstract

How much genome differences between species reflect neutral or adaptive evolution is a central question in evolutionary genomics. In humans and other mammals, the prevalence of adaptive versus neutral genomic evolution has proven particularly difficult to quantify. The difficulty notably stems from the highly heterogeneous organization of mammalian genomes at multiple levels (functional sequence density, recombination, etc.) that complicates the interpretation and distinction of adaptive vs. neutral evolution signals. Here, we introduce Mixture Density Regressions (MDRs) for the study of the determinants of recent adaptation in the human genome. MDRs provide a flexible regression model based on multiple Gaussian distributions. We use MDRs to model the association between recent selection signals and multiple genomic factors likely to affect positive selection, if the latter was common enough in the first place to generate these associations. We find that a MDR model with two Gaussian distributions provides an excellent fit to the genome-wide distribution of a common sweep summary statistic (iHS), with one of the two distributions likely enriched in positive selection. We further find several factors associated with recent adaptation, including the recombination rate, the density of regulatory elements in immune cells, GC-content, gene expression in immune cells, the density of mammal-wide conserved elements, and the distance to the nearest virus-interacting gene. These results support that strong positive selection was relatively common in recent human evolution and highlight MDRs as a powerful tool to make sense of signals of recent genomic adaptation.

**Author Summary:** Over the last 50,000 years, human populations have been exposed to selective pressures that can trigger adaptation in the genome. The search for signals of these selective events is however obscured by the substantial variation of factors that are relevant for the prevalence and detection of adaptation across the genome. Here, we analyze the impact of multiple factors on positive selection using a biologically meaningful approach that considers the influence of adaptative and non-adaptive processes in the human genome. Our results show that this novel approach is better suited to find genomic factors associated with recent positive selection compared to the classic regression or correlation approaches. We find multiple genomic functional elements associated with selection across the genome, including novel associations that emerge only after controlling for multiple confounding factors. This strongly suggests that adaptation was common enough in recent evolutionary times to produce a widespread correlation between functional elements and positive selection in the human genome.

**Note on the language used in this manuscript:** In this manuscript we use discrete population groups, such as the Yoruba. We want to emphasize that these discrete groups are only used for convenience and clarity when presenting our results, but in fact represent arbitrary human constructions, the same way that the boundaries of countries are arbitrary. There is an unbroken continuum and mixing of geographical ancestries across groups often identified as distinct populations across the world. The discrete groups we use are, as such, by no mean discrete genetic entities. Once grouped together, the grouped individuals only happen to be genetically more similar, with their ancestries coming from specific geographic locations more predominantly than individuals from the other groups. How much more predominantly is completely arbitrary.

## Introduction

The characteristics of the human genome can influence both the occurrence and the detection of recent positive selection in a specific genomic region. Similar to other mammals, the human genome has a complex organization, with highly heterogeneous recombination rates and distribution of functional elements (regulatory or coding) along chromosomes. This inherent heterogeneity in the factors that are likely to influence both the occurrence of positive selection and our ability to detect it, has complicated the study of the prevalence of recent adaptation in the human genome [1,2]. Another important complicating factor is that statistics of recent adaptation have complex distributions across the genome (see below), which limits the ability of classic correlation and regression approaches to analyze the genomic factors that contributed to recent genomic adaptation. There is nevertheless growing evidence from genomic scans suggesting that recent positive selection in the form of selective sweeps may have been relatively common during recent human evolution [3–9]. More recent and powerful selection scans using machine (deep) learning approaches suggest that a non-negligible proportion of the human genome may be affected by selective sweeps [10–13]. Advances on this topic will depend on our ability to understand what factors govern the local genomic rate of recent adaptation along human chromosomes. Indeed, if signatures of selective sweeps do not occur randomly, as expected if all sweep signals are false positives only reflecting genetic drift, but instead associate non-randomly with functional elements in the genome, then this provides evidence that sweeps had to be common enough in the first place to generate the association.

What factors then are *a priori* expected to matter for recent adaptation? Among all genomic factors, recombination is a key process determining the patterns of linkage disequilibrium between alleles across the human genome [14], that can strongly influence the probability to detect recent selection in the form of selective sweeps. The lower the recombination rate, the larger the genomic region where neutral variants will hitchhike to higher frequencies along with an advantageous variant, before recombination breaks down their linkage disequilibrium [15,16]. The higher the recombination rate, the smaller the genomic region affected by the sweep. Different sweep sizes in low and high recombination regions result in a higher statistical power to detect sweeps in low than in high recombination regions [8,17,18]. In parallel, the increase of linkage disequilibrium in low-recombination regions favors the appearance of larger neutral haplotypes making these regions more prone to detect false positive sweeps [17,18]. In addition, recombination rate influences the probability of deleterious variants interfering with the adaptive ones given the increased probability of linkage disequilibrium between them under low recombination [19–21]. This heterogeneity, together with other factors such as background selection, has contributed to a persistent debate around the prevalence of selective sweeps in the human genome [1,2].

If recent adaptation and selective sweeps in particular were relatively common during recent human evolution, we expect that other factors on top of recombination should be relevant for their occurrence and detection in the human genome. Generally, we expect that selective sweeps should occur more frequently around functional segments of the genome where adaptive mutations are expected to take place [1]. Therefore, we should find enough signals of positive association between selective sweeps and the overall density of functional elements, either coding or non-coding. In a pioneer analysis, Barreiro et al. (2008) showed for the first time a relationship between genome-wide signatures of positive selection and the distribution of functional elements. Among those highly differentiated Single Nucleotide Polymorphisms (SNPs), they found an excess of non-synonymous SNPs compared to non-genic SNPs. Note, however, that they did not explicitly control for the impact of background Selection (BGS), the coincident removal of neutral variants together with genetically linked deleterious mutations, which varies between the classes analyzed [2,23]. We also expect specific functions to be associated with an increased occurrence of selective sweeps. For example, the presence of genes coding for proteins that interact with pathogens, and viruses in particular (Virus-Interacting Proteins or VIPs), should influence the frequency of sweeps in a given genomic region, independently of other genomic features [9,24]. More generally, genomic regions with immune genes are expected to have experienced more sweeps [25–28]. Similarly, reproduction-related functions show signals of positive selection [5,29], thus tissues related with these functions should be associated with the frequency of sweeps.

Several summary statistics are currently available to detect recent genomic adaptation. Among all possible choices, the statistics that use the structure of haplotypes along chromosomes are the most appropriate to investigate the prevalence and determinants of recent positive selection. First, haplotype-based statistics have good statistical power to detect strong and recent incomplete sweeps [5,9,30,31]. Second, they are not confounded by BGS [1,32]. This insensitivity is a particularly important attribute when studying genomic regions subject to different levels of BGS. Given that multiple genomic factors correlate with this process, their association with selection can be confounded if the summary statistic used is also sensitive to BGS. In this regard, the integrated Haplotype Score (iHS) [5] is a haplotype-based statistic that has been extensively tested [1,5,8,25], and can detect selection signals both from *de novo* mutations and to some extent from standing genetic variation, provided that selection started from not too high initial allele frequency [9,30]. This summary statistic can be used to scan many genomes given the availability of fast implementations [33]. Therefore, we use iHS as a measure of recent positive selection across the human genome. As shown in Figure 1 (see also Methods), iHS has a complex, asymmetric distribution across the genome that is not well captured by classic distributions (Supplemental Results S1: Figure S1). This raises an important limitation of violated assumptions when using classic regression or correlation analysis to find the genomic determinants of iHS, with for example the assumption of statistics following a gaussian distribution clearly not met. Furthermore, classic regression and correlation analyses may not capture well the process of recent positive selection through selective sweeps measured by iHS. Indeed, classic regression and correlation assume a linear association between iHS and a factor contributing to recent positive selection in a distributed way across the entire genome. This neglects the localized nature of recent selection events, that have likely affected only a limited genomic portion, while leaving the rest of the genome (and likely substantial majority) without any correlation between iHS and a contributing factor, even though the latter did contribute to selection occurring locally in the genome. This creates a situation where only the higher values of a statistic such as iHS, those that are more likely to represent selection, might correlate at all with genomic factors influencing selection. In this respect, the distribution of iHS is particularly interesting, with an upper tail that is much heavier than the lower tail. We can hypothesize that localized selection in a limited portion of the genome might have generated this heavy tail, rather than a scenario where more generalized selection covering the entire genome would have shifted the whole distribution. What tools then can we use to properly account for this complex distribution of iHS?

**Figure 1.**
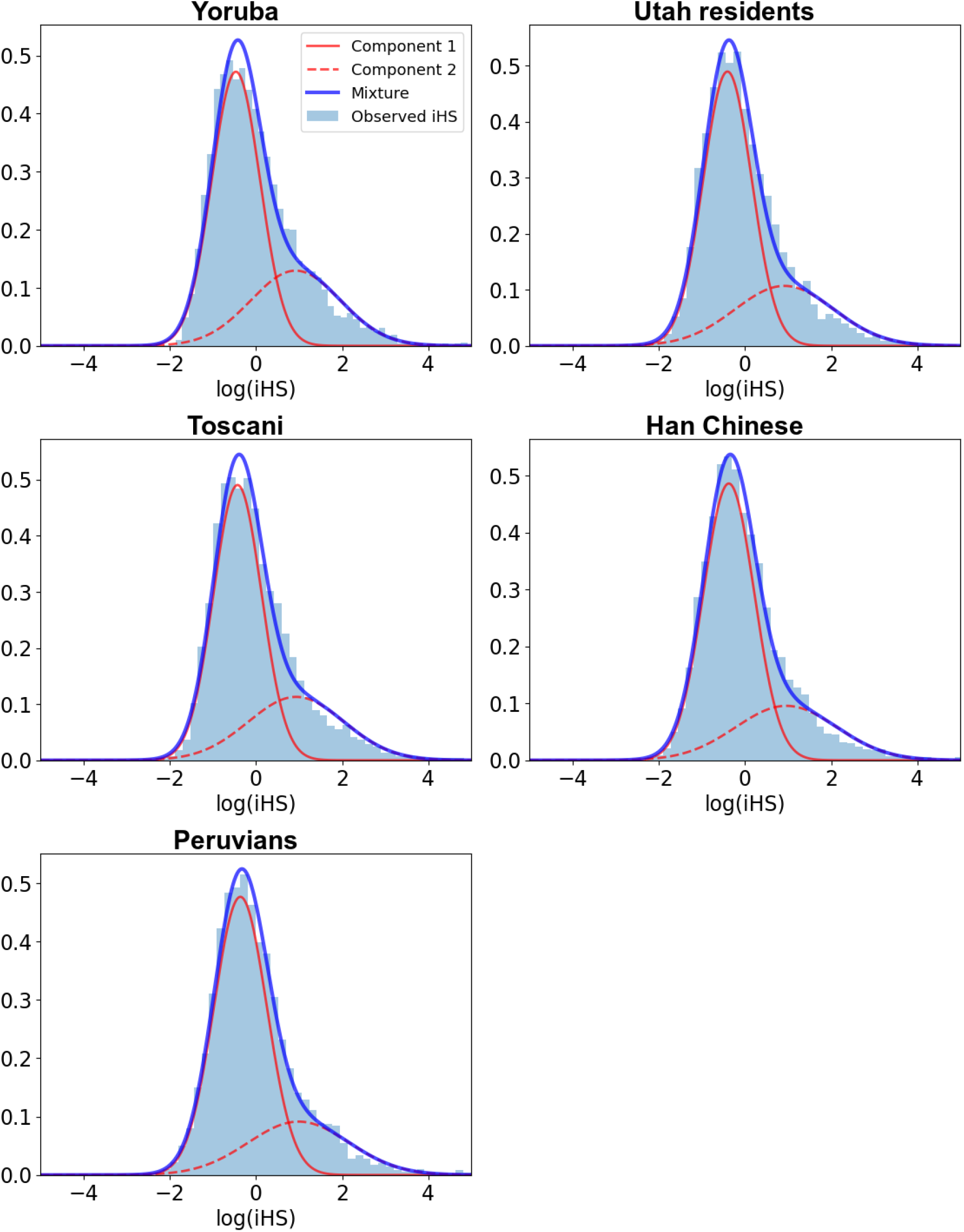
Mixture of Gaussian distributions fitting observed iHS (1,000 kb windows) for the five studied populations: A) Africa - Yoruba; B) Europe - Utah residents with Northern and Western European ancestry; C) Europe - Toscani; D) East Asia - Han Chinese; E) America - Peruvians. For each population, the figure shows two Gaussian distributions, components 1 and 2 of iHS, being the latter enriched in positive selection. In that component, iHS linearly depends on the genomic factors considered. Legend: Light blue = Observed iHS; Dark blue = Mixture model; Full red curve = Component 1 of the mixture model; Dashed red curve = Component 2 of the mixture model enriched in positive selection.

Here, we model the genomic determinants of recent positive selection in the human genome. We measure recent selective sweeps with the iHS statistic from five human populations represented in the 1,000 Genomes phase 3 dataset [34]. The iHS statistic has more power to detect recent and incomplete sweeps, while it has reduced power to detect complete sweeps or sweeps more than 30,000 years old [9,15]. Therefore, we restrict our analyses to selection that occurred after the main human migration out of Africa [35]. To account for the observed complexity of the distribution of iHS, we use Mixture Density Regressions (MDRs) to test the association between iHS and several genomic factors that are possible determinants of selection (Methods). MDRs can fit a mixture of several distributions to the observed statistic. In our case, we fit a mixture of two Gaussian-distributed components to the observed iHS (Methods). The first component fitting lower values of iHS is expected to match more the distribution of iHS under drift, the portion of the genome that was not affected by localized selection. The second component fitting higher iHS values is expected to be comparatively enriched in positive selection. Then, we look at the genomic factors that influence iHS in the regression model, interpreting the significant association of a genomic factor with the selection-enriched component as evidence for positive selection (Methods). This differs from other approaches trying to understand the determinants of selection. Previous studies have focused on the detection of classic partial correlations between selection and genomic factors (e.g., Lohmueller et al. [36]). Given that the impact of hitchhiking on the human genome is likely localized and not expected to influence the entire genome, the signal of recent sweeps could be diluted enough across the genome so that this classic approach misses true selection signals. An approach that specifically uses a selection-enriched component should have more power to reveal the impact of positive selection.

As expected according to previous studies, we find a strong association between recombination rate and recent positive selection signals in the human genome. In addition to recombination, we find that other factors, such as the density of regulatory elements in immune cells, GC-content, gene expression in immune cells, the density of mammal-wide conserved elements and the distance to VIPs, all show strong associations with recent positive selection. These results imply that recent positive selection had to be common enough to create these associations, and further highlight how MDRs can be used to clarify the prevalence and determinants of recent positive selection based on genome scans.

## Results

We first validate that the MDR with two gaussian distributions fits the distribution of iHS much better than a single gaussian (Supplemental Results S1: Figure S1). The tight, visually obvious fit between the observed iHS distribution and the two gaussians in our MDR model (Figure 1) also shows that more than two gaussians are very likely not needed and would negligibly improve the fit. We consider multiple genomic factors to model the association with recent selection signals. These factors are possible determinants of selection according to previous evidence [1,5,9,17,18,29,37,38]. We use a gene-centric perspective where each individual protein-coding gene in the human genome is associated with a set of factors measured by centering a genomic window on each gene (Methods). This approach is likely to detect the influence of factors that differ between genes, rather than only differences between genic and intergenic regions. The set of factors in the MDR model includes several genomic features, like the length of the gene at the center of the window or recombination estimates from the latest deCODE genetic map (Methods). We obtain other genomic factors, like the GC-content and the densities of genes, coding sequences and conserved elements. Moreover, we include the density of transcription factor binding sites according to the hypersensitivity to DNaseI and chromatin immunoprecipitation (ChIP-seq) experiments [39]. In the latter case, we also obtain the density of regulatory elements for specific subsets of cell lines: testis and immune cells. Table 1 provides a complete list of the factors considered, and the Methods section details how these factors were obtained.

**Table 1.** Slopes and p-values of the association between iHS and genomic factors for the five studied populations (Yoruba, Utah residents, Toscani, Han Chinese and Peruvians) in 1,000 kb windows within the selection-enriched component.

| Covariate | Yoruba |  | Utah residents |  | Toscani |  | Han Chinese |  | Peruvians |  |
| --- | --- | --- | --- | --- | --- | --- | --- | --- | --- | --- |
|  | Slope | p-value | Slope | p-value | Slope | p-value | Slope | p-value | Slope | p-value |
| Intercept | -1.43 | <1E-16 | -1.77 | <1E-16 | -1.66 | <1E-16 | -1.78 | <1E-16 | -1.55 | <1E-16 |
| Recombination rate | -2.44 | <1E-16 | -2.8 | <1E-16 | -2.7 | <1E-16 | -2.51 | <1E-16 | -1.74 | <1E-16 |
| GC-content | 0.55 | 1.34E-06 | 0.745 | 1.35E-08 | 0.516 | 2.47E-05 | 1.171 | <1E-16 | 0.351 | 2.63E-03 |
| Density of conserved elements | 0.159 | 6.40E-04 | 0.379 | 7.74E-13 | 0.337 | 2.70E-11 | 0.311 | 4.20E-10 | 0.057 | 2.43E-01 |
| Number PPIs | -0.07 | 4.25E-02 | -0.05 | 1.95E-01 | -0.04 | 2.67E-01 | -0.09 | 1.92E-02 | -0.04 | 3.00E-01 |
| Distance to VIPs | -0.17 | 4.16E-06 | -0.34 | 8.45E-14 | -0.27 | 1.92E-10 | -0.14 | 5.35E-04 | -0.18 | 2.77E-05 |
| Gene expression | -0.14 | 3.77E-02 | -0.34 | 9.70E-06 | -0.25 | 4.92E-04 | -0.25 | 1.14E-03 | -0.26 | 2.56E-04 |
| Gene expression in immune cells | 0.265 | 1.51E-05 | 0.408 | 6.87E-09 | 0.366 | 5.34E-08 | 0.307 | 9.51E-06 | 0.251 | 1.56E-04 |
| Gene expression in testis | 0.001 | 9.90E-01 | -0.03 | 5.45E-01 | -0.04 | 4.21E-01 | 0.065 | 2.06E-01 | 0.053 | 2.61E-01 |
| Regulatory density (ChIP-seq) | -0.15 | 2.37E-01 | 0.593 | 3.45E-06 | 0.742 | 3.06E-10 | 0.49 | 3.92E-05 | 0.327 | 7.64E-03 |
| Regulatory density in immune cells (ChIP-seq) | 0.047 | 6.24E-01 | -0.05 | 6.01E-01 | -0.08 | 3.94E-01 | -0.16 | 1.26E-01 | 0.146 | 1.32E-01 |
| Regulatory density in testis (ChIP-seq) | -0.8 | <1E-16 | -1.03 | <1E-16 | -0.86 | <1E-16 | -0.39 | 1.65E-09 | -0.11 | 4.76E-02 |
| Regulatory density (DNaseI) | -0.05 | 7.15E-01 | -0.61 | 5.45E-05 | -0.64 | 8.24E-06 | -0.84 | 1.66E-08 | -0.55 | 8.27E-05 |
| Gene length | -0.06 | 1.12E-01 | -0.05 | 2.84E-01 | -0.02 | 5.59E-01 | 0.004 | 9.13E-01 | 0.023 | 5.46E-01 |
| Gene number | -0.09 | 3.46E-01 | -0.52 | 1.28E-06 | -0.4 | 5.91E-05 | -0.46 | 6.23E-04 | -0.02 | 7.96E-01 |
| Coding density | 0.159 | 8.41E-02 | 0.535 | 8.72E-06 | 0.586 | 3.77E-08 | 0.396 | 1.93E-03 | 0.202 | 2.73E-02 |
| Number iHS data points | -0.01 | 8.19E-01 | 0.315 | <1E-16 | 0.33 | <1E-16 | 0.157 | 7.29E-08 | 0.081 | 1.51E-03 |

We also consider other functional factors like gene expression, including the average gene expression across 53 GTEx v7 tissues [40]. The expression in testis and immune cells could impact the frequency of sweeps, thus we also include them as independent expression variables [5,29]. We also consider the number of protein-protein interactions (PPIs) of each gene, given that this variable has been previously associated with the rate of sweeps [38]. Finally, we consider the influence of viruses on genomic adaptation, as previous evidence suggests that they have acted as a strong selective pressure during recent human evolution [9,24]. We use the distance of each gene to the closest gene coding for a protein that interacts with viruses. More details for all the factors considered are given in the Methods. Note that our analysis is not expected to detect all selective pressures affecting the human genome. We test factors that should associate with positive selection according to prior strong evidence, using them to test the ability of our new method to detect the genome-wide action of recent adaptation (see Discussion).

Genomic factors that are calculated across the genomic windows can vary depending on the window size used. Therefore, we infer the association between recent selection and factors measured in genomic windows of varying sizes (Methods). In total, we use five different window sizes (50, 100, 200, 500 and 1,000 kb) centered at the genomic center of Ensembl v99 coding genes [41]. For example, we measure the association between the average iHS within 50kb windows and the average recombination rate within the same 50kb windows, or between the average iHS and recombination within the same 1000kb windows (Methods). Using different window sizes also relaxes assumptions about the expected strength of selection, as larger windows are likely more sensitive to strong selection compared to smaller windows, and vice versa [9]. Using fixed size windows irrespective of gene length avoids a priori biasing measures of iHS by gene length. We also account for gene length in our model through the inclusion of multiple functional densities along with the genomic length of the gene (between transcription start and stop) at the center of the window as potential explanatory variables.

We estimate recent selection signals using the iHS statistic in five populations represented in the 1,000 Genomes Project [34]. Each population represents a different continent: Yoruba for Africa, Han Chinese for East Asia, Utah residents with Northern and Western European ancestry, Toscani for Southern Europe, and Peruvians for Americas. We select Peruvians given that they show the highest percentage of Native American ancestry among populations included in the 1,000 Genomes Project [34,42]. The iHS statistic measures how far the haplotypes carrying a specific derived allele extend upstream and downstream of this focal allele, compared to how far haplotypes carrying the ancestral allele extend [5]. As such, the approach provides one value of iHS for each biallelic SNP in the genome with clearly defined derived and ancestral alleles (Methods). Although isolated extreme values of iHS at a single SNP have often been used as a signature of selection, we recently found that measuring the average absolute value of iHS across all the SNPs in a genomic window provides high power to detect recent selection [9]. The average iHS in an entire window is also likely to better associate with other factors similarly measured as averages in the same windows. To account for the fact that the variance of the average iHS depends on the number of SNPs with individual iHS values in a window, we include this number as a predictor in the tested MDR models.

### MDR fit to recent selection signals

For each separate human population and window size, we model the association between genomic factors and iHS using a MDR with a mixture of two Gaussian distributions. We design one distribution to match more the drift component of iHS that is not expected to correlate with genomic factors. In contrast, the second distribution is designed to be sensitive to positive selection and it is expected to associate with the tested genomic factors (Methods). We apply this approach to account for the fact that only a part of the genome is expected to have undergone recent positive selection. In this context, a small second distribution and non-significant associations with the tested predictive factors would suggest a very small contribution of recent positive selection. The patterns of iHS would then be entirely dominated by genetic drift and past demography. We use this modeling approach together with an optimization algorithm to get the overall mixture distribution with the best fit for the observed iHS in each population and window size (Methods).

Overall, the MDR models fit well the distribution of observed iHS (Figure 1). iHS does not follow a Gaussian distribution, and our flexible approach combining a mixture of two Gaussian distributions and linear regression clearly fits better the iHS distribution than would a single distribution from a classic linear model. We find a clear separation between the Gaussian components, with the second component shifted toward higher iHS values, and thus likely enriched in positive selection (Figure 1). Together, the close fit and separation of the two Gaussian distributions likely provide an improved ability of our model to estimate the contributions of different factors to iHS.

### Patterns of selection across populations and window sizes

We find different patterns of selection across five populations from four different continents (Figure 1). The African Yoruba population shows the largest selection-enriched component (Figure 1A). This component is clearly shifted towards high iHS values, indicating stronger selection signals that stand out more from the first component. The selection-enriched components are less pronounced but still substantial in the four other populations (Figure 1B–1E). The Peruvian population shows the smallest selection-enriched component (Figure 1E). These results suggest that the visible (not necessarily the actual) contribution of selection to the whole variation of iHS is smaller in populations that were exposed to bottlenecks during migrations Out of Africa and consequently, more subject to genetic drift [34,35]. In other words, our results might be explained by the fact that selection is more visible in Yoruba due to a reduced effect of genetic drift.

With respect to window size, we find the largest selective components when using larger, 1,000 kb windows (Supplemental Results S2). This is an expected result, because it is easier to distinguish strong selection from genetic drift and background noise in larger windows. Smaller windows are less sensitive to strong selection, while larger windows can include a great accumulation of large iHS values. In addition, small windows are more influenced not only by the content of a genomic window, but also by genetically linked surrounding regions. These surrounding areas are in linkage with the genomic areas inside smaller windows as their center is closer to the edges. Therefore, smaller windows do not differentiate well between inside and outside genomic regions [1,9]. Given these results, we highlight primarily the results for the Yoruba population in the main text (while providing the results for all tested populations in main Tables and Figures), and we use 1,000 kb windows.

### Determinants of strong positive selection in human populations

Several genomic factors show significant associations with iHS in the selection-enriched component (Table 1). Unsurprisingly, we find the strongest association with recombination rate, which negatively correlates with iHS (e.g., Yoruba slope=-2.44; p-value<1E-16; Table 1). This is congruent with the eroding effect of recombination on selection signatures, impacting the probability to detect selective sweeps. Recombination can break haplotypes generated by selection, making it more difficult to detect the selective events [17,18]. These results confirm the role of recombination as a key determinant of the probability of detecting selection with iHS [8]. We also find an association between GC-content and recent positive selection. When the effect of recombination is not accounted for, that is, recombination is not included in the model, GC-content negatively correlates with iHS (Yoruba slope=-0.29; p-value=1.22E-03; Supplemental Results S3: Table S1). This is expected given that GC-content is positively associated with long-term recombination [43]. However, adding the control for recombination reveals a positive association between GC-content and iHS (Yoruba slope=0.55; p-value=1.34E-06; Table 1). Given that recombination already explains a great proportion of iHS variability including that shared with GC-content, the positive association between GC-content and iHS could be independent from recombination. Accounting for the eroding effect of recombination on haplotypes could make more visible a positive (direct or indirect) influence of GC-content on selection.

GC-content might be a better proxy of overall functional density than individual functional factors included in the regression model, which could explain its positive association with iHS. We test this hypothesis by removing different genomic factors from the model (see Supplemental Results S3 for modeling results after removing different sets of factors in Yoruba). The removal of GC-content makes emerge a positive association of iHS with DNAseI hypersensitivity, which is a measure of the overall density of transcription-factor binding sites (Yoruba slope=0.44 vs. −0.047; p-value=1.77E-07 vs. 7.15E-01; Table 1; Supplemental Results S3: Table S2). Since GC-content has been positively associated with the density of functional elements [44,45], removing GC-content may thus make more visible a positive association between iHS and functional density. We further test if this is also the case for more tissue-specific functional densities such as the density of regulatory elements in testis or immune cells that are expected to exhibit more positive selection (Methods), but we find no such evidence. The removal of GC-content does not affect to the association between selection and regulatory density in testis (slope= −0.765 vs. −0.795; p-value<1E-16 in both cases; Table 1; Supplemental Results S3: Table S2) and immune cells (slope=0.072 vs. 0.047; p-value=4.5E-01 vs. 6.23E-01; Table 1; Supplemental Results S3: Table S2). As shown later in this manuscript, fine-scale patterns of recombination seem to hinder the detection of associations for these regulatory variables. Finally, we find that the removal of coding density and DNAseI hypersensitivity increases the significance for the association between GC-content and iHS (p-value=2.40E-14 vs. 1.34E-06; Table 1; Supplemental Results S3: Table S3). From this model, we then further remove additional functional factors whose association with selection is also affected by the removal of GC-content, namely, gene number, gene length and ChIP-seq regulatory density (Methods). Accordingly, the removal of these functional factors leads to a highly significant association between GC-content and recent positive selection (slope=0.49; p-value<1E-16; Supplemental Results S3: Table S4). In summary, the removal of different functional genomic factors from the model supports that the positive association between positive selection and GC-content is mediated by the role of this factor as proxy of overall functional density. However, we cannot exclude the implication of other processes related to recombination that would require more detailed analyses (for example, see fine scale patterns related to regulatory density below).

Another genomic factor associated with selection is the density of conserved elements (Methods), showing a positive association with iHS (Yoruba slope=0.16; p-value=6.4E-04; Table 1). This is an expected result since coding sequences and non-coding regulatory elements tend to be conserved [37]. This factor shows a significant association in all populations except in the Peruvian population sample, which might reflect the impact of past bottlenecks on the visibility of the selective patterns (Table 1). The density of conserved elements is thus likely to be a good proxy of overall (coding and non-coding) functional density. The inclusion of this factor in the model could then, however, explain the lack of association for other factors related to functional density such as coding density. Indeed, the removal of conserved elements density makes significant the positive association between coding density and selection in Yoruba (Yoruba slope=0.26 vs. 0.16; p-value=2.35E-03 vs. 8.41E-02; Table 1; Supplemental Results S3: Table S5). All other populations have a clear significant positive correlation of iHS with coding density even without removing the density of conserved elements (Table 1). In addition, the simultaneous removal of conserved elements density and GC-content, which can also act as a proxy of functional density, increases even more the positive association of coding density with iHS in Yoruba (slope=0.29; p-value=7.76E-04; Supplemental Results S3: Table S6). These, and previous results about GC-content suggest that the lack of association for multiple functional factors could be explained, at least partially, by the simultaneous consideration of better measures of overall functional density in our models.

The number of protein-protein interactions is another factor that correlates with iHS. This factor associates negatively with selection, suggesting the existence of a slight depletion of selection for regions with higher number of PPIs (Yoruba slope=-0.07; p-value=4.25E-02; Table 1). This is opposite to that found by Luisi et al. [38], as they reported higher recent positive selection on central proteins within the human interactome. Higher number of PPIs has been also associated with higher positive selection in the chimpanzee lineage according to the McDonald-Kreitman (MK) test. However, this factor does not seems to be a determinant of positive selection when other genomic factors are simultaneously considered using a MK regression approach [46]. Given the inconsistency with previous evidence and the low strength of the association, the weak negative relationship between selection and the number of PPIs found in this study should be taken with caution.

The modeling approach presented in this study can also be used to test the influence of specific selective pressures. We illustrate this feature by showing that positive selection is associated with multiple variables related to viral interaction (Table 1). One of the factors more highly associated with iHS is the distance to VIPs in all tested populations (Yoruba slope=-0.17; p-value=4.16E-06; Table 1). Selection decreases further away from VIPs, in other words, we find an enrichment of selection around VIPs. Viruses have acted as a strong selective pressure during human evolution, shaping genomic adaptation, and previous studies have also found more positive selection at VIPs [9,24]. Gene expression is another functional factor significantly associated with selection (Table 1). The average expression across 53 tissues from GTEx [40] is negatively associated with iHS (Yoruba slope=-0.14; p-value=3.77E-02), while gene expression in immune cells shows a strong and positive association with selection in all populations (Yoruba slope=0.26, p-value=1.51E-05; Table 1). Given their relevance in the response to pathogens, genomic regions highly expressed in immune cells do represent expected targets of positive selection.

Overall, we find positive associations between the density of regulatory elements and iHS except in Yoruba where this association is not significant (Table 1; see below for analyses under neutral conditions explaining these results). However, the densities of regulatory elements in tissues expected to experience more positive selection (testis and immune cells) show surprising patterns of association with iHS (Table 1). In particular, a higher density of transcription factor binding sites in testis correlates with weaker iHS selection signals (slope=-0.79; p-value<1E-16). This is a counterintuitive result, given that more regulatory density would give more options to modulate gene expression and hence more room for positive selection to act. An explanation could be the following: hotspots of recombination are usually regions where the chromatin is more open, and hence more accessible for transcription factors to bind [47]. If the density of transcription factor binding sites in testis (where meiosis and recombination occur) is higher in areas with high recombination, then selection signals within these sites would be erased by recombination [17,18]. Fine scale patterns of recombination and selection support this hypothesis, as recombination tends to increase close to regulatory elements, while iHS tends to decrease (Supplemental Results S4: Figures S1-S3). Therefore, high levels of local recombination could erode haplotypes close to the selected variants, although the average recombination in the gene window covering that region is low. In other words, the 1,000 kb gene windows we use could have a low recombination rate on average, while containing local peaks of recombination close to regulatory elements. Our original model considers the average recombination in each 1,000 kb gene-centered window, so it is likely unable to detect the effect of these fine-scale patterns of recombination. We also observe a surprising lack of association between immune regulatory density and iHS (Table 1), even though we find a strong positive association between iHS and immune gene expression. Here again we find that the fine-scale patterns of recombination close to immune regulatory elements are likely to blame, with an increase in the fine-scale recombination rate as one gets closer to immune regulatory elements (Supplemental Results S4: Figure S3). It is surprising that we find a positive association between iHS and regulatory density across multiple tissues, but not when focusing on specific tissues where selection is expected to act (i.e., testis and immune cells). Although recombination rate tends to concentrate close to all regulatory elements, as it does more specifically around regulatory elements in testis and immune cells (Supplemental Results S4: Figure S1-S3), recombination declines rapidly when considering regulatory elements of all tissues compared to testis and immune cells (Supplemental Results S4: Figure S4). This suggests that we could have more power to detect positive associations with iHS and regulatory density in all tissues compared to testis or immune cells (see below for further evidence).

### Robustness of recent selection patterns to varying genomic window sizes in Yoruba

Next, we ask if the trends observed with 1,000 kb windows in the Yoruba genomes also hold or not when using smaller window sizes (Supplemental Results S2: Tables S1-S5). Some patterns visible in large 1000kb windows are also visible when using smaller genomic window sizes, while some are not (100 kb vs 1,000 kb windows; Supplemental Results S2: Tables S2, S5). The strong negative association between recombination rate and selection is found regardless of window size. On the contrary, the distance to VIPs for example shows a strong association with iHS in large windows but not in smaller windows (100 kb windows: slope=-0.02; p-value=5.11E-01; Supplemental Results S2: Tables S2, S5). This is consistent with previous evidence showing that VIPs are particularly enriched in strong selective sweeps, for which larger windows are more sensitive [9]. This is also the case for the density of conserved elements (100 kb windows: slope=0.1; p-value=9.3E-02) and especially regulatory elements in testis, with a slope going from −0.79 in 1,000 kb windows to −0.03 in 100 kb windows (Supplemental Results S2: Tables S2, S5). These results suggest that both factors may correlate with strong rather than weak selection. In the case of testis regulatory density, this pattern might be explained by the fact that smaller windows take better into account the fine scale patterns of recombination, which limits the negative influence of using average recombination (see below for fine-scale analyses of recombination). Note, however, that all these differences might also be explained by the fact that smaller windows do not differentiate well between inside and outside of the genomic regions they are supposed to delineate. That is, they may be more influenced by the surrounding, genetically linked genomic regions [1,9]. See Supplemental Results S2 for the complete results across populations and window sizes.

### Population simulations of neutral expectations of associations with iHS

As described above, we find multiple strong functional associations with iHS in the directions expected under recent positive selection. That said, some observations remain unexplained at this point, with for example a lack of positive association between overall regulatory density and iHS in Yoruba. We also observe a lack of association between the overall sum of regulatory plus coding functional density and iHS in Yoruba (slope=-0.063; p-value=0.65; Supplemental Results S5: Figure S1). The association of this overall coding plus regulatory density with iHS is strongly positive in the other tested populations (Supplemental Results S5: Figure S1). This prompted us to better characterize the null expectations of the associations between this overall coding plus regulatory density (as a measure encompassing the different functional types of elements considered in our model) and iHS in the absence of positive selection. In particular, the heterogenous, fine scale patterns of recombination and their variation around functional elements (Supplemental Results S4: Figures S1-S5) might make null neutral expectations deviate from a simple lack of association with slopes centered around zero.

We therefore use forward SLiM [48] neutral simulations to estimate the neutral expected associations between the overall coding plus regulatory density and iHS in our analysis (Methods). Under neutral conditions (i.e., only neutral mutations), and recreating the actual distribution of recombination rate and functional (coding plus regulatory) elements (Methods), we find that the association between iHS and functional density tends to be negative, and in every case much less positive compared to the studied populations including Yoruba (Supplemental Results S5, Figure S1). In addition, the magnitude of the second Gaussian component (estimated with *p*, see Methods) is also lower under neutral expectations compared to Yoruba (Supplemental Results S5: Figure S2). This suggests that the results observed with the MDR approach cannot be explained by neutral evolution. Moreover, these results support that, although the second Gaussian distribution captures neutral loci as shown by its presence even under strictly neutral evolution (Supplemental Results S5: Figure S2), this component is also sensitive to adaptation, supporting its enrichment in recent positive selection. These results also show that the lack of association between iHS and functional coding plus regulatory density in

Yoruba does not represent a lack of support for positive selection. Indeed, the null expected distribution for this association is negative and the observed Yoruba association is clearly above the neutral distribution (Supplemental Results S5: Figure S1). In the discussion we mention multiple reasons why recent positive selection may have been not less, or even more common in the Yoruba, while still having a weaker impact on iHS.

### Fine scale patterns of recombination around regulatory elements

The increase of recombination rate around regulatory elements may explain the fact that the null expectations for the association between functional density and iHS is negative (Supplemental Results S4: Figures S1-S5). Our neutral simulations recreate the distribution of recombination rate and functional (coding plus regulatory) elements of the human genome, thus regulatory and coding elements tend to be close to recombination peaks also in the simulations (Supplemental Results S4: Figure S5), leading to lower iHS and lowering the null expectation. This led us to further analyze the influence of local patterns of recombination around regulatory elements. Given the local increase of recombination around regulatory elements, it is likely that whole window estimates of recombination are not a sufficient measure of its effect on the association between iHS and regulatory densities. In particular, this might explain the surprising negative association between iHS and testis regulatory density or the lack of association for immune regulatory density (Table 1). Indeed, recombination rate tends to be higher specifically around these regulatory elements compared to the whole set of regulatory elements (Supplemental Results S4: Figure S4). We use three variables related to the recombination around regulatory elements in immune cells, testis and across multiple tissues, respectively. We calculate the more local, average recombination around regulatory elements within each gene window, up to a maximum distance of 5kb from each side of regulatory elements. Each variable is included in the original model of Yoruba 1000 kb separately, and replaces the previous, window-wide average recombination. Given that some gene windows can overlap with more regulatory elements than others, we also consider the number of recombination data points obtained for regulatory elements inside each gene window. In this way, we account for the fact that recombination may vary more broadly in windows that have less regulatory elements. See Supplemental Results S4 for further details about these calculations.

The iHS statistic decreases around regulatory elements in immune cells while recombination increases, staying very high further from immune regulatory elements compared to other tissues (Supplemental Results S4: Figures S1-S4). This might have confounded the expected positive association with selection in the original regression model with window-wide recombination rates. The control for local patterns of recombination around immune regulatory elements should however increase our power to detect this association. In line with this prediction, we find that the association between immune regulatory density and iHS becomes positive and highly significant when considering the local patterns of recombination around immune regulatory elements (Yoruba slope=0.68 vs. 0.047, p-value=4.60E-14 vs. 6.24E-01; Table 1; Supplemental Results S4: Table S1). In order to assess whether this increase is caused just by the lack of consideration of recombination patterns at a window-wide scale, we repeat this analysis but including the original window-wide average recombination variable. After including the average recombination rate at window-wide scale, the positive association remains stronger compared to the original model (slope=0.28 vs. 0.047, p-value=5.72E-03 vs. 6.24E-01; Table 1; Supplemental Results S4: Table S2).

As explained above, we hypothesize that the confounding effect of recombination is caused by a mismatch between different scales. Gene windows with low recombination can have local peaks of recombination around transcription factor binding sites (Supplemental Results S4: Figures S1-S4), as these peaks are more accessible to transcription factors [47]. Therefore, recombination could erode signals of positive selection around regulatory elements even if the region has an overall low recombination. In contrast, regions with high recombination at the average window-wide level should suffer less from this confounding effect, as recombination rate is then high both at a local and window-wide scale. Therefore, focusing on high-recombination regions should improve our ability to detect the expected positive association between selection and regulatory density. This approach, however, has the caveat of reducing the power to detect positive selection due the higher probability that recombination breaks selected haplotypes. We nevertheless repeat the analysis focusing only on gene windows with a recombination rate equal or higher than the second tertile (1.552 cM/Mb). Focusing on high-recombination regions, regulatory density in immune cells shows a stronger association with selection in a model including recombination rate around both gene windows and immune regulatory elements compared to the same model run across all genes (Yoruba slope=0.85 vs, 0.28, p-value=1.27E-08 vs. 5.72E-03; Supplemental Results S4: Tables S2-S3). Interestingly, regulatory density in testis also shows a positive association with selection in a model including recombination around both testis regulatory elements and gene windows and focused on high-recombination regions. Note that this regulatory variable shows one of the strongest negative associations with selection in the original model (Yoruba slope=0.58 vs. −0.79, p-value=4.98E-05 vs. <1E-16; Supplemental Results S4: Tables S4-S6), an unexpected result given the role of regulatory sequences in positive selection [1]. These results suggest that, indeed, a higher regulatory density can increase the probability for positive selection to occur, especially in some tissues. However, this signal is distorted by fine-scale patterns of recombination, which have to be carefully taken into account.

## Discussion

We find several functional factors independently associated with recent positive selection. For instance, viral interactions along with gene expression and regulatory density in immune cells are strongly and positively associated with selection. Viruses have acted as key drivers of adaptation in humans and VIPs show higher expression in lymphocytes compared to non-VIPs [26,28,49]. These results confirm the existence of strong viral and immune selective pressures during recent human evolution [9,24,28]. We also detect a positive association between testis regulatory density and selection, which is congruent with the fact that reproduction-related functions show signals of positive selection [5,29]. Note, however, that this latter result has been more difficult to reveal in our analyses, as it is only visible after controlling for fine-scale patterns of recombination around testis regulatory elements and after focusing on high-recombination regions, thus it is more questionable. Finally, the density of conserved elements and GC-content, which are related to overall (coding and non-coding) functional density [37,44,45], also show a positive association with selection. In summary, recent and positive selection is associated with the distribution of functional regions across the human genome.

If sweep signals are false positives only reflecting genetic drift, the signatures of sweeps should occur randomly across the genome. In contrast, if selective sweeps were relatively common during recent human evolution, we expect that other factors on top of recombination should associate with sweep signals across the human genome. In that case, we expect that selective sweeps should occur more frequently around functional (coding or non-coding) elements where adaptive mutations are expected to take place [1], also around genomic regions associated with specific functions. We find strong evidence for the latter scenario, as multiple functional factors are associated with recent positive selection. In other words, the distribution of sweep signals is not random relative to the distribution of functional elements and is associated with the functional characteristics of the human genome. That said, stochasticity still plays a role in the occurrence of advantageous mutations in the first place, and we still expect randomness when considering which specific loci of the human genome with equivalent functional characteristics have experienced positive selection. Our results thus support that selective sweeps were at least common enough during recent human evolution to generate associations between selection and functional factors.

Our MDR approach shows a good fit to the distribution of iHS across the human genome, with its two components showing a clear differentiation and the second being enriched in positive selection. This suggests that the consideration of a more biologically meaningful model, which assumes two Gaussian distributions instead of one, works better to analyze recent adaptation than simple correlations. Given that the MDR approach represents more directly genome evolution, the associations detected by this approach are more informative than classic linear models and partial correlations. The associations detected by these classical approaches may be caused by a deficit of weak iHS values, not by an enrichment of high iHS values as shown in the second component of our MDR approach, a scenario more specifically expected under strong selection. Indeed, we cannot replicate our results using classic linear models and partial correlations in 1,000 kb windows. These classic approaches show much lower correlations between iHS and functional factors expected to associate with positive selection (Supplemental Results S1: Tables S1-S2). For example, they are unable to replicate the strong enrichment in positive selection around VIPs reported by previous studies [9,24]. This is also expected given the poor fit of a single Gaussian distribution to the whole iHS distribution compared to our approach assuming two Gaussian distributions (Supplemental Results S1: Figure S1). Therefore, our results support the consideration of two distributions to model positive selection in the human genome. The characteristics of our MDR approach make it possible to reveal expected, but also unexpected patterns. For instance, GC-content is strongly and positively associated with iHS. Regions with high GC-content also tend to have high long-term recombination rates [43], which would predict a negative association between iHS and GC content, not positive. A possible explanation for this pattern is that the MDR approach can control for recombination as a confounder by simultaneously analyzing recombination rate and other genomic factors. Therefore, it can control for the impact of recombination on the probability to detect selection. This can unravel new patterns, like a positive association between GC-content and sweep signals. This association only emerged after adding recombination in the model, becoming even more significant after the removal of functional factors like coding density, gene number or regulatory density. This supports the independence of this association from recombination and the role of GC-content as a better proxy of overall functional density than individual factors. These results illustrate the utility of our approach. The simultaneous consideration of multiple predictors lets us unravel novel associations between genomic factors and adaptation. In addition, it enables easier testing of different hypotheses by modifying the set of predictors considered. Therefore, the MDR approach is useful to model recent genomic adaptation, and to analyze its prevalence and determinants.

We find other unexpected results using the MDR approach. For instance, we do not find a positive association between selection and testis or immune regulatory density in the original model. This is an unexpected result given that a higher regulatory density would lead to a larger mutational target for adaptation through changes in expression and these tissues are expected targets of positive selection [5,29]. As previously posited, the adaptive signals associated with regulatory density could be eroded due to the higher tendency of transcription factors to bind DNA at recombination hotspots [47]. This is supported by the higher recombination and lower iHS found closer to regulatory elements, along with the fact that the positive associations for immune and testis regulatory density only emerge when considering the fine-scale patterns of recombination. These results support the role of regulatory sequences as targets of positive selection independently of other genomic features [1], along with the relevance of testis and specially immune cells in recent human evolution [5,29]. Our results also illustrate how the detection of selection can be hindered by fine-scale characteristics of the genome, like the sharp increase of recombination we observe around regulatory elements. This might mask selection signals around regulatory elements, but not only because of the breaking of haplotypes caused by recombination. An additional mechanism might be the existence of biased gene conversion in these recombinant regions, which could specifically favor GC mutations (gBGC) [43,50,51]. This could lead to an increase in the frequency of GC mutations that are associated with short haplotypes (due to the eroding effect of recombination). BGC might therefore make it even more difficult to detect recent positive selection around regulatory regions, where it is expected to happen [1]. Note, however, that the present study does not provide evidence about the implication of gBGC in the observed patterns. It can be difficult to disentangle the eroding effect of recombination from GC-biased gene conversion on haplotypes because gBGC is caused by and therefore very correlated with recombination. Distinguishing the specific effect of biased-gene conversion may require precise recombination maps for African populations, where we find the largest selection-enriched second Gaussian distribution. Although we have not found evidence of the implication of gBGC yet, the fact that this hypothesis has emerged from implementing our model illustrates its utility to propose new hypotheses about the different processes that influence the detection and occurrence of positive selection.

Although the MDR approach provides an excellent fit to the observed selection statistic, we may still not detect associations with specific factors because they do not have a simple linear relationship with recent selection. Although more biologically meaningful than a model assuming just one distribution, our approach is still linear, thus more complex relationships may not be detected. For example, the weaker association of iHS with coding density in Yoruba could be explained not only by the consideration of better proxies of overall functional density, but also by the existence of non-linear relationships with selection. Advantageous mutations are more likely to appear in regions with high coding density, thus a high density of coding sequences should favor the appearance of selection [1]. However, high coding density can also mean a higher frequency of deleterious mutations. These mutations can interfere with adaptation when they are in linkage disequilibrium with advantageous mutations. This would be especially relevant for genomic regions with low recombination, where the linkage disequilibrium between deleterious and advantageous mutations is more likely [19,52]. Therefore, a higher coding density could favor or hinder positive selection depending on the circumstances, complicating the detection of an association between coding density and selection with our approach. The existence of this genetic interference could also explain the different results observed across populations in relation to coding density. Despite having the largest selection-enriched component, the Yoruba population shows a weaker association between selection and coding density compared to other populations. This association becomes visible only after removing proxies of overall functional density (i.e., density of conserved elements and GC-content). A potential, but still speculative explanation at this point might be the fact that the bottleneck related to the migration out of Africa may have removed segregating recessive deleterious variants, which may reduce their interference over advantageous mutations, thus making it easier for positive selection to act on regions with a high coding density [52]. In contrast, there was no similar bottleneck to decrease the genetic interference in the Yoruba, and regions of the genome with high a coding density may still experience stronger interference from recessive deleterious mutations. Interestingly, despite this potential limitation of positive selection, we still see a larger selection-enriched component in the entire Yoruba genome, suggesting that selection might still be more abundant in this population. There are other possible explanations for this apparent contradiction. For example, the ability of iHS to detect selective variants could be more limited in African populations due to lower overall linkage disequilibrium, together with the larger genetic diversity in these populations, which could favor the existence of sweeps from standing genetic variants that are harder to detect with iHS [30]. These potential explanations are speculative at this point and will require further investigation. Our results nevertheless demonstrate that the complex and heterogenous distributions of multiple genomic factors, together with summary statistics of hitchhiking robust to background selection, need to be used in order to quantify the determinants and importance of recent positive selection in the human genome.

## Methods

### Window sizes and genes coordinates

All analyses were performed using hg19 genomic coordinates and protein-coding gene annotations from Ensembl v99 [41]. We selected hg19 instead of GRCh38 mainly because this was the assembly used by the 1,000 Genomes consortium to generate and phase the genetic variants considered in our analysis. Note that the assembly selection should not influence our results as we performed genome-wide scale analyses, thus avoiding focusing on specific regions where the assembly could change after hg19. We considered windows of different sizes (50, 100, 200, 500 and 1,000 kb). Each window was centered at the genomic center of a gene, in the middle between the most upstream transcription start site and the most downstream transcription end site. We used a fixed window size to avoid biases related to gene length, as larger genes are more likely to overlap with high local iHS values just by chance compared to shorter genes. This would bias power in favour of larger genes, increasing the probability to detect sweeps in them just because of their length. In addition, the consideration of different window sizes provides the opportunity to detect different types of sweeps as larger windows are specifically sensitive to strong selective sweeps compared to smaller windows [9].

### Genomic features

Multiple genomic factors were calculated inside the gene windows. We considered the following factors that are likely to influence the frequency of sweeps:

- Length of the gene at the center of each window.
- Number of genes overlapped with each window. We used this value as an estimate of gene density.
- Recombination rate. For each window, we calculated the genetic distance between the edges of the window and then divided it by the physical distance between them. Genetic position of each window edge was obtained from the deCODE 2019 genetic map [53]. In case no data was available for a window edge, we selected the closest genetic position data points within 50 kb at each side of the window edge to estimate its genetic position. We used linear interpolation to this end, considering the genetic and physical position of the two points around the window edge. In other words, we assumed that genetic distance increased linearly between the two selected data points, and hence we can estimate the genetic position of the window edge based on its physical position respect to the other two positions. If no genetic position data was available at 50 kb or closer to the window edge, no genetic position was calculated and the whole window was discarded for recombination calculations and subsequent analyses. Note that we searched for genetic position data points only up to 50 kb from each window edge to avoid that the total distance between points at each edge could be too large (especially for larger windows). In that scenario, linear interpolation could be inappropriate because we would use very distant data to calculate recombination rate. For example, if a 1,000 kb window had no genetic position for any of its edges and we looked for genetic positions up to 1,000 kb at each side, it would possible that the selected data points of genetic position downstream and upstream of the window were separated by 3,000 kb. In addition, the recombination rate would not correspond with the size of the window.
- Density of coding sequences. We used Ensembl v99 coding sequences. The density was calculated as the proportion of coding bases respect to the whole length of the window without considering gaps (gap locations obtained from the UCSC Genome Browser; https://genome.ucsc.edu/; http://hgdownload.cse.ucsc.edu/goldenPath/hg19/database/). A similar approach was used for the rest of density estimates.
- Density of mammalian phastCons conserved elements [37], also downloaded from the UCSC Genome Browser. Given that each conserved segment had a score, we considered as conserved only those segments above a given threshold. In order to minimize the inclusion of non-conserved elements, we used a threshold that only considers 4.17% of the genome as conserved, as it is unlikely that much more than that is strongly constrained [37].
- Density of regulatory elements. We calculated several variables related to the density of binding sites for transcription factors. In all cases, we calculated the proportion of sequences that are considered binding sites within a window. The data were also obtained from the UCSC Genome Browser [39].

- Density of DNaseI hypersensitive sites (wgEncodeRegDnaseClusteredV3 track). We considered as binding sites those segments with a score higher than a given threshold. The selected threshold was a score value according to which only 10% of the genome is considered accessible to DNaseI, and hence a binding site for transcription factors. In this way, we minimize the probability to consider non-binding sites (i.e., false positives).
- Density of binding sites according to the technique of chromatin immunoprecipitation followed by sequencing (ChIP-seq; encRegTfbsClustered track). We calculated a threshold to consider a given segment as a binding site following the same approach used for DNaseI hypersensitive sites. This database includes 1264 experiments representing 338 transcription factors in 130 cell types. We calculated the density of binding sites using the whole dataset, but also considering two subsets, one for experiments performed on cells lines of testis, and another for experiments performed on immune cells (lymphocytes in most cases). Note that the threshold was calculated considering the 1264 experiments, being then applied to the whole dataset and the two subsets.
- GC-content, calculated as a percentage per window. It was also obtained from the UCSC Genome Browser.
- Gene expression: We used the log (base 2) of TPM (Transcripts Per Million), which was obtained from GTEx [40] (https://www.gtexportal.org/home/). We considered the average gene expression across 53 GTEx v7 tissues, but also expression in immune cells (lymphocytes) and testis.
- Number of protein-protein interactions (PPIs) in the human protein interaction network [38]. We used the log (base 2) of the number of PPIs of each gene. We summed 1 to the original dataset to avoid problems applying the logarithm to genes with no interactions (i.e., PPI=0).
- Proportion of genes that interact with viruses (VIPs). We used a previously published dataset [9] that includes around 4500 VIPs with evidence of physical interaction with viruses. 1920 of these VIPs were manually curated from the virology literature, while the remaining 2600 VIPs were identified using high-throughput methods and retrieved from the VirHostnet 2.0 database and additional studies. See the original publication for further details [9]. The variable included in the model was calculated as the distance of each gene to the closest gene that interacts with viruses.

### iHS calculation

In order to calculate iHS, we used polymorphism data from the 1,000 Genomes Project phase 3 [34]. We calculated iHS for 5 populations: Yoruba, Utah residents with Northern and Western European ancestry, Toscani, Han Chinese and Peruvians. We used the hapbin software [33] in order to perform fast scans of iHS across the genomes of these populations. Note that the genetic maps used as input in hapbin were calculated using the deCODE 2019 genetic map [53]. In case that no data was available for the focal variant, we again used linear interpolation to estimate its genetic position. We followed the same approach used to calculate the genetic position of window edges during recombination calculations. In this case, we searched for genetic position data points at both sides of the focal variant until 1,000 kb. We discarded variants with no data points around, as they were present in genomic regions with low density of recombination data. The genetic maps based on the deCODE 2019 map were used as an input in order to calculate the iHS for the 5 populations.

The raw iHS values obtained with hapbin were standardized by frequency. For each population, we divided variants in 50 bins of frequency (from 0 to 1) considering the whole genome. We then calculated the mean and standard deviation of raw iHS within each bin, using them to standardize raw iHS values: (raw iHS - mean iHS) / sd iHS. The mean of standardized iHS was calculated per window, considering the absolute value of iHS. In our case, large positive iHS values can be caused by unusually long haplotypes carrying the derived or the ancestral allele, in the case the ancestral alleles hitchhiked together with the actual, selected derived allele. Therefore, high iHS can be interpreted as selection in favor of the derived or ancestral haplotype. We also obtained the number of iHS values per window to have an estimate of data density. Mean iHS in windows with low number of iHS values can be more influenced by outliers (very low or very high iHS values). Therefore, we included the number of iHS values per window as predictor in the models to control for this.

### Modeling

We modeled the association between genomic factors and iHS in each population and window size by combining a Gaussian mixture distribution with linear regression. Mixture distributions are useful to model data that cannot be fully described by a single distribution and that are likely generated by multiple processes. In this case, a mixture of two Gaussian distributions could fit better a scenario where selection influences some genomic regions but not others, and hence a bimodal distribution of iHS would be observed across the genome. Therefore, we posit that sweep signals in the human genome may have components differentially influenced by positive selection. According to this, we assumed that observed log iHS followed a mixture of two Gaussian distributions representing two components, one more influenced by drift and a second one enriched in positive selection. Because selective sweeps often result in elevated iHS, we assumed that the Gaussian distribution for the selection-enriched component had a higher mean than that for the first component. Furthermore, to model the association of genomic factors with the selection-enriched component of iHS, we assumed that the probability of the selection-enriched component was a linear combination of genomic factors followed by a sigmoid transformation. Specifically, for each factor, we calculated the product of its observed value with a slope indicating its association with the selection-enriched component of iHS. A positive slope indicates that the probability of the selection-enriched component increases with the increasing value of a factor, whereas a negative slope indicates the probability decreases with the factor. Then, we summed the product over all factors for a given gene to model the accumulative effect of genomic factors on the probability of the selection-enriched component. Then, we added an intercept indicating the baseline level of selection when all genomic factors are equal to 0 and transformed the result to a probability (0 to 1) using a sigmoid function. This probability was obtained for each gene and can be regarded as the overall influence of all genomic factors on the frequency of recent selective sweeps. These steps are described by the equation 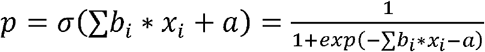 where *p* is a probability summarizing the influence of all studied genomic factors on the selection-enriched component of iHS, *σ* is the sigmoid function, *b_i_*, and *x_i_*, are the slope and the observed value for genomic factor *i* in a given gene, respectively, and *a* is the intercept.

To obtain the likelihood of log iHS in each gene, we calculated a weighted sum of Gaussian density functions, *P*(*Y*) – *p*\* *f*(*Y*|*μ*_1_ *σ*_1_) + (1 – *p*) * *f*(*Y*|*μ*_0_, *σ*_0_), where *Y* and *P*(*Y*) are the observed log iHS and its likelihood in a given gene, respectively, *p* is the probability of the selection-enriched component calculated in the previous step, *f*(*Y*|*μ*_1_, *σ*_1_) is the Gaussian density function with mean *μ_1_* and standard deviation *σ_1_* describing the selection-enriched component, and *f*(*y*|*μ*_0_ *σ*_0_) is the Gaussian density function with mean *μ_0_* and standard deviation *σ_0_* describing the first component. Finally, we summed the logarithm of *P*(*Y*) over all genes to obtain the log likelihood of the entire dataset.

We estimated model parameters by maximizing the log likelihood of the entire dataset (L-BFGS-B optimization). These parameters included the mean and standard deviation of each Gaussian distribution, along with the intercept and the slopes representing the association between the selection-enriched component of iHS and each genomic factor. For each genomic factor, we tested the significance of its slope by comparing the log likelihood between the full model with all genomic factors and a nested null model without the selected factor. We assumed that two times the difference in log likelihood between the two models followed a chi-square distribution with one degree of freedom. The script and input data needed to run the MDR are publicly available at https://github.com/dtortosa/Mixture_Density_Regression_pipeline.

### Population simulations to recreate the neutral expectation

We used SLiM [48] to simulate the neutral expectations in our analyses and assess in that way if our results could be fully explained by neutral patterns. First, we randomly selected 100 genomic segments of 20 mb each one, totaling 2000 mb. We recreated the functional (coding and regulatory) elements observed in the human genome using the same coordinates considered in the calculation of functional densities for the main analyses (see Genomic features section). We also considered the same recombination map (deCODE 2019) to recreate the actual recombination heterogeneity in these segments. Therefore, we simulated a total of 2000 mb using the observed distribution of functional elements and recombination rate. We only simulated neutral mutations in all the simulated genomic regions. The population size was set to 10,000 individuals. We first run a burn-in period of 100,000 generations, running then 100 independent neutral simulations with 2,000 additional generations. For each independent neutral simulation, we calculated iHS using the same approach than in the actual genomes. As the simulated segments had direct correspondence with the human genome, we extracted the genes included in each segment, and calculate 1000 kb windows centered around them. Inside each window, we calculated the average of iHS and the number of iHS data points, along with recombination rate and functional density as done in the actual genomes. Finally, we used the MDR to model iHS as a function of recombination rate, functional density and the number of iHS data points measured all in 1000 kb windows. Therefore, we model iHS under neutral conditions across 100 independent runs simulating each one 2000 mb. In addition to the genomic factors already mentioned, we also included the GC-content observed for each gene window in the actual genome. It can be useful to simulate GC to control for varying mutations rates, however we used a uniform mutation rate across the simulated regions, so we did not include it in the simulations. We are still adding it as a predictor using the GC-content in the actual gene windows because this factor is associated with both functional density and long-term recombination [43–45]. This can help to make visible more local patterns of recombination. In addition, it makes fairer the comparison between the neutral simulations and the observed genomes, as in the latter iHS was also modeled using GC-content.

From the model of each neutral run, we obtain the slope for the association between functional density and iHS, while controlling for the rest of genomic factors calculated. We also took *p*, i.e., the accumulative effect of all genomic factors on the probability of the second component of iHS (see Modeling section). We calculated its average across all genes within each neutral run, using it as a measure of the magnitude of the second component of iHS. We compared the functional associations and the magnitude of the second Gaussian component between the neutral expectations and the actual genomes. Note that in the latter comparison, we only considered Yoruba. As previously noted, the distribution of iHS seems to be more influenced by drift in non-African populations due to their greater exposition to bottlenecks. This explains the reduction of the second Gaussian component in these populations. Indeed, the population that suffered the most extreme bottlenecks is the one showing the smallest second component, i.e., Peruvians. Our simulations reproduce a constant population size of 10,000 individuals and no bottlenecks, thus it is fairer to compare the magnitude of the second Gaussian component in the simulations with the African Yoruba population, which was not exposed to bottlenecks in the same degree than the rest of studied populations.

## Supporting information

Supplemental Results S1

Supplemental Results S2

Supplemental Results S3

Supplemental Results S4

Supplemental Results S5

## Acknowledgments

We thank the members of the Enard, Masel and Gutenkunst labs for their helpful comments. DFST was supported by a Marie S. Curie Global Fellowship within the European Union research and innovation framework programme (2014-2020; ClimAHealth; https://doi.org/10.3030/101030971). DE was supported by the National Institute of General Medical Sciences of the National Institutes of Health under award number R35GM142677 (https://www.nigms.nih.gov/). Y-FH was supported by the National Institute of General Medical Sciences of the National Institutes of Health under award number R35GM142560 and by startup funds from Pennsylvania State University (https://www.nigms.nih.gov/; https://www.psu.edu/). The content is solely the responsibility of the authors and does not necessarily represent the official views of the National Institutes of Health or the European Union. The funders had no role in study design, data collection and analysis, decision to publish, or preparation of the manuscript.

## Author contributions

Y-FH, DE and DFST designed the study. DE and DFST collected the data. Y-FH and DFST performed the analyses. DE and DFST wrote the paper and interpreted the results. Y-FH, DE and DFST edited and approved the paper.

## Supporting information captions

Supplemental Results S1. Results obtained using classic linear models and partial correlations.

Supplemental Results S2. Associations between genomic factors and iHS across populations and window sizes.

Supplemental Results S3. Associations between genomic factors and iHS after removing specific predictors.

Supplemental Results S4. Fine scale analyses of recombination rate, regulatory density and positive selection.

Supplemental Results S5. Results of population simulations.

