## Supplemental Results S1 for "Mixture Density Regression reveals frequent recent adaptation in the human genome"

### Classical partial correlations

We have performed classic partial correlations (Spearman's rank correlations) between the whole iHS distribution and individual genomic factors while controlling for the rest of factors. See results for some functional factors expected to be associated with positive selection.

| Covariate | Yoruba | | Toscani | | Utah residents | | Han Chinese | | Peruvians | |
|---|---|---|---|---|---|---|---|---|---|---|
| | ρ | p-value | ρ | p-value | ρ | p-value | ρ | p-value | ρ | p-value |
| Density of conserved elements | -0.033 | 3.88E-05 | 0.053 | 5.16E-11 | 0.046 | 8.93E-09 | 0.039 | 1.48E-06 | 0.056 | 2.63E-12 |
| Distance to VIPs | -0.012 | 1.46E-01 | -0.007 | 4.07E-01 | -0.002 | 7.69E-01 | 0.005 | 4.98E-01 | -0.027 | 8.47E-04 |
| Gene expression in immune cells | 0.026 | 9.57E-04 | 0.042 | 1.79E-07 | 0.040 | 5.58E-07 | 0.032 | 5.44E-05 | 0.022 | 5.28E-03 |
| Regulatory density (ChIP-seq) | -0.029 | 2.77E-04 | 0.036 | 7.21E-06 | 0.007 | 3.60E-01 | -0.005 | 5.70E-01 | 0.002 | 8.50E-01 |
| Coding density | 0.036 | 6.67E-06 | 0.045 | 2.13E-08 | 0.050 | 5.57E-10 | 0.052 | 5.49E-11 | 0.041 | 2.55E-07 |

**Table S1: Correlation coefficients (Spearman's ρ) and p-values of the association between iHS and genomic factors for the five studied populations in 1,000 kb windows: Yoruba, Toscani, Utah residents, Han Chinese and Peruvians.**

### Classical linear models

We have run classic linear models considering the whole iHS distribution as response and genomic factors as predictors. See results for some functional factors expected to be associated with recent positive selection.

| Covariate | Yoruba | | Toscani | | Utah residents | | Han Chinese | | Peruvians | |
|---|---|---|---|---|---|---|---|---|---|---|
| | slope | p-value | slope | p-value | slope | p-value | slope | p-value | slope | p-value |
| Density of conserved elements | 0.005 | 6.25E-01 | 0.085 | 2.28E-16 | 0.086 | 5.89E-17 | 0.085 | 6.61E-16 | 0.062 | 4.61E-09 |
| Distance to VIPs | -0.024 | 2.87E-03 | -0.042 | 3.12E-07 | -0.042 | 2.53E-07 | -0.028 | 9.80E-04 | -0.034 | 5.82E-05 |
| Gene expression in immune cells | 0.062 | 1.13E-05 | 0.090 | 1.22E-10 | 0.097 | 3.86E-12 | 0.073 | 3.51E-07 | 0.060 | 3.19E-05 |
| Regulatory density (ChIP-seq) | -0.036 | 1.10E-01 | 0.136 | 9.71E-10 | 0.088 | 7.88E-05 | 0.036 | 1.13E-01 | -0.043 | 5.98E-02 |
| Coding density | 0.037 | 6.68E-02 | 0.092 | 4.89E-06 | 0.089 | 8.49E-06 | 0.104 | 4.16E-07 | 0.068 | 1.02E-03 |

**Table S2: Slopes and p-values of the association between iHS and genomic factors for the five studied populations in 1,000 kb windows: Yoruba, Toscani, Utah residents, Han Chinese and Peruvians.**

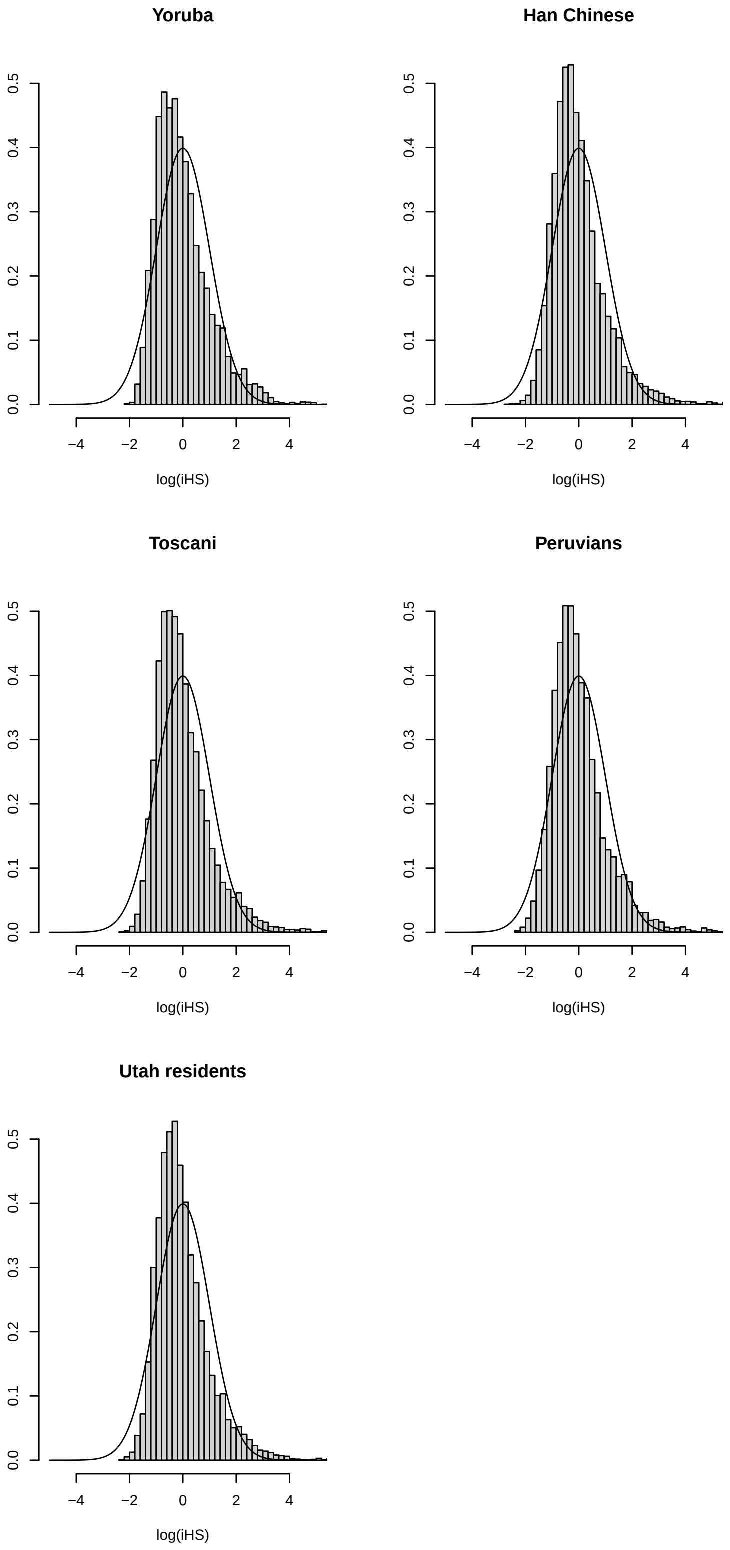
Yoruba
Han Chinese
Toscani
Peruvians
Utah residents
0.0
0.1
0.2
0.3
0.4
0.5
−4
−2
0
2
4
log(iHS)

**Figure S1: Single gaussian distribution fitting observed iHS (1,000 kb windows) for the five studied populations:** A) Africa - Yoruba; B) East Asia - Han Chinese; C) Europe – Toscani; D) America – Peruvians; E) Europe - Utah residents with Northern and Western European ancestry. For each population, the figure shows an histogram of the observed iHS (1,000 kb). It also shows a single Gaussian distribution (black solid line) with the mean and standard deviation of the observed iHS.
