## Supplemental Results S2 for "Mixture Density Regression reveals frequent recent adaptation in the human genome"

### Yoruba

#### *Yoruba 50kb*

Figure S1: Mixture of Gaussian distributions fitting observed iHS (50kb windows) for Yoruba. The figure shows the two Gaussian distributions, component 1 and 2, being the latter enriched in positive selection. In that component, iHS linearly depends on the genomic factors considered. Legend: Light blue = Observed iHS; Dark blue = Mixture model; Full red curve = Component 1 of the mixture model; Dashed red curve = Component 2 of the mixture model enriched in selection.

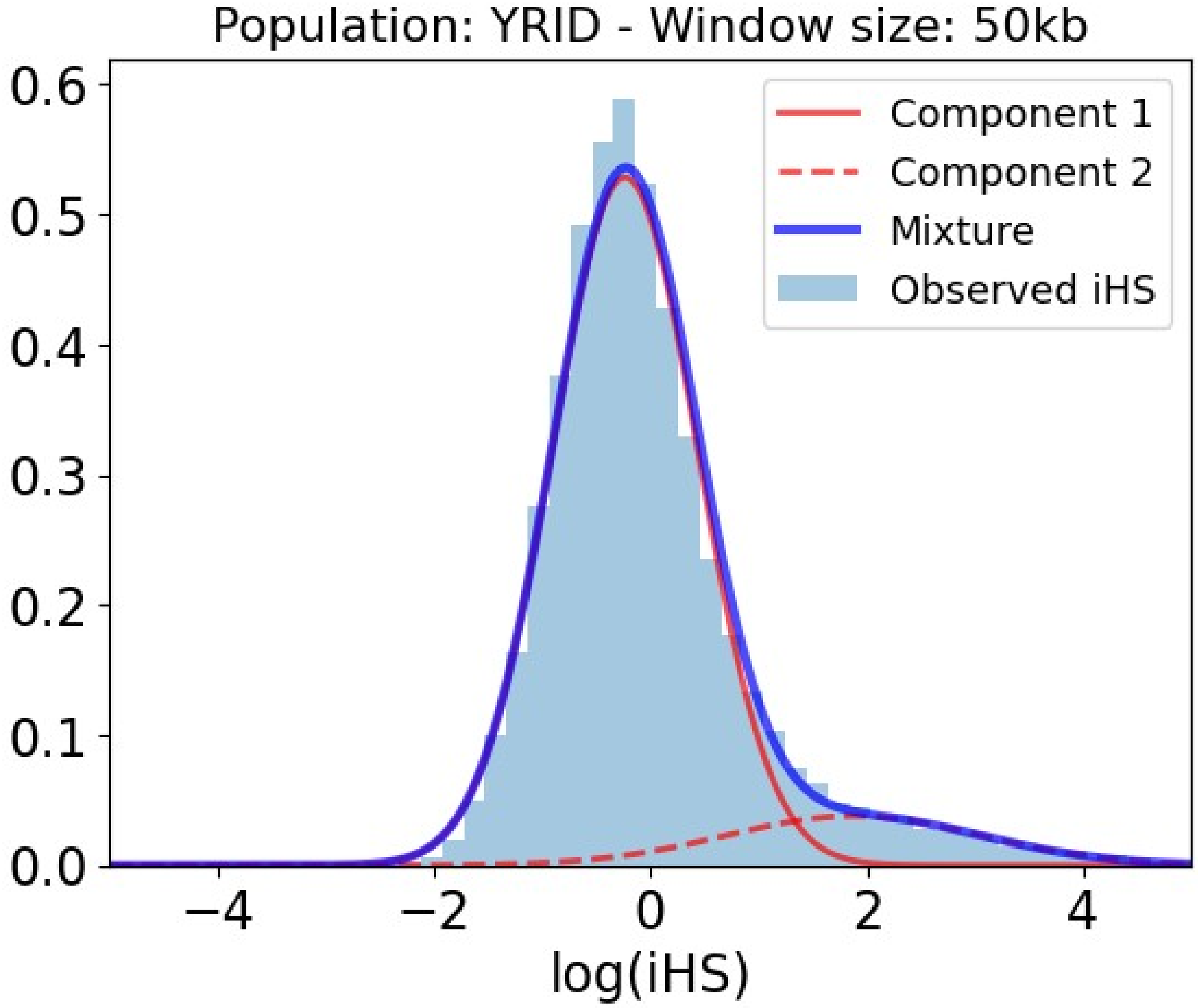


Table S1: Slopes and p-values of the association between iHS and genomic factors for the

Yoruba population in 50kb within the selection-enriched component.

| **Covariate** | **Slope** | **P-value** |
|---|---|---|
| Intercept | -115.603 | 0.000E+00 |
| Number iHS data points | 0.804 | 0.000E+00 |
| Density of conserved elements | -0.069 | 5.897E-01 |
| Recombination rate | -180.136 | 0.000E+00 |
| Number PPIs | 0.116 | 9.713E-02 |
| Regulatory density (ChIP-seq) | 0.211 | 3.791E-01 |
| Distance to VIPs | -0.025 | 7.000E-01 |
| Gene number | -0.012 | 9.123E-01 |
| Coding density | 0.054 | 6.526E-01 |
| Gene length | -0.077 | 4.490E-01 |
| Regulatory density in immune cells (ChIP-seq) | -0.447 | 5.618E-02 |
| Gene expression | 0.220 | 2.447E-01 |
| Gene expression in testis | 0.223 | 4.685E-02 |
| Gene expression in immune cells | 0.007 | 9.561E-01 |
| Regulatory density in testis (ChIP-seq) | -0.130 | 2.602E-01 |
| Regulatory density (DNaseI) | 0.188 | 4.407E-01 |
| GC-content | -0.323 | 6.891E-02 |

#### *Yoruba 100kb*

Figure S2: Mixture of Gaussian distributions fitting observed iHS (100kb windows) for Yoruba. The figure shows the two Gaussian distributions, component 1 and 2, being the latter enriched in positive selection. In that component, iHS linearly depends on the genomic factors considered. Legend: Light blue = Observed iHS; Dark blue = Mixture model; Full red curve = Component 1 of the mixture model; Dashed red curve = Component 2 of the mixture model enriched in selection.

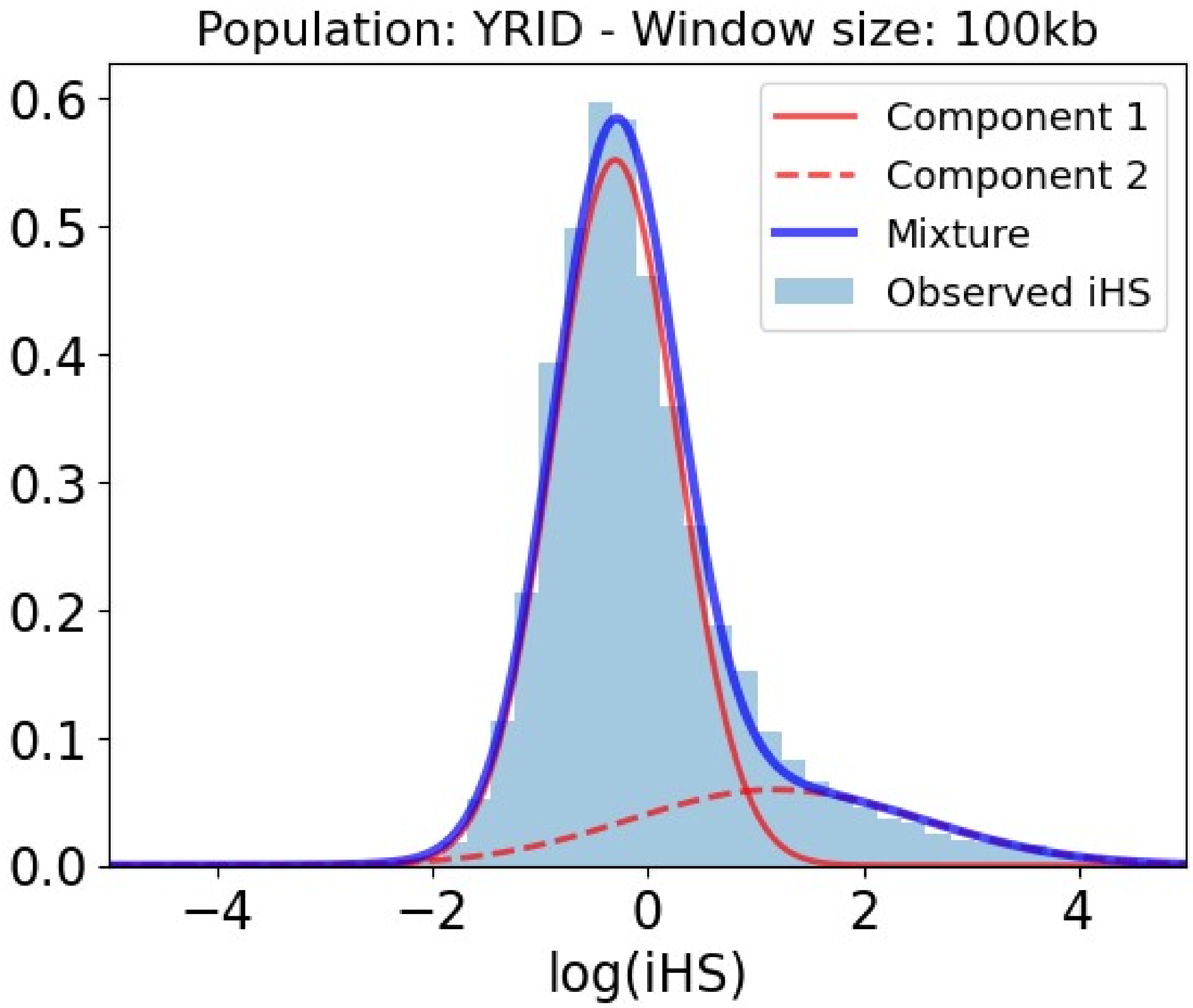


Table S2: Slopes and p-values of the association between iHS and genomic factors for the Yoruba population in 100kb within the selection-enriched component.

| **Covariate** | **Slope** | **P-value** |
| --- | --- | --- |
| Intercept | -2.740 | 0.000E+00 |
| Number iHS data points | -0.143 | 1.919E-04 |
| Density of conserved elements | 0.097 | 9.301E-02 |
| Recombination rate | -2.680 | 0.000E+00 |
| Number PPIs | 0.040 | 2.492E-01 |
| Regulatory density (ChIP-seq) | -0.186 | 4.779E-02 |
| Distance to VIPs | -0.024 | 5.115E-01 |
| Gene number | -0.095 | 1.096E-01 |
| Coding density | 0.140 | 2.953E-02 |
| Gene length | -0.040 | 3.348E-01 |
| Regulatory density in immune cells (ChIP-seq) | -0.326 | 1.181E-03 |

| **Covariate** | **Slope** | **P-value** |
|---|---|---|
| Gene expression | -0.082 | 3.189E-01 |
| Gene expression in testis | 0.200 | 3.742E-04 |
| Gene expression in immune cells | 0.434 | 2.118E-08 |
| Regulatory density in testis (ChIP-seq) | -0.038 | 4.838E-01 |
| Regulatory density (DNaseI) | -0.102 | 3.672E-01 |
| GC-content | -0.253 | 5.460E-03 |

#### *Yoruba 200kb*

Figure S3: Mixture of Gaussian distributions fitting observed iHS (200kb windows) for Yoruba. The figure shows the two Gaussian distributions, component 1 and 2, being the latter enriched in positive selection. In that component, iHS linearly depends on the genomic factors considered. Legend: Light blue = Observed iHS; Dark blue = Mixture model; Full red curve = Component 1 of the mixture model; Dashed red curve = Component 2 of the mixture model enriched in selection.

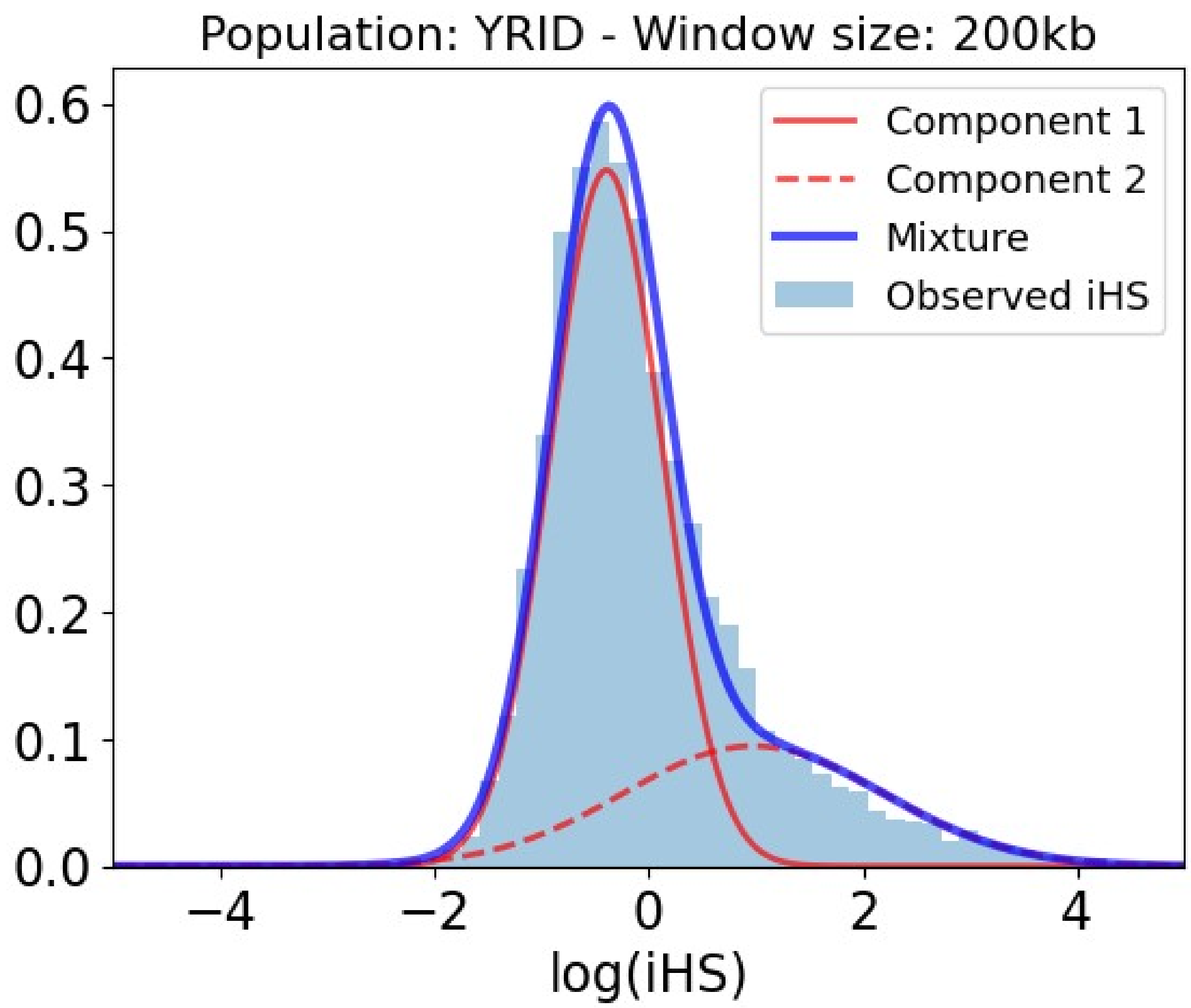


Table S3: Slopes and p-values of the association between iHS and genomic factors for the Yoruba population in 200kb within the selection-enriched component.

| Covariate | Slope | P-value |
|---|---|---|
| Intercept | -1.377 | 0.000E+00 |
| Number iHS data points | -0.336 | 0.000E+00 |
| Density of conserved elements | -0.058 | 2.247E-01 |
| Recombination rate | -1.290 | 0.000E+00 |
| Number PPIs | 0.022 | 4.770E-01 |
| Regulatory density (ChIP-seq) | -0.041 | 6.043E-01 |
| Distance to VIPs | -0.046 | 1.347E-01 |
| Gene number | 0.032 | 5.768E-01 |
| Coding density | 0.083 | 1.708E-01 |
| Gene length | -0.010 | 7.725E-01 |
| Regulatory density in immune cells (ChIP-seq) | -0.197 | 7.978E-03 |

| Covariate | Slope | P-value |
|---|---|---|
| Gene expression | -0.127 | 5.274E-02 |
| Gene expression in testis | 0.167 | 1.668E-04 |
| Gene expression in immune cells | 0.362 | 2.182E-09 |
| Regulatory density in testis (ChIP-seq) | -0.047 | 3.231E-01 |
| Regulatory density (DNaseI) | -0.088 | 3.746E-01 |
| GC-content | -0.181 | 2.904E-02 |

#### *Yoruba 500kb*

Figure S4: Mixture of Gaussian distributions fitting observed iHS (500kb windows) for Yoruba. The figure shows the two Gaussian distributions, component 1 and 2, being the latter enriched in positive selection. In that component, iHS linearly depends on the genomic factors considered. Legend: Light blue = Observed iHS; Dark blue = Mixture model; Full red curve = Component 1 of the mixture model; Dashed red curve = Component 2 of the mixture model enriched in selection.

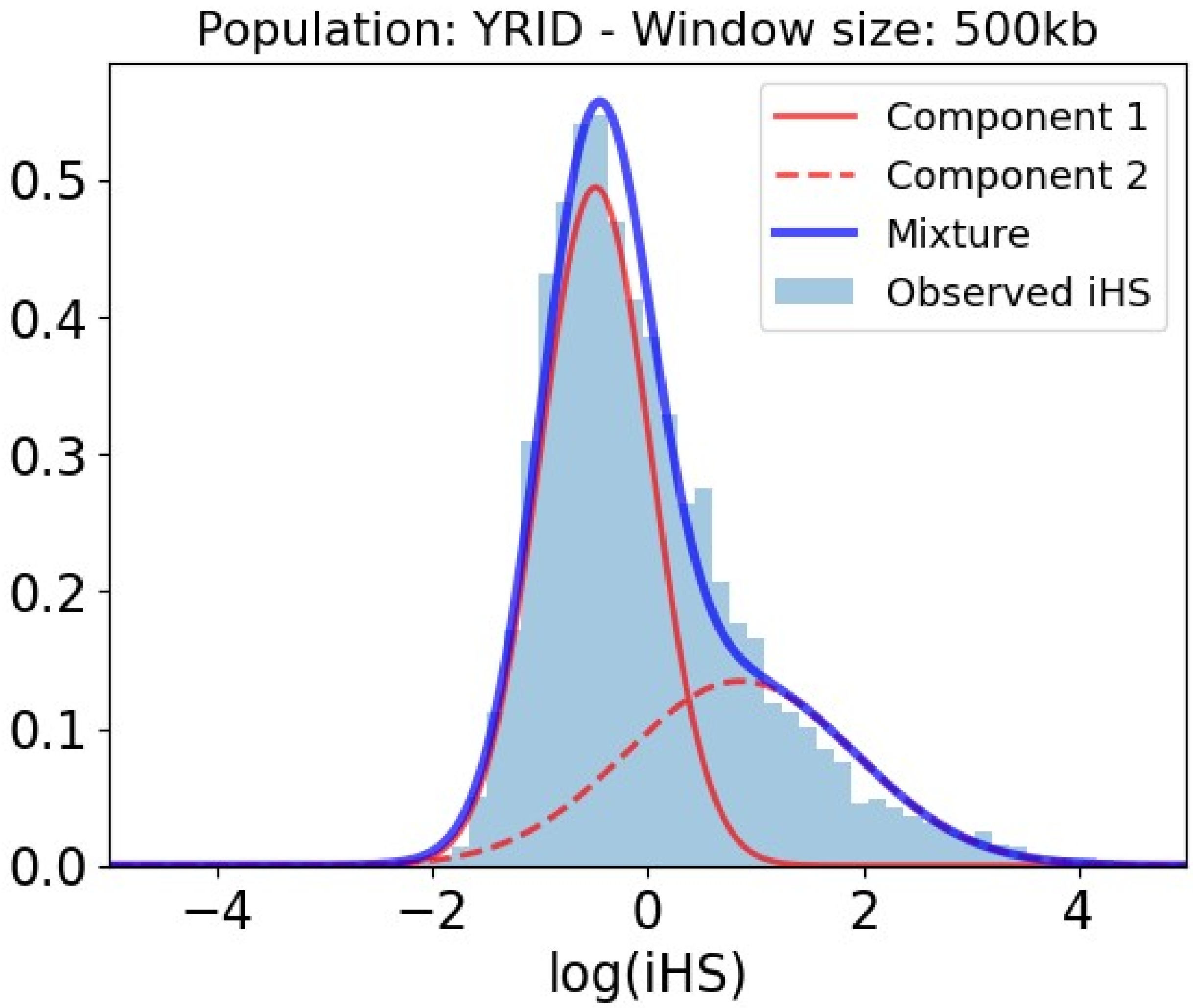


Table S4: Slopes and p-values of the association between iHS and genomic factors for the Yoruba population in 500kb within the selection-enriched component.

| Covariate | Slope | P-value |
|---|---|---|
| Intercept | -0.994 | 0.000E+00 |
| Number iHS data points | -0.305 | 3.331E-16 |
| Density of conserved elements | -0.030 | 4.976E-01 |
| Recombination rate | -1.560 | 0.000E+00 |
| Number PPIs | -0.044 | 1.559E-01 |
| Regulatory density (ChIP-seq) | -0.002 | 9.915E-01 |
| Distance to VIPs | -0.090 | 4.146E-03 |
| Gene number | 0.009 | 9.019E-01 |
| Coding density | 0.075 | 2.787E-01 |
| Gene length | -0.048 | 1.707E-01 |
| Regulatory density in immune cells (ChIP-seq) | 0.092 | 1.891E-01 |

| Covariate | Slope | P-value |
|---|---|---|
| Gene expression | -0.123 | 4.693E-02 |
| Gene expression in testis | 0.137 | 1.248E-03 |
| Gene expression in immune cells | 0.200 | 3.962E-04 |
| Regulatory density in testis (ChIP-seq) | -0.510 | 0.000E+00 |
| Regulatory density (DNaseI) | 0.062 | 5.669E-01 |
| GC-content | -0.035 | 6.941E-01 |

#### *Yoruba 1000kb*

Figure S5: Mixture of Gaussian distributions fitting observed iHS (1000kb windows) for Yoruba. The figure shows the two Gaussian distributions, component 1 and 2, being the latter enriched in positive selection. In that component, iHS linearly depends on the genomic factors considered. Legend: Light blue = Observed iHS; Dark blue = Mixture model; Full red curve = Component 1 of the mixture model; Dashed red curve = Component 2 of the mixture model enriched in selection.

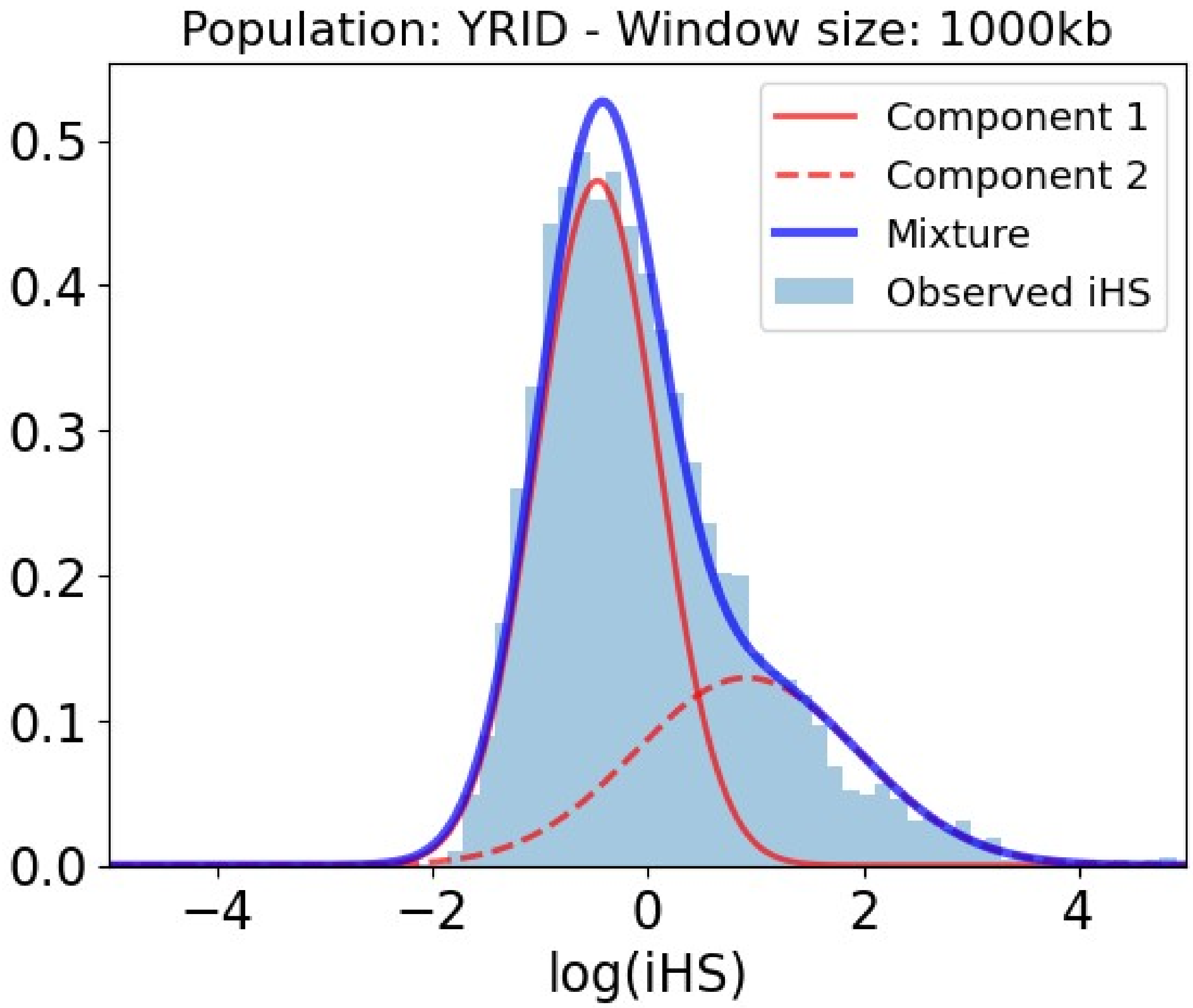


Table S5: Slopes and p-values of the association between iHS and genomic factors for the Yoruba population in 1000kb within the selection-enriched component.

| **Covariate** | **Slope** | **P-value** |
|---|---|---|
| Intercept | -1.426 | 0.000E+00 |
| Number iHS data points | -0.007 | 8.193E-01 |
| Density of conserved elements | 0.159 | 6.401E-04 |
| Recombination rate | -2.435 | 0.000E+00 |
| Number PPIs | -0.070 | 4.252E-02 |
| Regulatory density (ChIP-seq) | -0.147 | 2.372E-01 |
| Distance to VIPs | -0.170 | 4.161E-06 |
| Gene number | -0.086 | 3.458E-01 |
| Coding density | 0.159 | 8.412E-02 |
| Gene length | -0.059 | 1.125E-01 |
| Regulatory density in immune cells (ChIP-seq) | 0.047 | 6.243E-01 |

| Covariate | Slope | P-value |
|---|---|---|
| Gene expression | -0.139 | 3.768E-02 |
| Gene expression in testis | 0.001 | 9.900E-01 |
| Gene expression in immune cells | 0.265 | 1.512E-05 |
| Regulatory density in testis (ChIP-seq) | -0.795 | 0.000E+00 |
| Regulatory density (DNaseI) | -0.047 | 7.149E-01 |
| GC-content | 0.550 | 1.340E-06 |

### Utah residents

#### *Utah residents 50kb*

Figure S6: Mixture of Gaussian distributions fitting observed iHS (50kb windows) for Utah residents. The figure shows the two Gaussian distributions, component 1 and 2, being the latter enriched in positive selection. In that component, iHS linearly depends on the genomic factors considered. Legend: Light blue = Observed iHS; Dark blue = Mixture model; Full red curve = Component 1 of the mixture model; Dashed red curve = Component 2 of the mixture model enriched in selection.

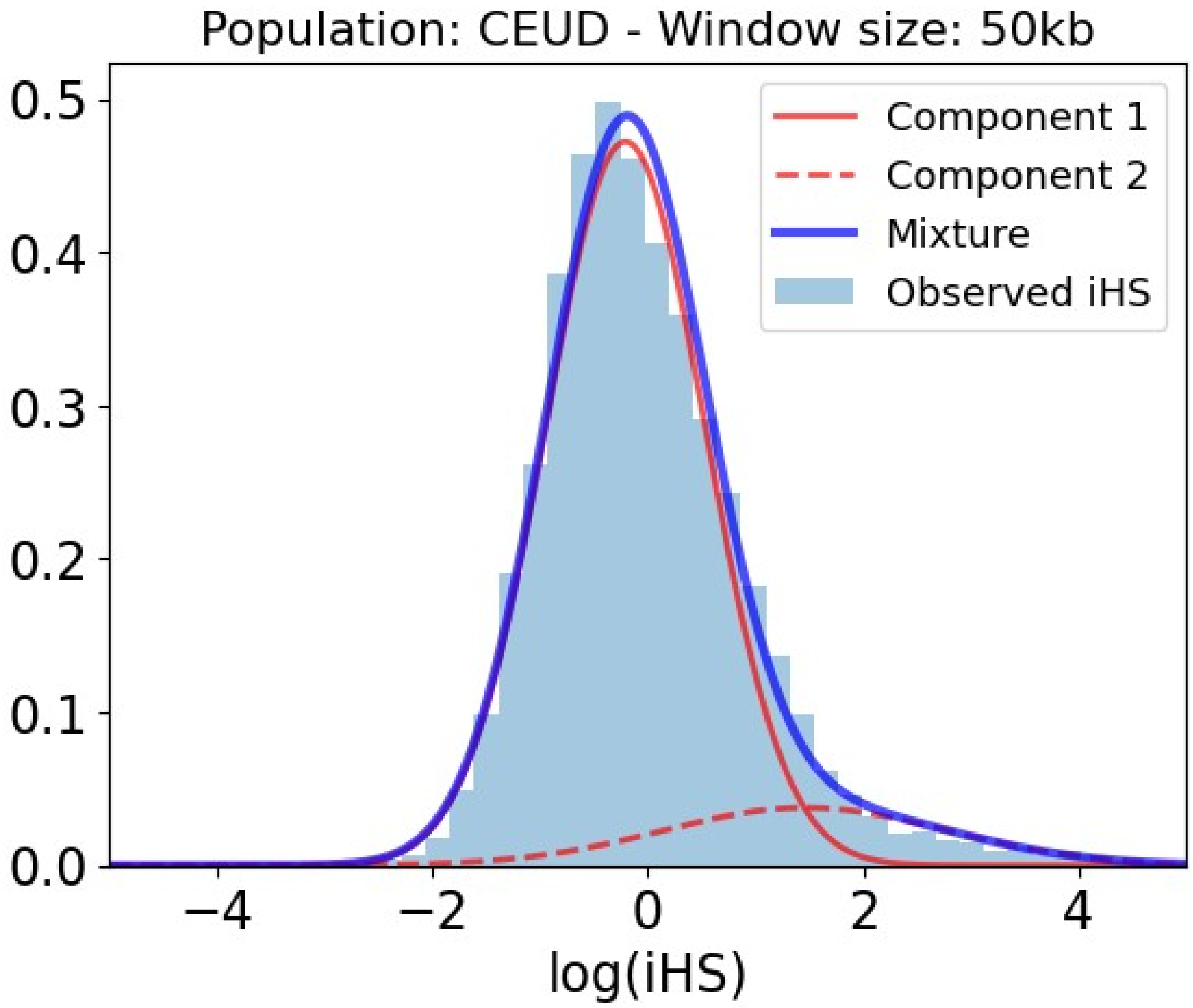


Table S6: Slopes and p-values of the association between iHS and genomic factors for the Utah residents population in 50kb within the selection-enriched component.

| **Covariate** | **Slope** | **P-value** |
|---|---|---|
| Intercept | -99.047 | 0.000E+00 |
| Number iHS data points | 0.966 | 0.000E+00 |
| Density of conserved elements | -0.108 | 5.375E-01 |
| Recombination rate | -154.830 | 0.000E+00 |
| Number PPIs | 0.140 | 1.122E-01 |
| Regulatory density (ChIP-seq) | 0.628 | 1.056E-02 |
| Distance to VIPs | -0.082 | 2.687E-01 |
| Gene number | 0.165 | 2.155E-01 |
| Coding density | 0.076 | 6.096E-01 |
| Gene length | 0.328 | 1.359E-02 |
| Regulatory density in immune cells (ChIP-seq) | -0.984 | 1.394E-04 |

| Covariate | Slope | P-value |
|---|---|---|
| Gene expression | 0.010 | 9.667E-01 |
| Gene expression in testis | 0.458 | 2.294E-03 |
| Gene expression in immune cells | 0.250 | 3.006E-01 |
| Regulatory density in testis (ChIP-seq) | 0.144 | 3.101E-01 |
| Regulatory density (DNaseI) | -0.338 | 2.616E-01 |
| GC-content | -0.067 | 7.399E-01 |

#### *Utah residents 100kb*

Figure S7: Mixture of Gaussian distributions fitting observed iHS (100kb windows) for Utah residents. The figure shows the two Gaussian distributions, component 1 and 2, being the latter enriched in positive selection. In that component, iHS linearly depends on the genomic factors considered. Legend: Light blue = Observed iHS; Dark blue = Mixture model; Full red curve = Component 1 of the mixture model; Dashed red curve = Component 2 of the mixture model enriched in selection.

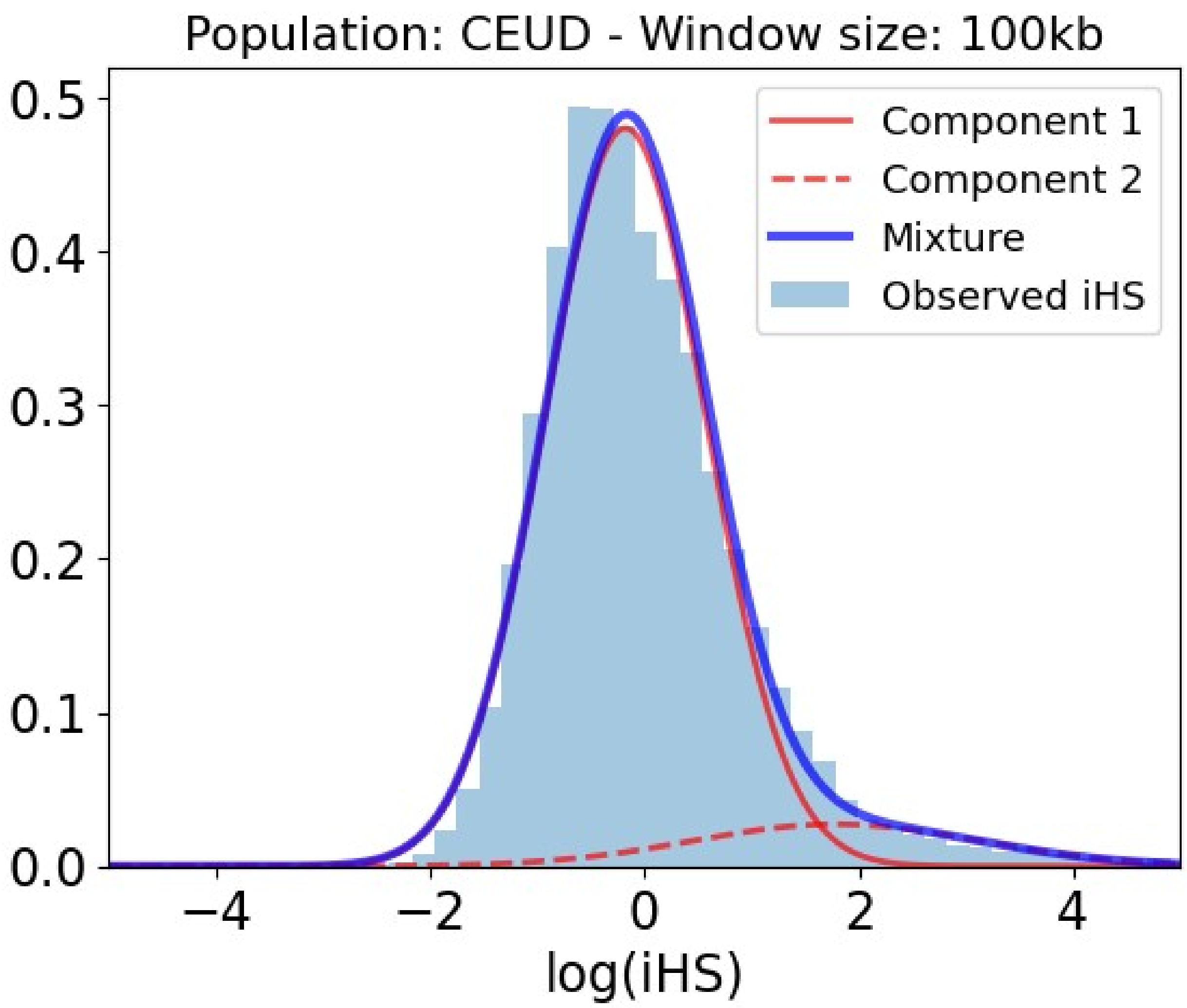


Table S7: Slopes and p-values of the association between iHS and genomic factors for the Utah residents population in 100kb within the selection-enriched component.

| **Covariate** | **Slope** | **P-value** |
|---|---|---|
| Intercept | -41.468 | 0.000E+00 |
| Number iHS data points | 0.558 | 2.960E-13 |
| Density of conserved elements | -0.028 | 8.499E-01 |
| Recombination rate | -52.197 | 0.000E+00 |
| Number PPIs | 0.080 | 2.794E-01 |
| Regulatory density (ChIP-seq) | -0.144 | 4.999E-01 |
| Distance to VIPs | -0.087 | 2.095E-01 |
| Gene number | 0.048 | 7.240E-01 |
| Coding density | -0.005 | 9.818E-01 |
| Gene length | 0.047 | 6.440E-01 |
| Regulatory density in immune cells (ChIP-seq) | -0.013 | 9.515E-01 |

| Covariate | Slope | P-value |
|---|---|---|
| Gene expression | 0.112 | 5.865E-01 |
| Gene expression in testis | 0.341 | 4.043E-03 |
| Gene expression in immune cells | 0.304 | 8.982E-02 |
| Regulatory density in testis (ChIP-seq) | -0.341 | 1.194E-02 |
| Regulatory density (DNaseI) | -0.117 | 6.535E-01 |
| GC-content | -0.034 | 8.598E-01 |

#### *Utah residents 200kb*

Figure S8: Mixture of Gaussian distributions fitting observed iHS (200kb windows) for Utah residents. The figure shows the two Gaussian distributions, component 1 and 2, being the latter enriched in positive selection. In that component, iHS linearly depends on the genomic factors considered. Legend: Light blue = Observed iHS; Dark blue = Mixture model; Full red curve = Component 1 of the mixture model; Dashed red curve = Component 2 of the mixture model enriched in selection.

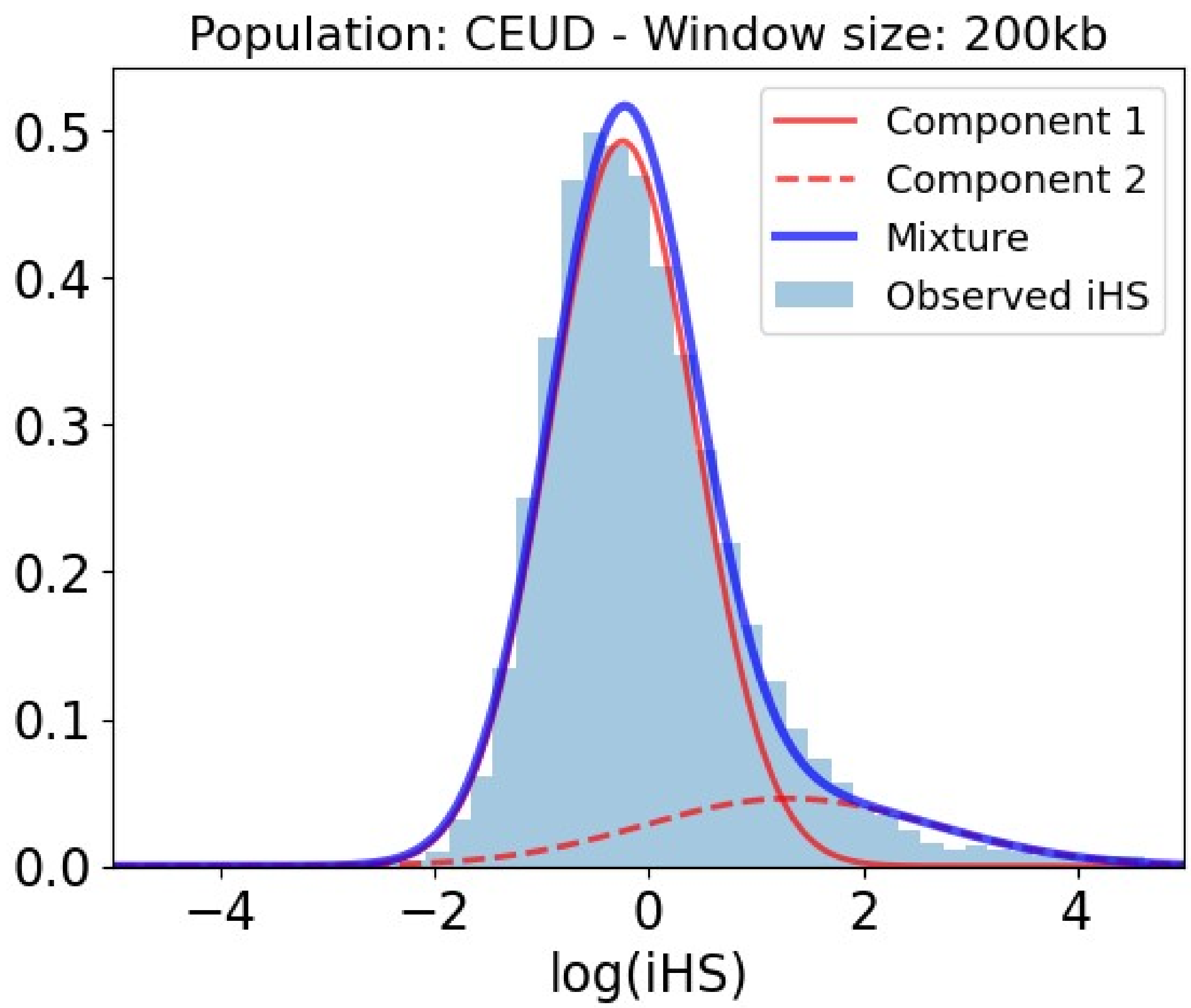


Table S8: Slopes and p-values of the association between iHS and genomic factors for the Utah residents population in 200kb within the selection-enriched component.

| **Covariate** | **Slope** | **P-value** |
|---|---|---|
| Intercept | -3.400 | 0.000E+00 |
| Number iHS data points | 0.035 | 1.692E-01 |
| Density of conserved elements | 0.262 | 7.451E-05 |
| Recombination rate | -3.015 | 0.000E+00 |
| Number PPIs | 0.047 | 2.519E-01 |
| Regulatory density (ChIP-seq) | -0.200 | 9.420E-02 |
| Distance to VIPs | -0.165 | 1.175E-04 |
| Gene number | -0.312 | 1.112E-03 |
| Coding density | 0.208 | 1.838E-02 |
| Gene length | 0.044 | 3.334E-01 |
| Regulatory density in immune cells (ChIP-seq) | -0.142 | 2.101E-01 |

| Covariate | Slope | P-value |
|---|---|---|
| Gene expression | -0.265 | 7.272E-03 |
| Gene expression in testis | 0.192 | 3.291E-03 |
| Gene expression in immune cells | 0.576 | 1.895E-10 |
| Regulatory density in testis (ChIP-seq) | -0.124 | 7.328E-02 |
| Regulatory density (DNaseI) | -0.140 | 3.582E-01 |
| GC-content | -0.123 | 2.845E-01 |

##### *Utah residents 500kb*

Figure S9: Mixture of Gaussian distributions fitting observed iHS (500kb windows) for Utah residents. The figure shows the two Gaussian distributions, component 1 and 2, being the latter enriched in positive selection. In that component, iHS linearly depends on the genomic factors considered. Legend: Light blue = Observed iHS; Dark blue = Mixture model; Full red curve = Component 1 of the mixture model; Dashed red curve = Component 2 of the mixture model enriched in selection.

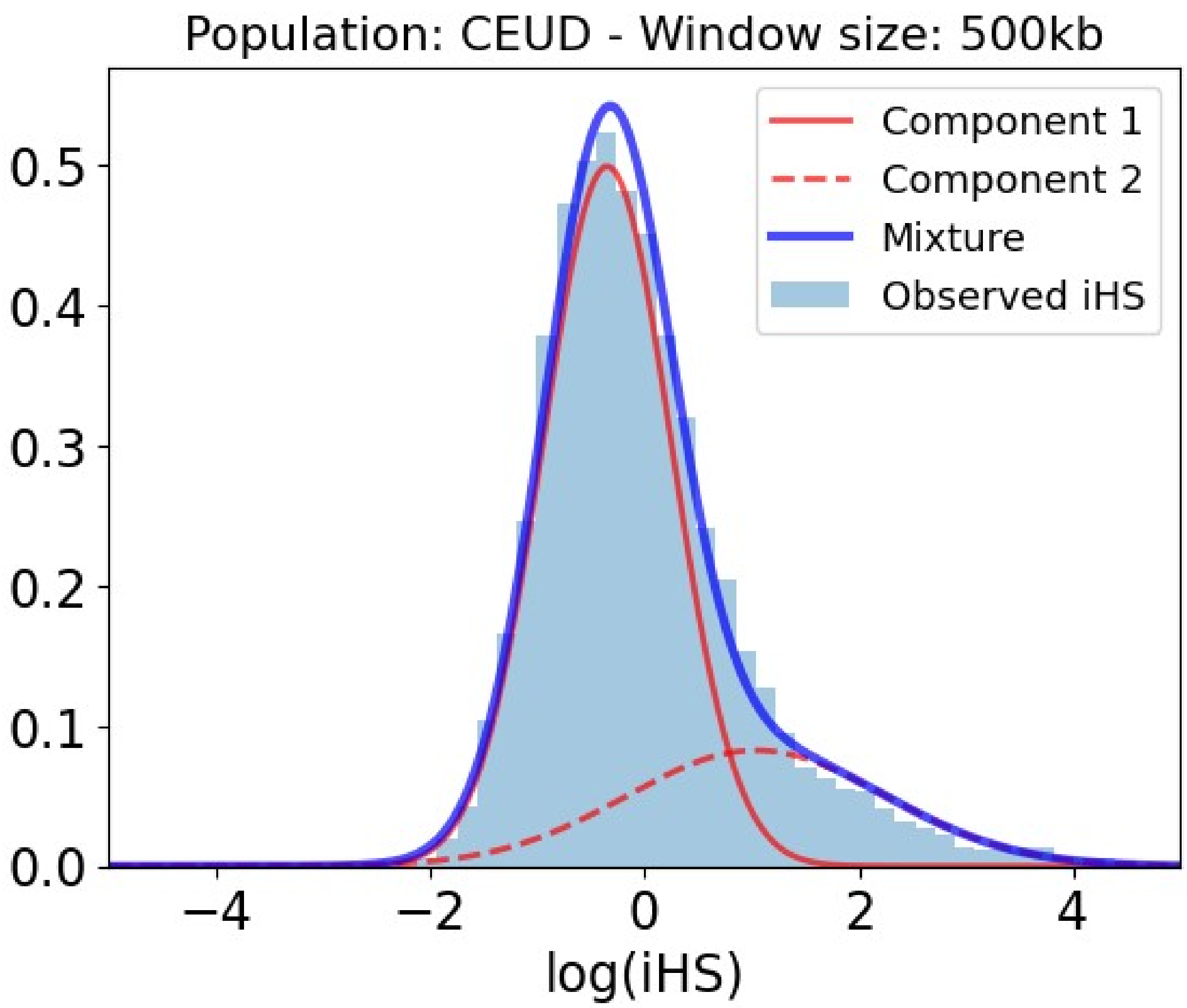


Table S9: Slopes and p-values of the association between iHS and genomic factors for the Utah residents population in 500kb within the selection-enriched component.

| **Covariate** | **Slope** | **P-value** |
|---|---|---|
| Intercept | -1.924 | 0.000E+00 |
| Number iHS data points | 0.075 | 5.070E-03 |
| Density of conserved elements | 0.183 | 4.455E-04 |
| Recombination rate | -2.127 | 0.000E+00 |
| Number PPIs | -0.038 | 2.921E-01 |
| Regulatory density (ChIP-seq) | 0.162 | 1.474E-01 |
| Distance to VIPs | -0.216 | 2.541E-08 |
| Gene number | -0.335 | 1.595E-04 |
| Coding density | 0.299 | 1.320E-03 |
| Gene length | 0.018 | 6.375E-01 |
| Regulatory density in immune cells (ChIP-seq) | 0.174 | 3.061E-02 |

| Covariate | Slope | P-value |
|---|---|---|
| Gene expression | -0.182 | 1.284E-02 |
| Gene expression in testis | 0.034 | 4.919E-01 |
| Gene expression in immune cells | 0.376 | 2.216E-08 |
| Regulatory density in testis (ChIP-seq) | -0.489 | 0.000E+00 |
| Regulatory density (DNaseI) | -0.270 | 4.698E-02 |
| GC-content | 0.097 | 3.703E-01 |

##### *Utah residents 1000kb*

Figure S10: Mixture of Gaussian distributions fitting observed iHS (1000kb windows) for Utah residents. The figure shows the two Gaussian distributions, component 1 and 2, being the latter enriched in positive selection. In that component, iHS linearly depends on the genomic factors considered. Legend: Light blue = Observed iHS; Dark blue = Mixture model; Full red curve = Component 1 of the mixture model; Dashed red curve = Component 2 of the mixture model enriched in selection.

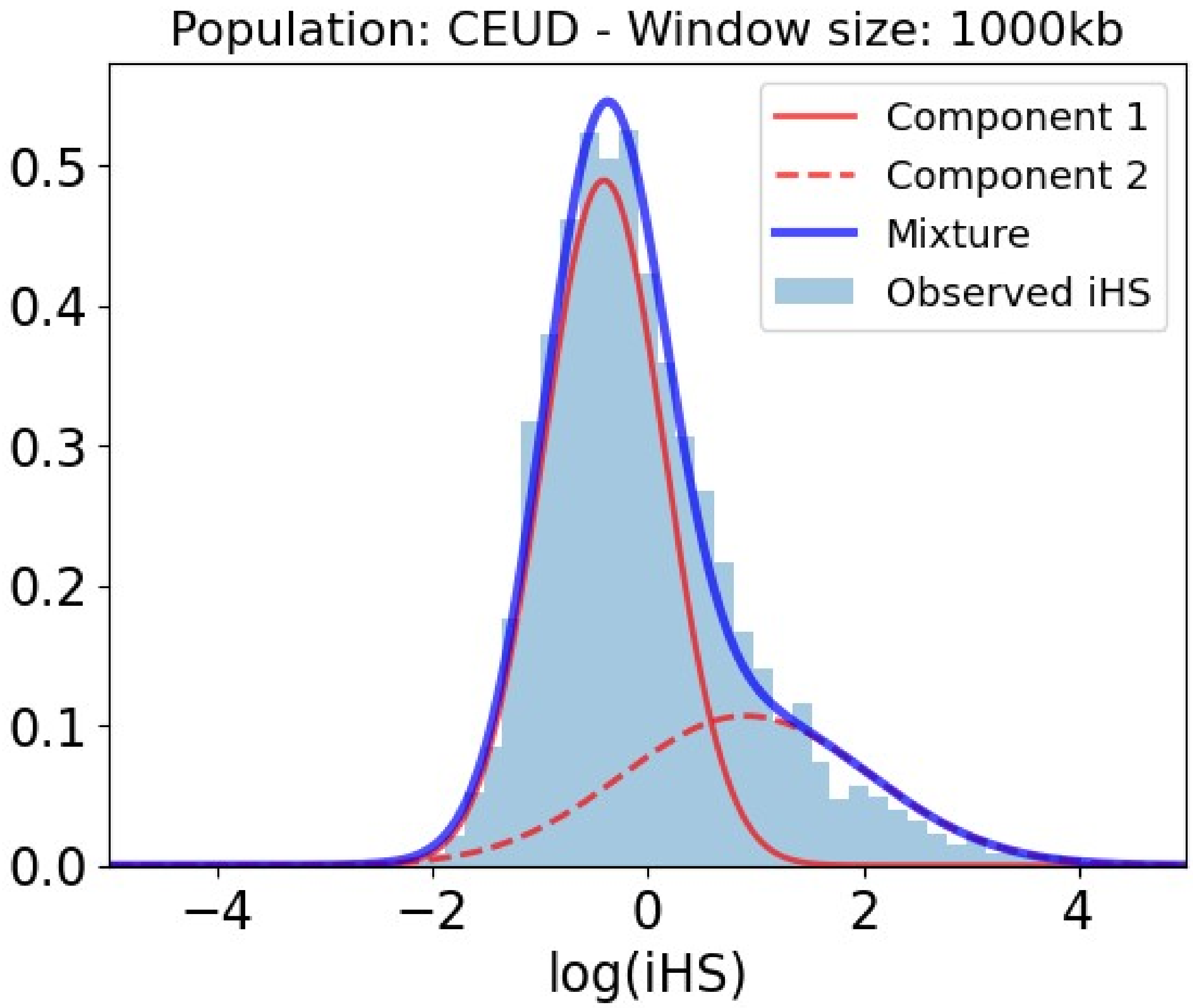


Table S10: Slopes and p-values of the association between iHS and genomic factors for the Utah residents population in 1000kb within the selection-enriched component.

| Covariate | Slope | P-value |
|---|---|---|
| Intercept | -1.766 | 0.000E+00 |
| Number iHS data points | 0.315 | 0.000E+00 |
| Density of conserved elements | 0.379 | 7.738E-13 |
| Recombination rate | -2.798 | 0.000E+00 |
| Number PPIs | -0.050 | 1.951E-01 |
| Regulatory density (ChIP-seq) | 0.593 | 3.450E-06 |
| Distance to VIPs | -0.337 | 8.449E-14 |
| Gene number | -0.521 | 1.284E-06 |
| Coding density | 0.535 | 8.716E-06 |
| Gene length | -0.047 | 2.840E-01 |
| Regulatory density in immune cells (ChIP-seq) | -0.051 | 6.010E-01 |

| Covariate | Slope | P-value |
|---|---|---|
| Gene expression | -0.335 | 9.695E-06 |
| Gene expression in testis | -0.031 | 5.450E-01 |
| Gene expression in immune cells | 0.408 | 6.868E-09 |
| Regulatory density in testis (ChIP-seq) | -1.031 | 0.000E+00 |
| Regulatory density (DNaseI) | -0.609 | 5.447E-05 |
| GC-content | 0.745 | 1.350E-08 |

### Toscani

#### *Toscani 50kb*

Figure S11: Mixture of Gaussian distributions fitting observed iHS (50kb windows) for Toscani. The figure shows the two Gaussian distributions, component 1 and 2, being the latter enriched in positive selection. In that component, iHS linearly depends on the genomic factors considered. Legend: Light blue = Observed iHS; Dark blue = Mixture model; Full red curve = Component 1 of the mixture model; Dashed red curve = Component 2 of the mixture model enriched in selection.

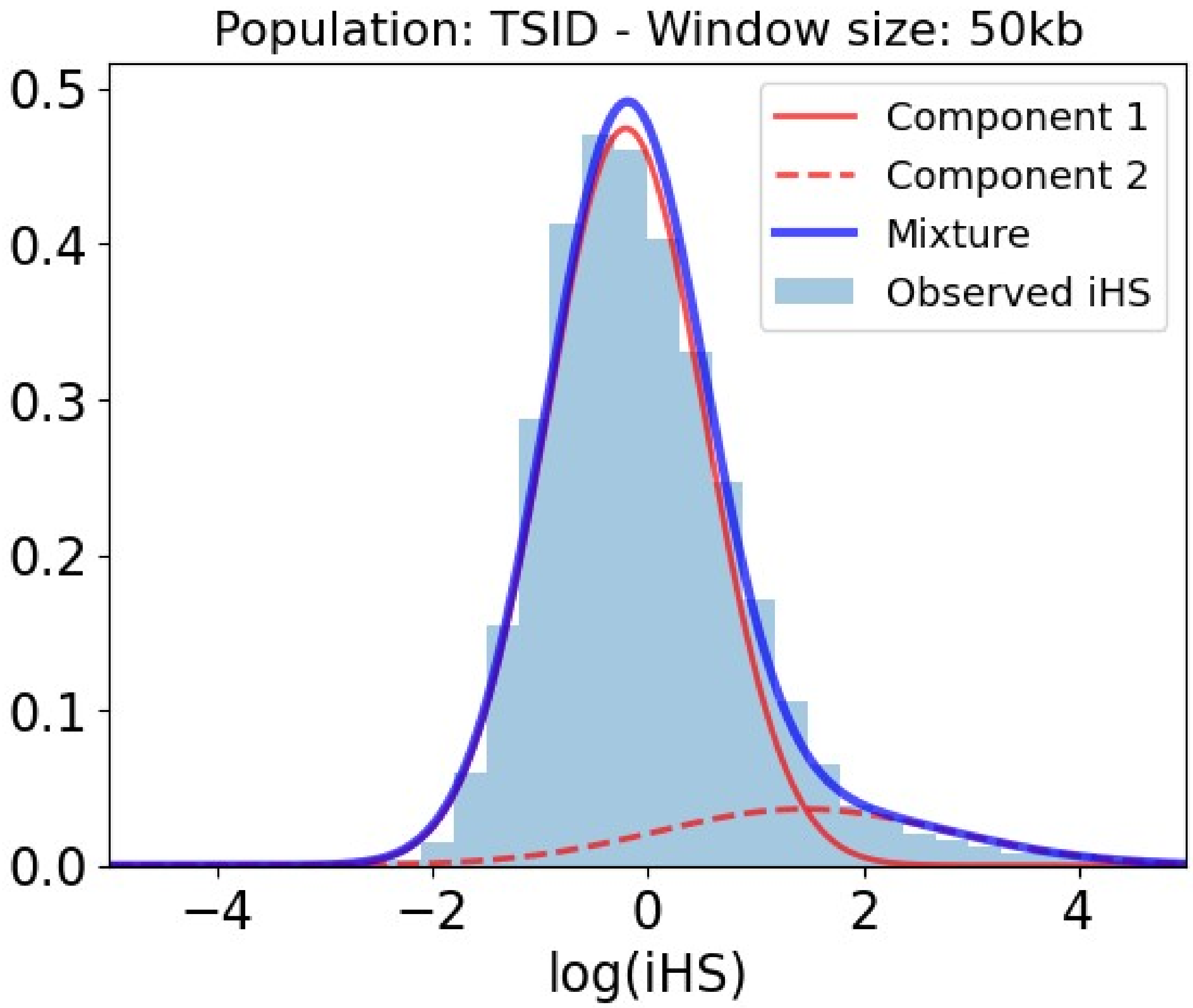


Table S11: Slopes and p-values of the association between iHS and genomic factors for the Toscani population in 50kb within the selection-enriched component.

| Covariate | Slope | P-value |
|---|---|---|
| Intercept | -107.704 | 0.000E+00 |
| Number iHS data points | 1.332 | 0.000E+00 |
| Density of conserved elements | 0.016 | 9.181E-01 |
| Recombination rate | -168.545 | 0.000E+00 |
| Number PPIs | 0.266 | 3.654E-03 |
| Regulatory density (ChIP-seq) | 0.667 | 3.454E-02 |
| Distance to VIPs | -0.088 | 2.359E-01 |
| Gene number | 0.171 | 2.920E-01 |
| Coding density | 0.129 | 4.685E-01 |
| Gene length | 0.136 | 2.698E-01 |
| Regulatory density in immune cells (ChIP-seq) | -0.909 | 5.142E-03 |

| Covariate | Slope | P-value |
|---|---|---|
| Gene expression | 0.151 | 5.291E-01 |
| Gene expression in testis | 0.363 | 1.632E-02 |
| Gene expression in immune cells | 0.194 | 4.504E-01 |
| Regulatory density in testis (ChIP-seq) | 0.072 | 6.815E-01 |
| Regulatory density (DNaseI) | -0.310 | 3.124E-01 |
| GC-content | -0.078 | 7.724E-01 |

#### *Toscani 100kb*

Figure S12: Mixture of Gaussian distributions fitting observed iHS (100kb windows) for Toscani. The figure shows the two Gaussian distributions, component 1 and 2, being the latter enriched in positive selection. In that component, iHS linearly depends on the genomic factors considered. Legend: Light blue = Observed iHS; Dark blue = Mixture model; Full red curve = Component 1 of the mixture model; Dashed red curve = Component 2 of the mixture model enriched in selection.

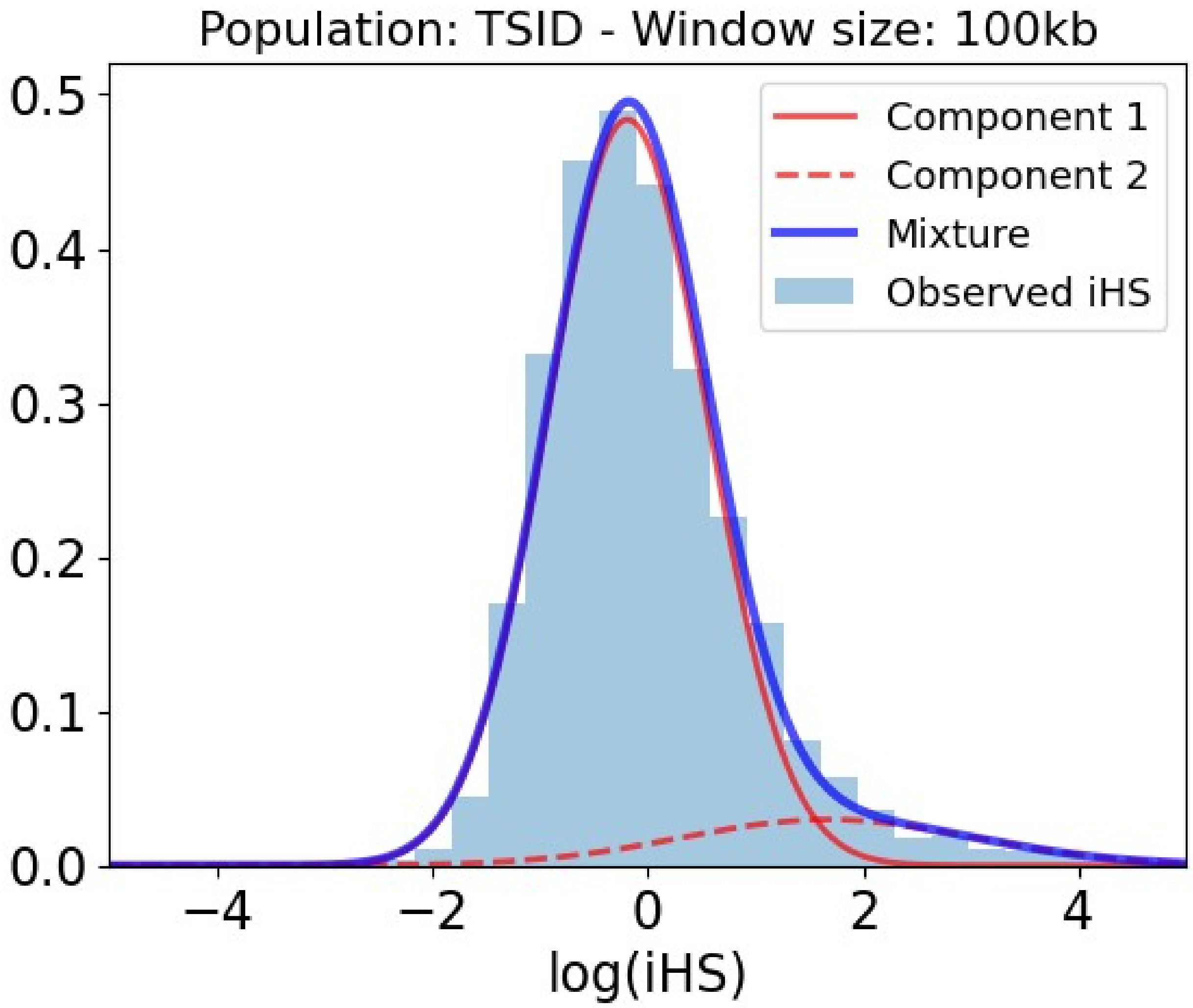


Table S12: Slopes and p-values of the association between iHS and genomic factors for the Toscani population in 100kb within the selection-enriched component.

| **Covariate** | **Slope** | **P-value** |
|---|---|---|
| Intercept | -23.200 | 0.000E+00 |
| Number iHS data points | 0.499 | 3.463E-12 |
| Density of conserved elements | 0.143 | 2.029E-01 |
| Recombination rate | -29.339 | 0.000E+00 |
| Number PPIs | 0.153 | 9.134E-03 |
| Regulatory density (ChIP-seq) | 0.131 | 4.527E-01 |
| Distance to VIPs | -0.065 | 2.178E-01 |
| Gene number | 0.024 | 8.024E-01 |
| Coding density | -0.014 | 8.975E-01 |
| Gene length | 0.108 | 1.694E-01 |
| Regulatory density in immune cells (ChIP-seq) | -0.143 | 4.497E-01 |

| Covariate | Slope | P-value |
|---|---|---|
| Gene expression | -0.154 | 3.359E-01 |
| Gene expression in testis | 0.253 | 9.575E-03 |
| Gene expression in immune cells | 0.387 | 9.657E-03 |
| Regulatory density in testis (ChIP-seq) | -0.134 | 1.789E-01 |
| Regulatory density (DNaseI) | -0.107 | 5.990E-01 |
| GC-content | -0.131 | 4.052E-01 |

#### *Toscani 200kb*

Figure S13: Mixture of Gaussian distributions fitting observed iHS (200kb windows) for Toscani. The figure shows the two Gaussian distributions, component 1 and 2, being the latter enriched in positive selection. In that component, iHS linearly depends on the genomic factors considered. Legend: Light blue = Observed iHS; Dark blue = Mixture model; Full red curve = Component 1 of the mixture model; Dashed red curve = Component 2 of the mixture model enriched in selection.

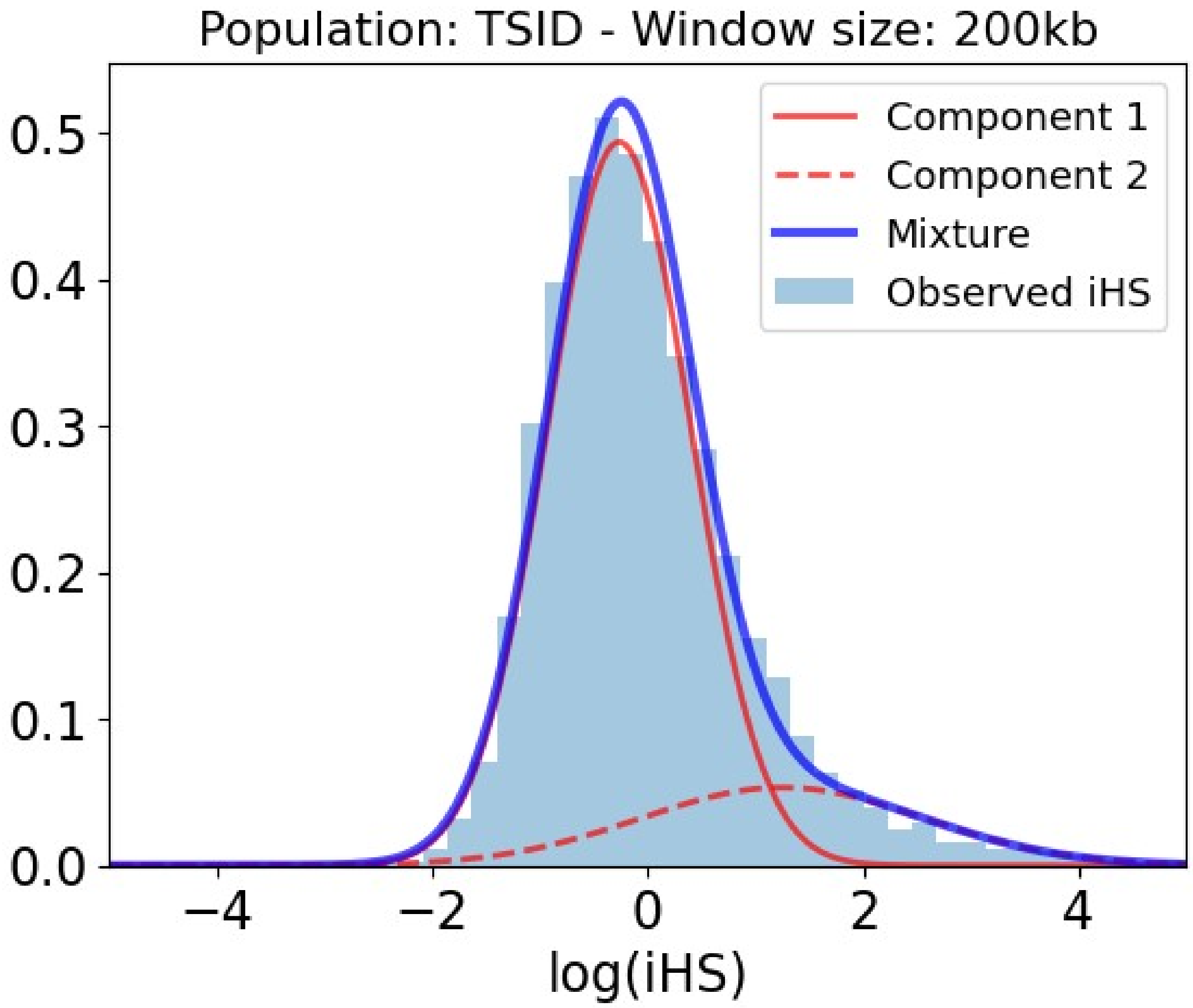


Table S13: Slopes and p-values of the association between iHS and genomic factors for the Toscani population in 200kb within the selection-enriched component.

| **Covariate** | **Slope** | **P-value** |
| --- | --- | --- |
| Intercept | -2.868 | 0.000E+00 |
| Number iHS data points | 0.073 | 1.139E-02 |
| Density of conserved elements | 0.182 | 2.434E-03 |
| Recombination rate | -2.571 | 0.000E+00 |
| Number PPIs | 0.054 | 1.503E-01 |
| Regulatory density (ChIP-seq) | 0.164 | 1.165E-01 |
| Distance to VIPs | -0.154 | 1.622E-04 |
| Gene number | -0.112 | 1.334E-01 |
| Coding density | 0.180 | 2.083E-02 |
| Gene length | 0.089 | 3.656E-02 |
| Regulatory density in immune cells (ChIP-seq) | -0.186 | 4.183E-02 |

| Covariate | Slope | P-value |
|---|---|---|
| Gene expression | -0.355 | 6.950E-05 |
| Gene expression in testis | 0.260 | 1.088E-05 |
| Gene expression in immune cells | 0.526 | 1.051E-10 |
| Regulatory density in testis (ChIP-seq) | 0.071 | 2.765E-01 |
| Regulatory density (DNaseI) | -0.188 | 1.610E-01 |
| GC-content | -0.339 | 1.435E-03 |

#### *Toscani 500kb*

Figure S14: Mixture of Gaussian distributions fitting observed iHS (500kb windows) for Toscani. The figure shows the two Gaussian distributions, component 1 and 2, being the latter enriched in positive selection. In that component, iHS linearly depends on the genomic factors considered. Legend: Light blue = Observed iHS; Dark blue = Mixture model; Full red curve = Component 1 of the mixture model; Dashed red curve = Component 2 of the mixture model enriched in selection.

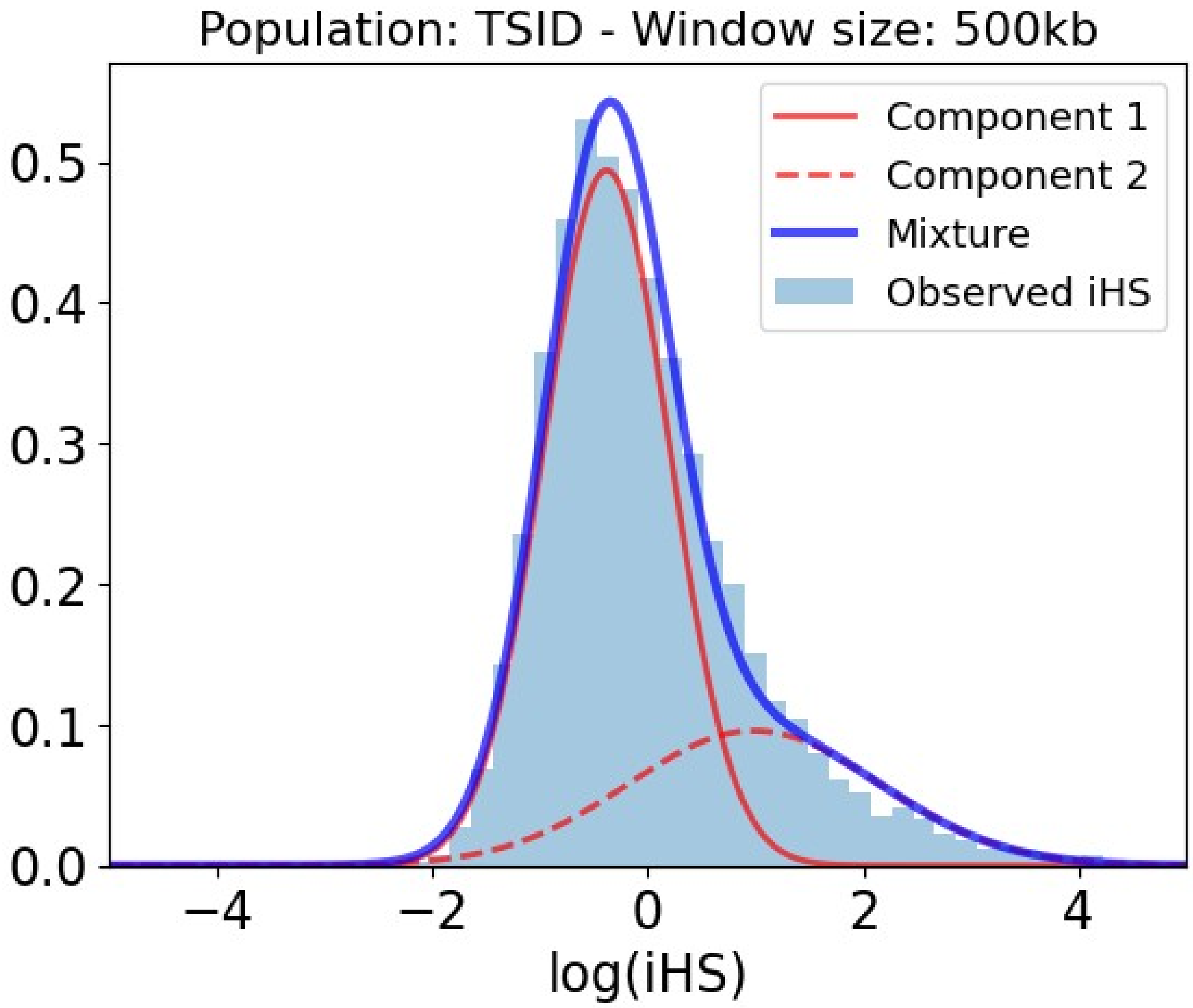


Table S14: Slopes and p-values of the association between iHS and genomic factors for the Toscani population in 500kb within the selection-enriched component.

| **Covariate** | **Slope** | **P-value** |
|---|---|---|
| Intercept | -1.679 | 0.000E+00 |
| Number iHS data points | 0.092 | 4.913E-04 |
| Density of conserved elements | 0.213 | 1.402E-05 |
| Recombination rate | -2.025 | 0.000E+00 |
| Number PPIs | -0.024 | 4.767E-01 |
| Regulatory density (ChIP-seq) | 0.463 | 2.710E-06 |
| Distance to VIPs | -0.154 | 4.266E-05 |
| Gene number | -0.199 | 1.038E-02 |
| Coding density | 0.344 | 2.734E-05 |
| Gene length | 0.004 | 9.115E-01 |
| Regulatory density in immune cells (ChIP-seq) | 0.121 | 1.064E-01 |

| Covariate | Slope | P-value |
|---|---|---|
| Gene expression | -0.182 | 8.404E-03 |
| Gene expression in testis | 0.066 | 1.686E-01 |
| Gene expression in immune cells | 0.333 | 2.317E-07 |
| Regulatory density in testis (ChIP-seq) | -0.444 | 3.275E-14 |
| Regulatory density (DNaseI) | -0.324 | 1.123E-02 |
| GC-content | -0.138 | 1.985E-01 |

#### *Toscani 1000kb*

Figure S15: Mixture of Gaussian distributions fitting observed iHS (1000kb windows) for Toscani. The figure shows the two Gaussian distributions, component 1 and 2, being the latter enriched in positive selection. In that component, iHS linearly depends on the genomic factors considered. Legend: Light blue = Observed iHS; Dark blue = Mixture model; Full red curve = Component 1 of the mixture model; Dashed red curve = Component 2 of the mixture model enriched in selection.

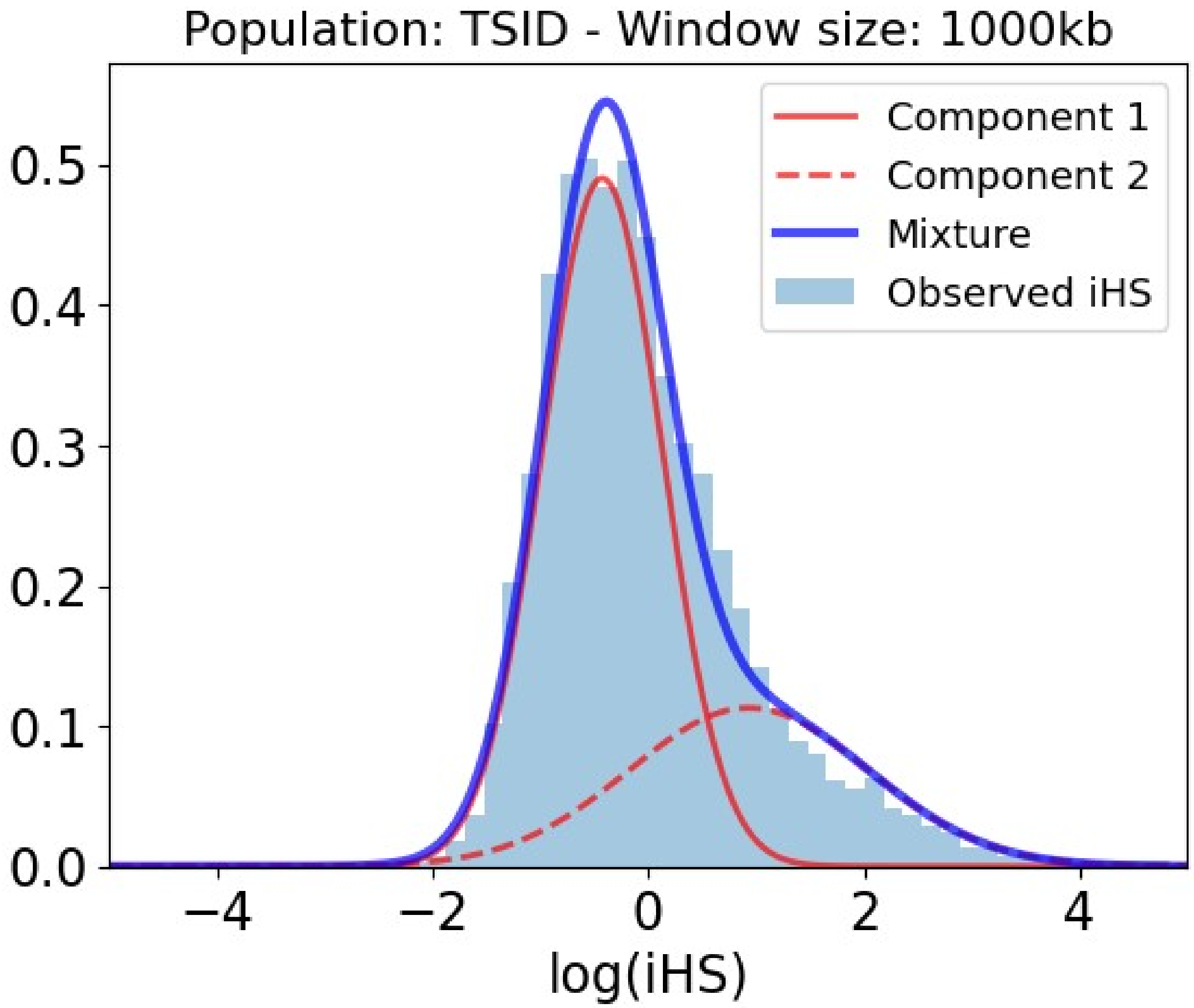


Table S15: Slopes and p-values of the association between iHS and genomic factors for the Toscani population in 1000kb within the selection-enriched component.

| **Covariate** | **Slope** | **P-value** |
|---|---|---|
| Intercept | -1.660 | 0.000E+00 |
| Number iHS data points | 0.330 | 0.000E+00 |
| Density of conserved elements | 0.337 | 2.701E-11 |
| Recombination rate | -2.696 | 0.000E+00 |
| Number PPIs | -0.041 | 2.671E-01 |
| Regulatory density (ChIP-seq) | 0.742 | 3.063E-10 |
| Distance to VIPs | -0.266 | 1.917E-10 |
| Gene number | -0.397 | 5.912E-05 |
| Coding density | 0.586 | 3.772E-08 |
| Gene length | -0.024 | 5.588E-01 |
| Regulatory density in immune cells (ChIP-seq) | -0.079 | 3.944E-01 |

| Covariate | Slope | P-value |
|---|---|---|
| Gene expression | -0.253 | 4.915E-04 |
| Gene expression in testis | -0.040 | 4.213E-01 |
| Gene expression in immune cells | 0.366 | 5.339E-08 |
| Regulatory density in testis (ChIP-seq) | -0.861 | 0.000E+00 |
| Regulatory density (DNaseI) | -0.635 | 8.237E-06 |
| GC-content | 0.516 | 2.469E-05 |

### Han Chinese

#### *Han Chinese 50kb*

Figure S16: Mixture of Gaussian distributions fitting observed iHS (50kb windows) for Han Chinese. The figure shows the two Gaussian distributions, component 1 and 2, being the latter enriched in positive selection. In that component, iHS linearly depends on the genomic factors considered. Legend: Light blue = Observed iHS; Dark blue = Mixture model; Full red curve = Component 1 of the mixture model; Dashed red curve = Component 2 of the mixture model enriched in selection.

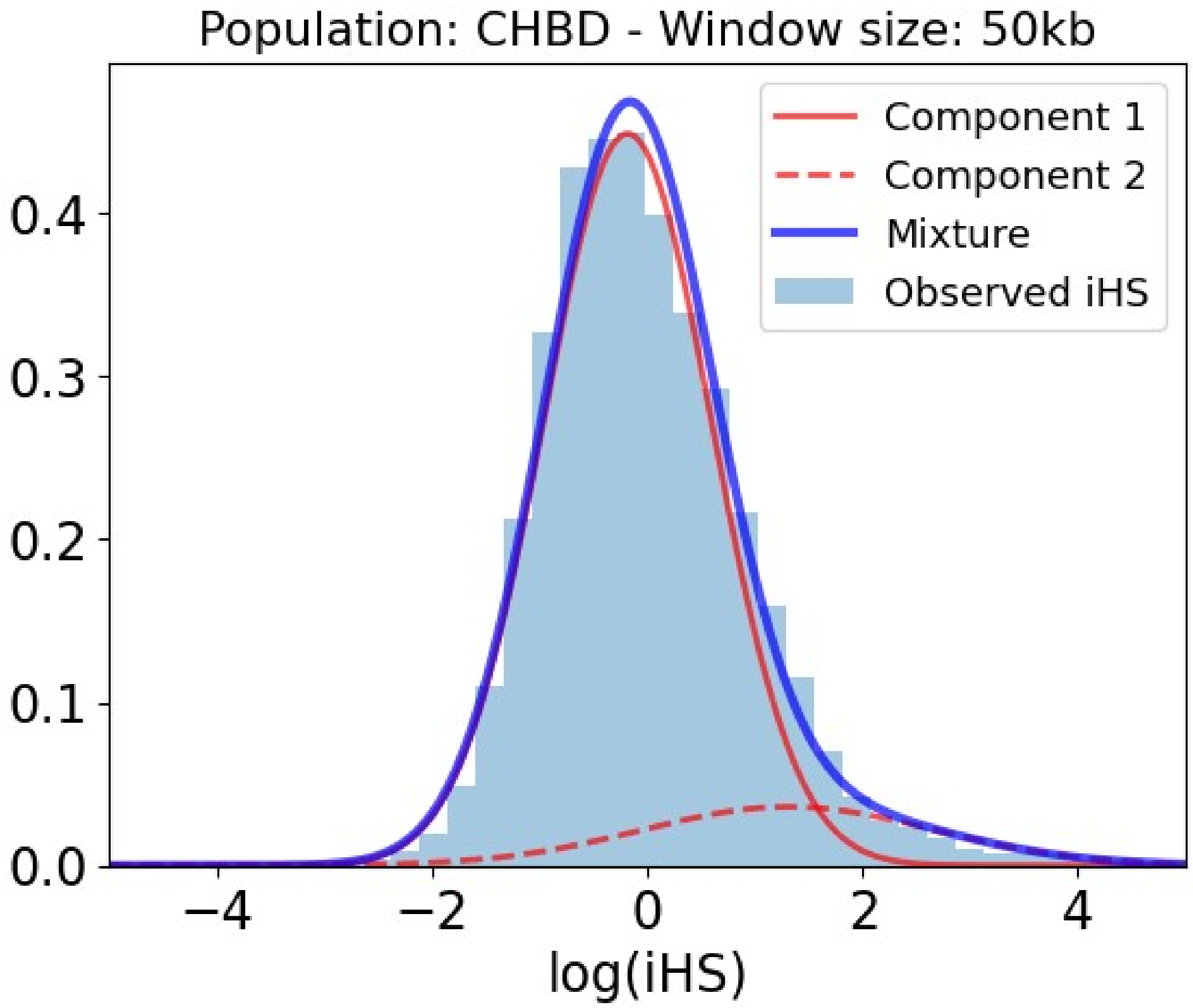


Table S16: Slopes and p-values of the association between iHS and genomic factors for the Han Chinese population in 50kb within the selection-enriched component.

| **Covariate** | **Slope** | **P-value** |
|---|---|---|
| Intercept | -92.249 | 0.000E+00 |
| Number iHS data points | 1.219 | 0.000E+00 |
| Density of conserved elements | -0.122 | 4.844E-01 |
| Recombination rate | -144.576 | 0.000E+00 |
| Number PPIs | 0.075 | 4.238E-01 |
| Regulatory density (ChIP-seq) | 0.320 | 1.732E-01 |
| Distance to VIPs | -0.108 | 9.944E-02 |
| Gene number | 0.119 | 3.918E-01 |
| Coding density | 0.258 | 1.073E-01 |
| Gene length | 0.907 | 2.363E-06 |
| Regulatory density in immune cells (ChIP-seq) | -0.999 | 1.356E-04 |

| Covariate | Slope | P-value |
|---|---|---|
| Gene expression | -0.594 | 2.694E-02 |
| Gene expression in testis | 0.204 | 2.022E-01 |
| Gene expression in immune cells | 0.855 | 8.659E-04 |
| Regulatory density in testis (ChIP-seq) | 0.004 | 9.712E-01 |
| Regulatory density (DNaseI) | 0.261 | 4.141E-01 |
| GC-content | 0.033 | 9.028E-01 |

#### *Han Chinese 100kb*

Figure S17: Mixture of Gaussian distributions fitting observed iHS (100kb windows) for Han Chinese. The figure shows the two Gaussian distributions, component 1 and 2, being the latter enriched in positive selection. In that component, iHS linearly depends on the genomic factors considered. Legend: Light blue = Observed iHS; Dark blue = Mixture model; Full red curve = Component 1 of the mixture model; Dashed red curve = Component 2 of the mixture model enriched in selection.

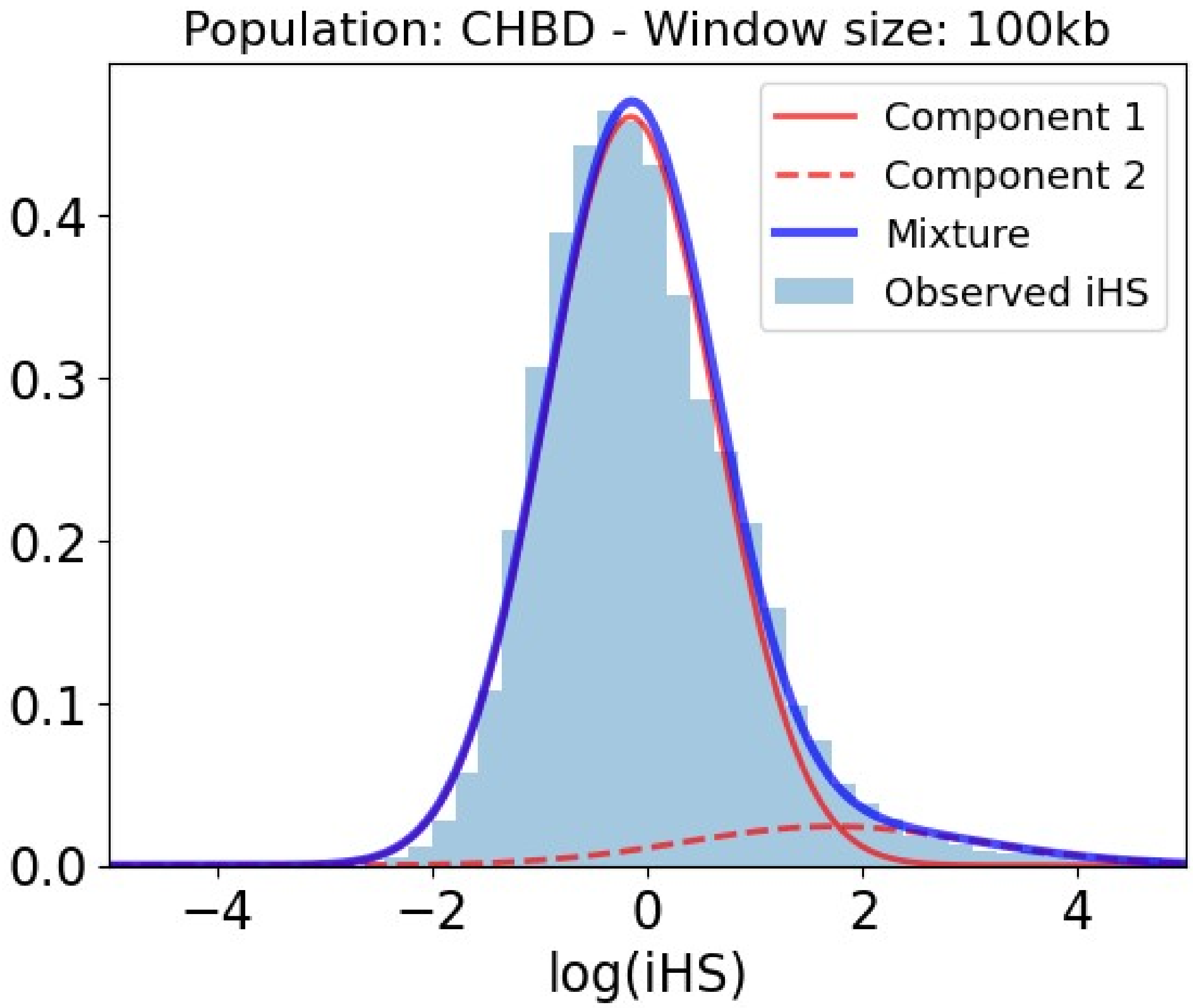


Table S17: Slopes and p-values of the association between iHS and genomic factors for the Han Chinese population in 100kb within the selection-enriched component.

| **Covariate** | **Slope** | **P-value** |
|---|---|---|
| Intercept | -58.905 | 0.000E+00 |
| Number iHS data points | 1.538 | 0.000E+00 |
| Density of conserved elements | 0.085 | 6.065E-01 |
| Recombination rate | -74.503 | 0.000E+00 |
| Number PPIs | 0.137 | 8.902E-02 |
| Regulatory density (ChIP-seq) | 0.490 | 8.604E-02 |
| Distance to VIPs | -0.048 | 5.595E-01 |
| Gene number | 0.466 | 2.526E-03 |
| Coding density | -0.160 | 3.587E-01 |
| Gene length | 0.585 | 5.204E-04 |
| Regulatory density in immune cells (ChIP-seq) | -0.768 | 5.987E-03 |

| Covariate | Slope | P-value |
|---|---|---|
| Gene expression | -0.345 | 1.618E-01 |
| Gene expression in testis | 0.275 | 5.362E-02 |
| Gene expression in immune cells | 0.701 | 1.941E-03 |
| Regulatory density in testis (ChIP-seq) | 0.114 | 3.893E-01 |
| Regulatory density (DNaseI) | -0.041 | 9.046E-01 |
| GC-content | -0.060 | 7.931E-01 |

#### *Han Chinese 200kb*

Figure S18: Mixture of Gaussian distributions fitting observed iHS (200kb windows) for Han Chinese. The figure shows the two Gaussian distributions, component 1 and 2, being the latter enriched in positive selection. In that component, iHS linearly depends on the genomic factors considered. Legend: Light blue = Observed iHS; Dark blue = Mixture model; Full red curve = Component 1 of the mixture model; Dashed red curve = Component 2 of the mixture model enriched in selection.

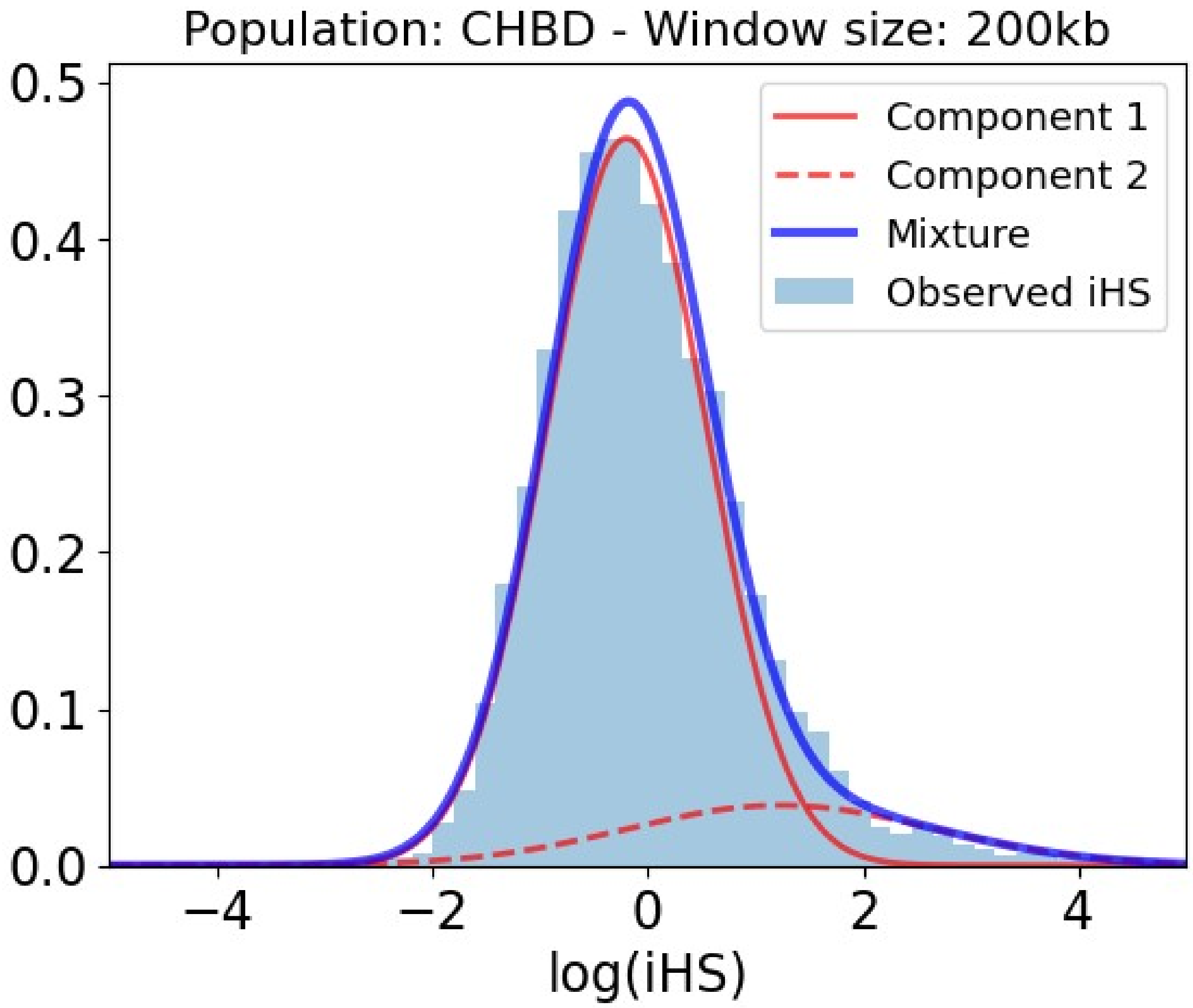


Table S18: Slopes and p-values of the association between iHS and genomic factors for the Han Chinese population in 200kb within the selection-enriched component.

| **Covariate** | **Slope** | **P-value** |
|---|---|---|
| Intercept | -3.774 | 0.000E+00 |
| Number iHS data points | 0.143 | 7.126E-05 |
| Density of conserved elements | 0.115 | 1.193E-01 |
| Recombination rate | -3.444 | 0.000E+00 |
| Number PPIs | 0.046 | 3.054E-01 |
| Regulatory density (ChIP-seq) | 0.170 | 2.119E-01 |
| Distance to VIPs | -0.172 | 1.465E-03 |
| Gene number | -0.001 | 9.979E-01 |
| Coding density | 0.251 | 8.256E-03 |
| Gene length | 0.196 | 4.565E-05 |
| Regulatory density in immune cells (ChIP-seq) | -0.429 | 5.853E-04 |

| **Covariate** | **Slope** | **P-value** |
|---|---|---|
| Gene expression | -0.453 | 4.496E-05 |
| Gene expression in testis | 0.207 | 3.392E-03 |
| Gene expression in immune cells | 0.623 | 9.617E-10 |
| Regulatory density in testis (ChIP-seq) | 0.112 | 2.204E-01 |
| Regulatory density (DNaseI) | -0.283 | 9.546E-02 |
| GC-content | -0.020 | 8.817E-01 |

##### *Han Chinese 500kb*

Figure S19: Mixture of Gaussian distributions fitting observed iHS (500kb windows) for Han Chinese. The figure shows the two Gaussian distributions, component 1 and 2, being the latter enriched in positive selection. In that component, iHS linearly depends on the genomic factors considered. Legend: Light blue = Observed iHS; Dark blue = Mixture model; Full red curve = Component 1 of the mixture model; Dashed red curve = Component 2 of the mixture model enriched in selection.

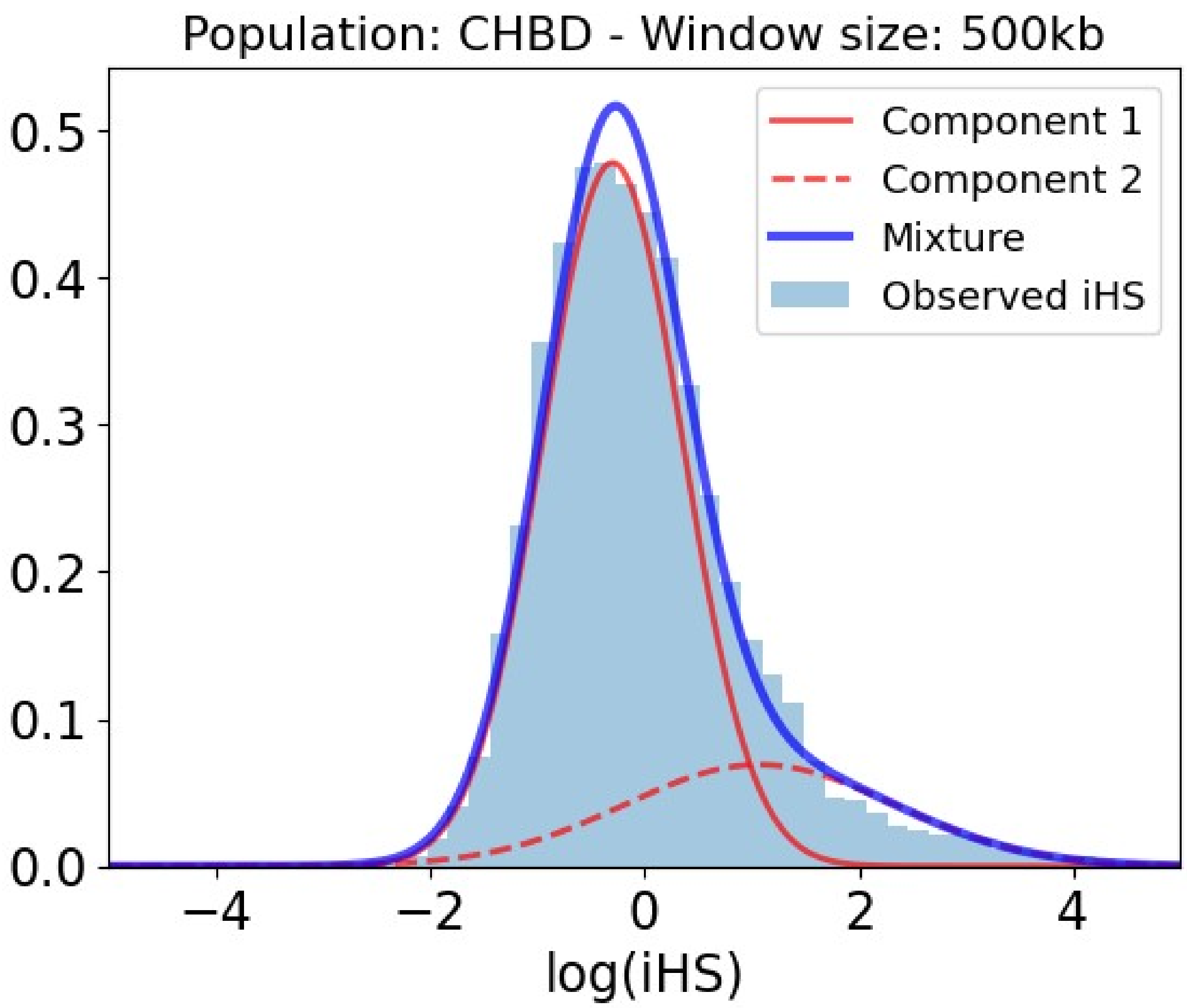


Table S19: Slopes and p-values of the association between iHS and genomic factors for the Han Chinese population in 500kb within the selection-enriched component.

| Covariate | Slope | P-value |
|---|---|---|
| Intercept | -2.146 | 0.000E+00 |
| Number iHS data points | 0.141 | 5.436E-07 |
| Density of conserved elements | 0.196 | 2.236E-04 |
| Recombination rate | -2.191 | 0.000E+00 |
| Number PPIs | -0.044 | 2.385E-01 |
| Regulatory density (ChIP-seq) | 0.294 | 1.344E-02 |
| Distance to VIPs | -0.156 | 2.071E-04 |
| Gene number | -0.128 | 1.567E-01 |
| Coding density | 0.201 | 3.041E-02 |
| Gene length | 0.070 | 7.385E-02 |
| Regulatory density in immune cells (ChIP-seq) | -0.158 | 9.246E-02 |

| Covariate | Slope | P-value |
| --- | --- | --- |
| Gene expression | -0.314 | 5.479E-05 |
| Gene expression in testis | 0.117 | 2.815E-02 |
| Gene expression in immune cells | 0.372 | 2.474E-07 |
| Regulatory density in testis (ChIP-seq) | -0.123 | 5.385E-02 |
| Regulatory density (DNaseI) | -0.415 | 4.798E-03 |
| GC-content | 0.341 | 4.762E-03 |

#### *Han Chinese 1000kb*

Figure S20: Mixture of Gaussian distributions fitting observed iHS (1000kb windows) for Han Chinese. The figure shows the two Gaussian distributions, component 1 and 2, being the latter enriched in positive selection. In that component, iHS linearly depends on the genomic factors considered. Legend: Light blue = Observed iHS; Dark blue = Mixture model; Full red curve = Component 1 of the mixture model; Dashed red curve = Component 2 of the mixture model enriched in selection.

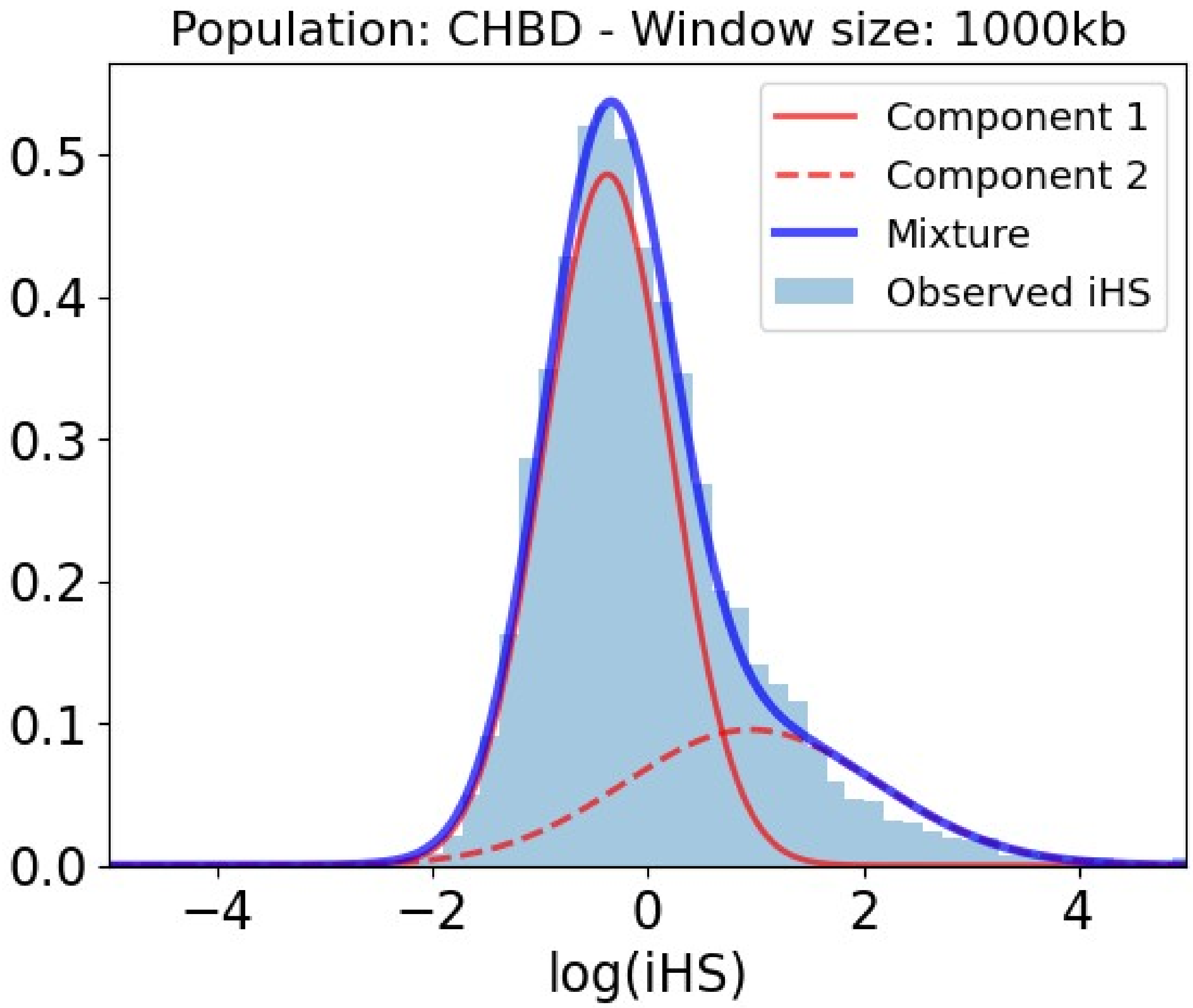


Table S20: Slopes and p-values of the association between iHS and genomic factors for the Han Chinese population in 1000kb within the selection-enriched component.

| **Covariate** | **Slope** | **P-value** |
|---|---|---|
| Intercept | -1.778 | 0.000E+00 |
| Number iHS data points | 0.157 | 7.292E-08 |
| Density of conserved elements | 0.311 | 4.199E-10 |
| Recombination rate | -2.510 | 0.000E+00 |
| Number PPIs | -0.088 | 1.925E-02 |
| Regulatory density (ChIP-seq) | 0.490 | 3.921E-05 |
| Distance to VIPs | -0.142 | 5.348E-04 |
| Gene number | -0.460 | 6.230E-04 |
| Coding density | 0.396 | 1.928E-03 |
| Gene length | 0.004 | 9.125E-01 |
| Regulatory density in immune cells (ChIP-seq) | -0.160 | 1.258E-01 |

| Covariate | Slope | P-value |
|---|---|---|
| Gene expression | -0.245 | 1.135E-03 |
| Gene expression in testis | 0.065 | 2.056E-01 |
| Gene expression in immune cells | 0.307 | 9.507E-06 |
| Regulatory density in testis (ChIP-seq) | -0.393 | 1.650E-09 |
| Regulatory density (DNaseI) | -0.838 | 1.660E-08 |
| GC-content | 1.171 | 0.000E+00 |

### Peruvians

#### *Peruvians 50kb*

Figure S21: Mixture of Gaussian distributions fitting observed iHS (50kb windows) for Peruvians. The figure shows the two Gaussian distributions, component 1 and 2, being the latter enriched in positive selection. In that component, iHS linearly depends on the genomic factors considered. Legend: Light blue = Observed iHS; Dark blue = Mixture model; Full red curve = Component 1 of the mixture model; Dashed red curve = Component 2 of the mixture model enriched in selection.

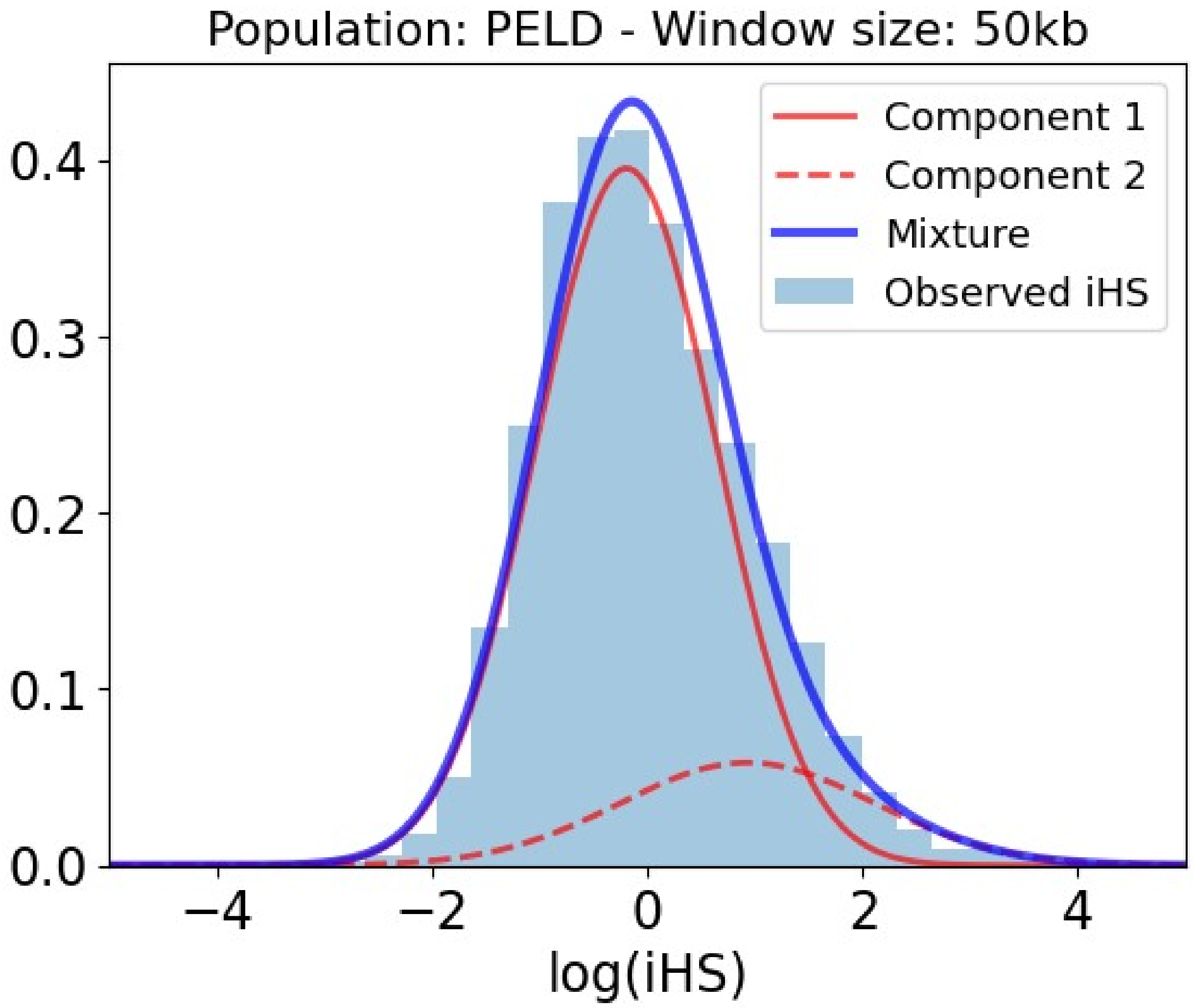


Table S21: Slopes and p-values of the association between iHS and genomic factors for the Peruvians population in 50kb within the selection-enriched component.

| Covariate | Slope | P-value |
|---|---|---|
| Intercept | -36.485 | 0.000E+00 |
| Number iHS data points | 0.648 | 6.796E-09 |
| Density of conserved elements | -0.443 | 7.655E-03 |
| Recombination rate | -58.880 | 0.000E+00 |
| Number PPIs | 0.229 | 7.376E-03 |
| Regulatory density (ChIP-seq) | -0.059 | 7.992E-01 |
| Distance to VIPs | -0.219 | 6.141E-03 |
| Gene number | -0.077 | 3.939E-01 |
| Coding density | 0.068 | 6.490E-01 |
| Gene length | 0.423 | 1.041E-03 |
| Regulatory density in immune cells (ChIP-seq) | 0.062 | 7.900E-01 |

| Covariate | Slope | P-value |
|---|---|---|
| Gene expression | -0.632 | 5.121E-03 |
| Gene expression in testis | 0.449 | 8.541E-04 |
| Gene expression in immune cells | 0.535 | 1.707E-02 |
| Regulatory density in testis (ChIP-seq) | 0.243 | 7.025E-02 |
| Regulatory density (DNaseI) | 0.391 | 1.777E-01 |
| GC-content | -0.023 | 9.263E-01 |

#### *Peruvians 100kb*

Figure S22: Mixture of Gaussian distributions fitting observed iHS (100kb windows) for Peruvians. The figure shows the two Gaussian distributions, component 1 and 2, being the latter enriched in positive selection. In that component, iHS linearly depends on the genomic factors considered. Legend: Light blue = Observed iHS; Dark blue = Mixture model; Full red curve = Component 1 of the mixture model; Dashed red curve = Component 2 of the mixture model enriched in selection.

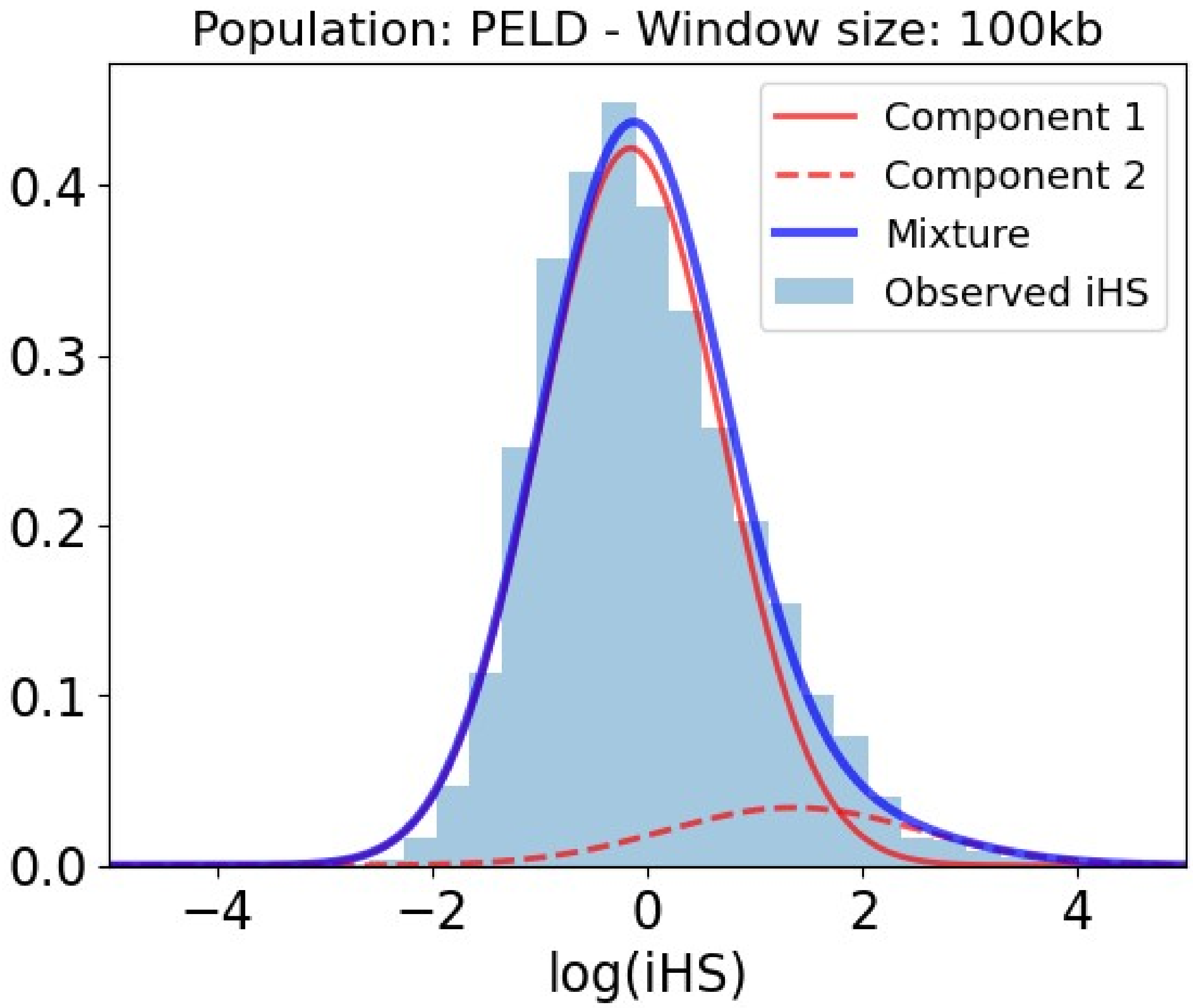


Table S22: Slopes and p-values of the association between iHS and genomic factors for the Peruvians population in 100kb within the selection-enriched component.

| Covariate | Slope | P-value |
|---|---|---|
| Intercept | -46.010 | 0.000E+00 |
| Number iHS data points | 1.430 | 0.000E+00 |
| Density of conserved elements | -0.050 | 7.650E-01 |
| Recombination rate | -59.143 | 0.000E+00 |
| Number PPIs | 0.123 | 1.741E-01 |
| Regulatory density (ChIP-seq) | -0.368 | 1.963E-01 |
| Distance to VIPs | -0.183 | 4.899E-02 |
| Gene number | 0.247 | 1.160E-01 |
| Coding density | -0.392 | 2.096E-02 |
| Gene length | 0.318 | 3.375E-03 |
| Regulatory density in immune cells (ChIP-seq) | 0.135 | 6.664E-01 |

| Covariate | Slope | P-value |
|---|---|---|
| Gene expression | -0.128 | 5.601E-01 |
| Gene expression in testis | 0.313 | 3.456E-02 |
| Gene expression in immune cells | 0.340 | 1.265E-01 |
| Regulatory density in testis (ChIP-seq) | 0.190 | 1.681E-01 |
| Regulatory density (DNaseI) | -0.102 | 7.513E-01 |
| GC-content | 0.294 | 2.438E-01 |

#### *Peruvians 200kb*

Figure S23: Mixture of Gaussian distributions fitting observed iHS (200kb windows) for Peruvians. The figure shows the two Gaussian distributions, component 1 and 2, being the latter enriched in positive selection. In that component, iHS linearly depends on the genomic factors considered. Legend: Light blue = Observed iHS; Dark blue = Mixture model; Full red curve = Component 1 of the mixture model; Dashed red curve = Component 2 of the mixture model enriched in selection.

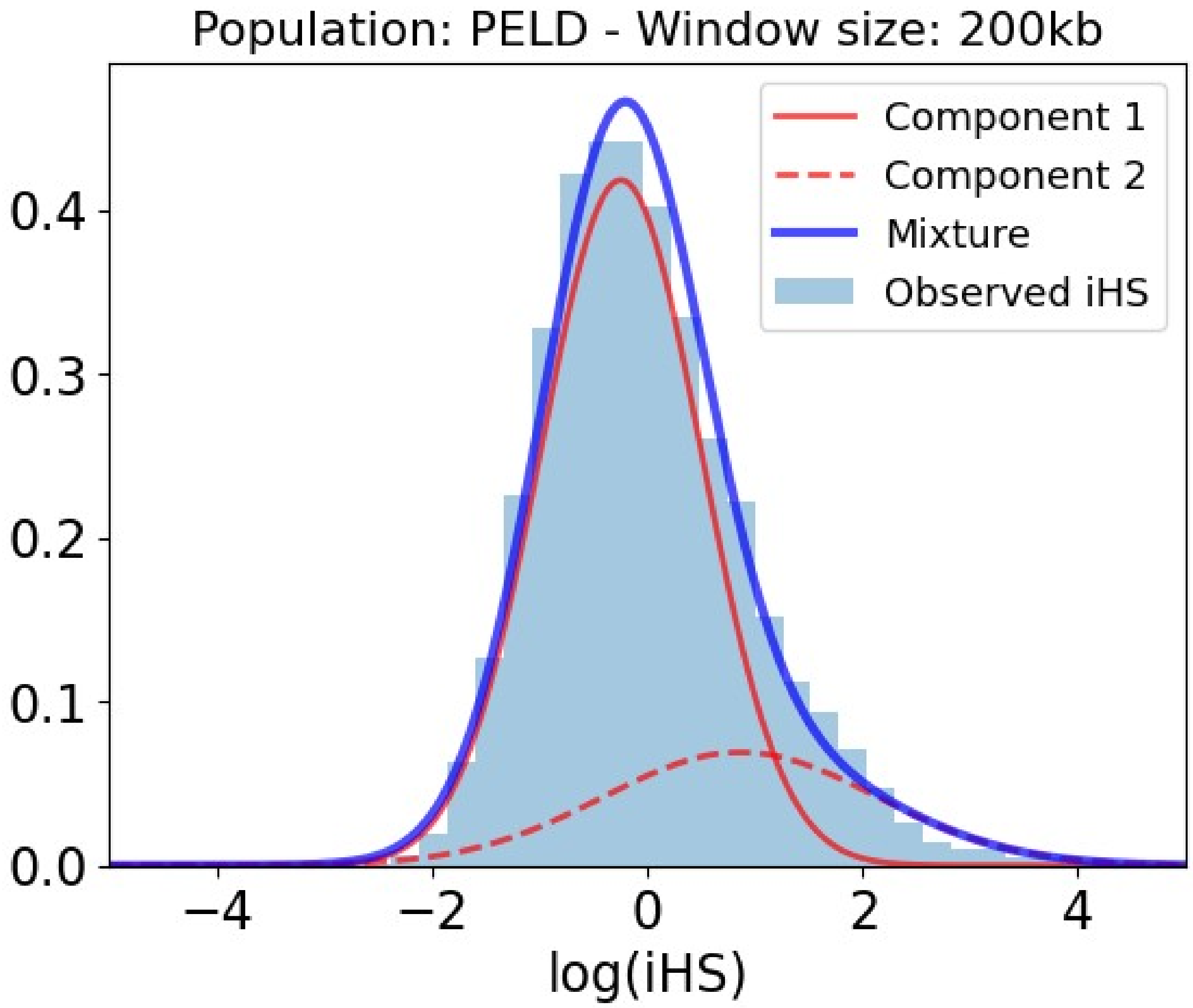


Table S23: Slopes and p-values of the association between iHS and genomic factors for the Peruvians population in 200kb within the selection-enriched component.

| **Covariate** | **Slope** | **P-value** |
|---|---|---|
| Intercept | -2.553 | 0.000E+00 |
| Number iHS data points | 0.077 | 1.952E-02 |
| Density of conserved elements | -0.216 | 2.002E-03 |
| Recombination rate | -2.963 | 0.000E+00 |
| Number PPIs | 0.030 | 4.892E-01 |
| Regulatory density (ChIP-seq) | 0.134 | 2.563E-01 |
| Distance to VIPs | -0.231 | 3.345E-05 |
| Gene number | 0.024 | 7.596E-01 |
| Coding density | 0.163 | 4.123E-02 |
| Gene length | 0.135 | 6.324E-03 |
| Regulatory density in immune cells (ChIP-seq) | -0.122 | 2.385E-01 |

| Covariate | Slope | P-value |
|---|---|---|
| Gene expression | -0.311 | 2.186E-03 |
| Gene expression in testis | 0.090 | 1.676E-01 |
| Gene expression in immune cells | 0.442 | 2.461E-06 |
| Regulatory density in testis (ChIP-seq) | 0.139 | 1.327E-02 |
| Regulatory density (DNaseI) | -0.056 | 6.978E-01 |
| GC-content | 0.069 | 5.587E-01 |

#### *Peruvians 500kb*

Figure S24: Mixture of Gaussian distributions fitting observed iHS (500kb windows) for Peruvians. The figure shows the two Gaussian distributions, component 1 and 2, being the latter enriched in positive selection. In that component, iHS linearly depends on the genomic factors considered. Legend: Light blue = Observed iHS; Dark blue = Mixture model; Full red curve = Component 1 of the mixture model; Dashed red curve = Component 2 of the mixture model enriched in selection.

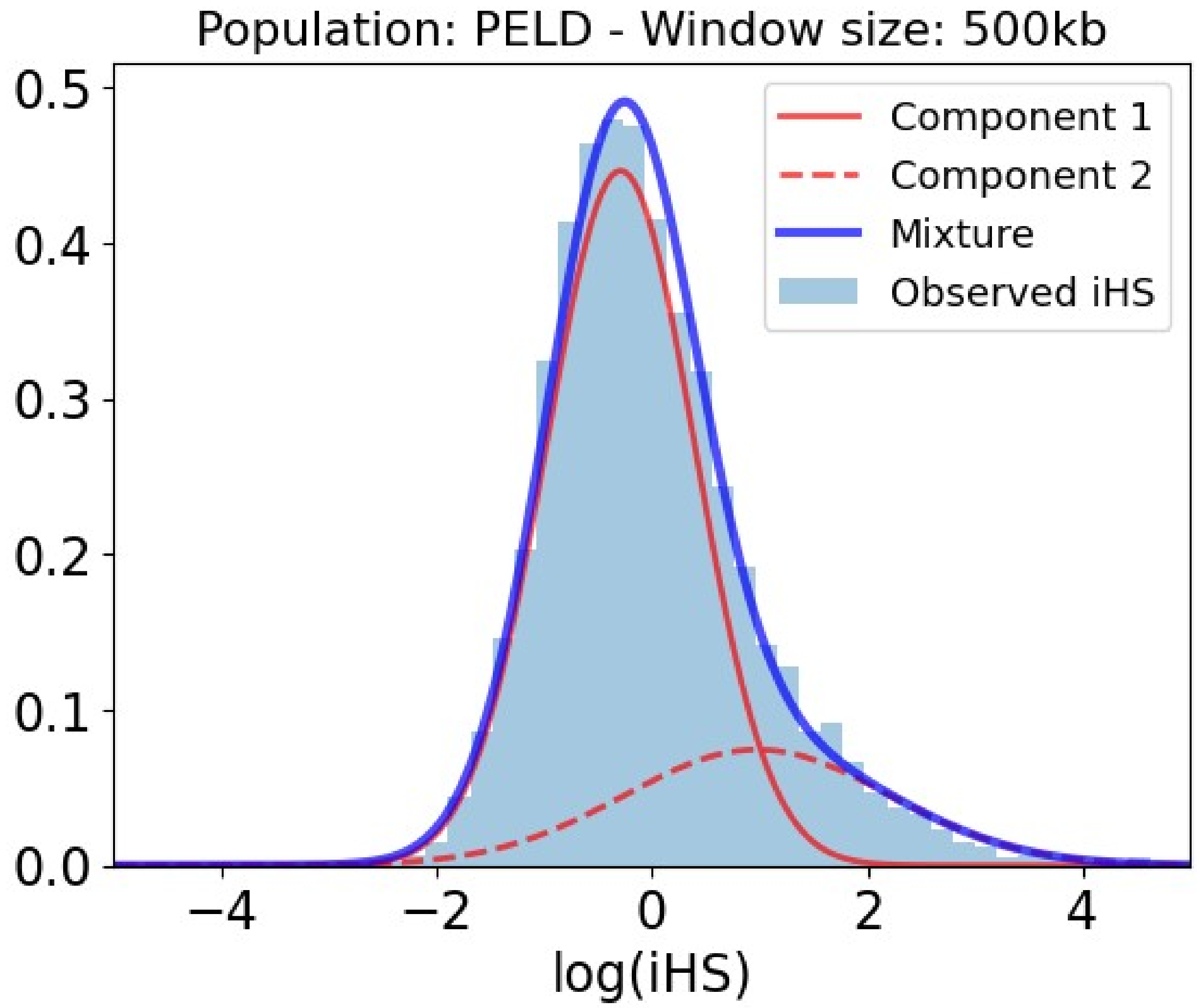


Table S24: Slopes and p-values of the association between iHS and genomic factors for the Peruvians population in 500kb within the selection-enriched component.

| **Covariate** | **Slope** | **P-value** |
|---|---|---|
| Intercept | -1.970 | 0.000E+00 |
| Number iHS data points | 0.093 | 1.343E-03 |
| Density of conserved elements | -0.137 | 1.499E-02 |
| Recombination rate | -2.033 | 0.000E+00 |
| Number PPIs | -0.040 | 3.115E-01 |
| Regulatory density (ChIP-seq) | -0.044 | 7.231E-01 |
| Distance to VIPs | -0.295 | 2.812E-07 |
| Gene number | -0.072 | 3.523E-01 |
| Coding density | 0.216 | 8.404E-03 |
| Gene length | 0.077 | 8.007E-02 |
| Regulatory density in immune cells (ChIP-seq) | 0.221 | 1.628E-02 |

| Covariate | Slope | P-value |
|---|---|---|
| Gene expression | -0.287 | 4.502E-04 |
| Gene expression in testis | 0.130 | 1.819E-02 |
| Gene expression in immune cells | 0.284 | 1.561E-04 |
| Regulatory density in testis (ChIP-seq) | 0.017 | 7.460E-01 |
| Regulatory density (DNaseI) | -0.208 | 1.577E-01 |
| GC-content | 0.180 | 1.323E-01 |

#### *Peruvians 1000kb*

Figure S25: Mixture of Gaussian distributions fitting observed iHS (1000kb windows) for Peruvians. The figure shows the two Gaussian distributions, component 1 and 2, being the latter enriched in positive selection. In that component, iHS linearly depends on the genomic factors considered. Legend: Light blue = Observed iHS; Dark blue = Mixture model; Full red curve = Component 1 of the mixture model; Dashed red curve = Component 2 of the mixture model enriched in selection.

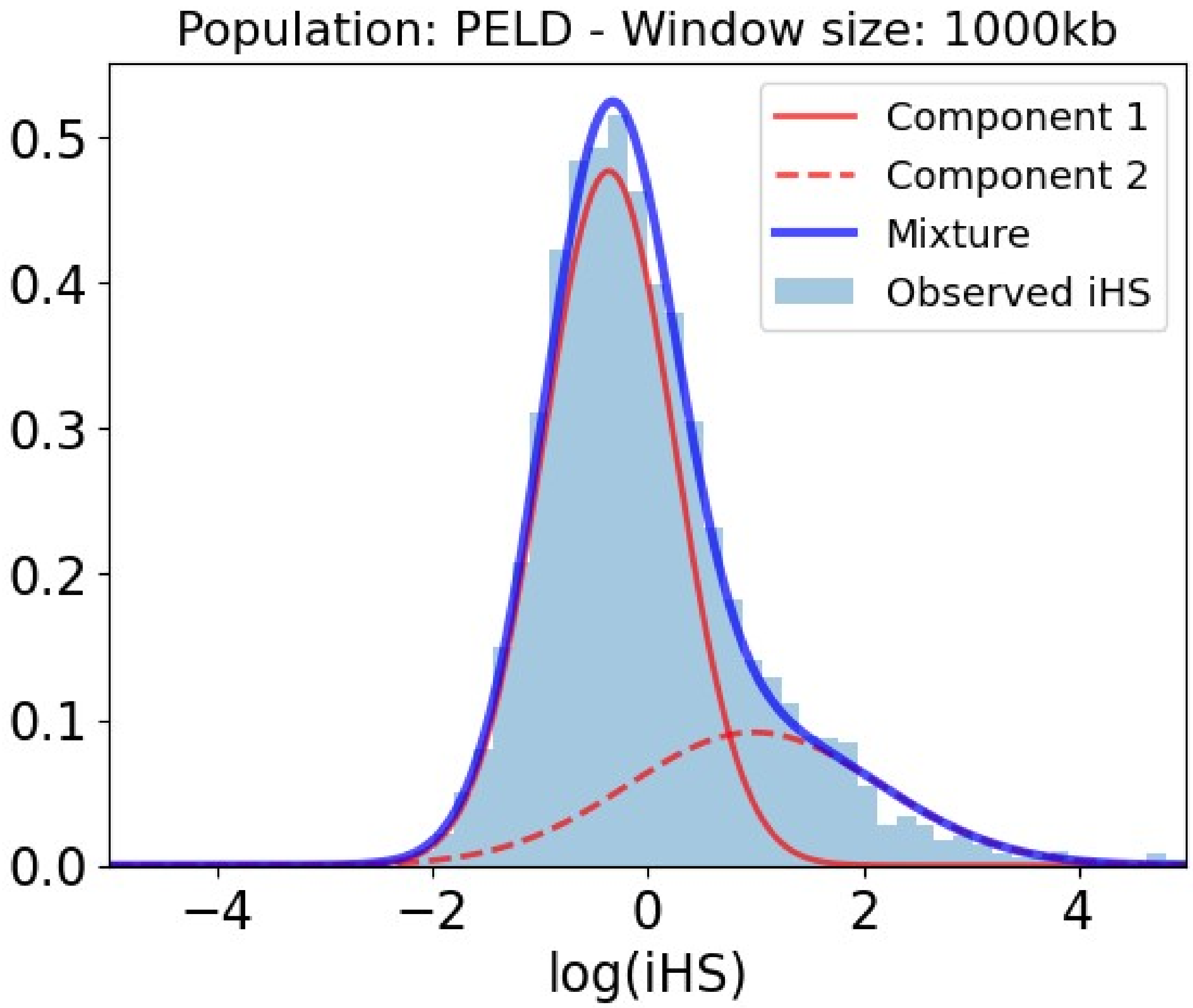


Table S25: Slopes and p-values of the association between iHS and genomic factors for the Peruvians population in 1000kb within the selection-enriched component.

| **Covariate** | **Slope** | **P-value** |
|---|---|---|
| Intercept | -1.548 | 0.000E+00 |
| Number iHS data points | 0.081 | 1.506E-03 |
| Density of conserved elements | 0.057 | 2.433E-01 |
| Recombination rate | -1.738 | 0.000E+00 |
| Number PPIs | -0.037 | 3.003E-01 |
| Regulatory density (ChIP-seq) | 0.327 | 7.640E-03 |
| Distance to VIPs | -0.178 | 2.765E-05 |
| Gene number | -0.021 | 7.960E-01 |
| Coding density | 0.202 | 2.729E-02 |
| Gene length | 0.023 | 5.465E-01 |
| Regulatory density in immune cells (ChIP-seq) | 0.146 | 1.317E-01 |

| Covariate | Slope | P-value |
|---|---|---|
| Gene expression | -0.261 | 2.560E-04 |
| Gene expression in testis | 0.053 | 2.606E-01 |
| Gene expression in immune cells | 0.251 | 1.561E-04 |
| Regulatory density in testis (ChIP-seq) | -0.107 | 4.757E-02 |
| Regulatory density (DNaseI) | -0.551 | 8.267E-05 |
| GC-content | 0.351 | 2.628E-03 |
