## Supplemental Results S3 for "Mixture Density Regression reveals frequent recent adaptation in the human genome"

### Yoruba

#### *Yoruba 1000kb: Removing recombination rate.*

Figure S1: Mixture of Gaussian distributions fitting observed iHS (1000kb windows) for Yoruba. The figure shows the two Gaussian distributions, component 1 and 2, being the latter enriched in positive selection. In that component, iHS linearly depends on the genomic factors considered. Legend: Light blue = Observed iHS; Dark blue = Mixture model; Full red curve = Component 1 of the mixture model; Dashed red curve = Component 2 of the mixture model enriched in selection.

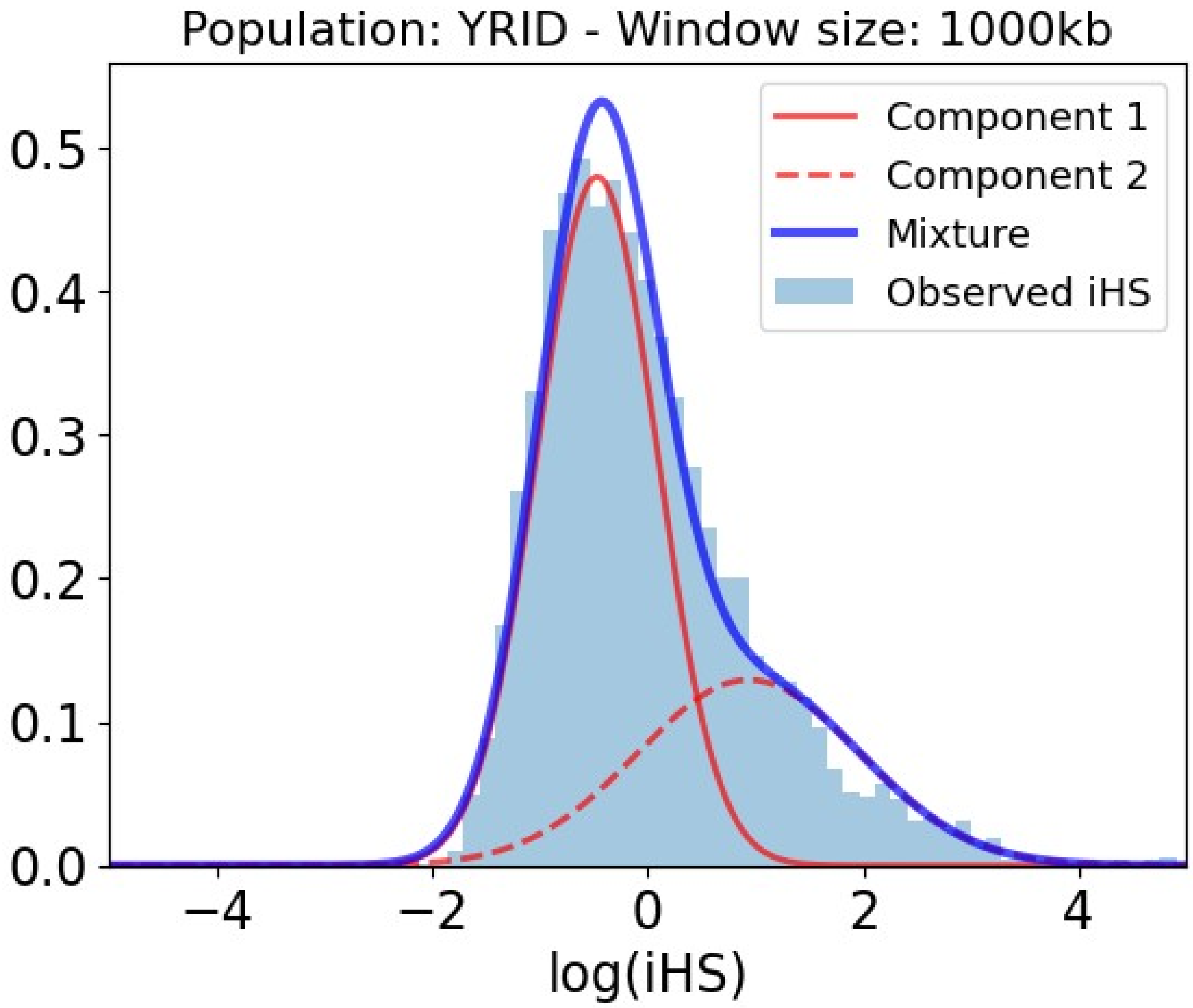


Table S1: Slopes and p-values of the association between iHS and genomic factors for the Yoruba population in 1000kb within the selection-enriched component.

| Covariate | Slope | P-value |
|---|---|---|
| Intercept | -0.863 | 0.000E+00 |

| **Covariate** | **Slope** | **P-value** |
|---|---|---|
| Number iHS data points | -0.872 | 0.000E+00 |
| Regulatory density (ChIP-seq) | -0.305 | 6.125E-04 |
| Regulatory density in immune cells (ChIP-seq) | 0.271 | 4.487E-04 |
| Regulatory density in testis (ChIP-seq) | -0.523 | 0.000E+00 |
| Coding density | 0.414 | 1.209E-07 |
| Density of conserved elements | 0.095 | 1.462E-02 |
| Gene expression | -0.138 | 1.354E-02 |
| GC-content | -0.287 | 1.224E-03 |
| Gene length | -0.039 | 2.278E-01 |
| Gene number | -0.063 | 4.293E-01 |
| Gene expression in immune cells | 0.260 | 4.759E-07 |
| Number PPIs | -0.078 | 6.274E-03 |
| Regulatory density (DNaseI) | -0.127 | 2.198E-01 |
| Gene expression in testis | 0.075 | 5.629E-02 |
| Distance to VIPs | -0.181 | 1.460E-07 |

#### *Yoruba 1000kb: Removing GC-content.*

Figure S2: Mixture of Gaussian distributions fitting observed iHS (1000kb windows) for Yoruba. The figure shows the two Gaussian distributions, component 1 and 2, being the latter enriched in positive selection. In that component, iHS linearly depends on the genomic factors considered. Legend: Light blue = Observed iHS; Dark blue = Mixture model; Full red curve = Component 1 of the mixture model; Dashed red curve = Component 2 of the mixture model enriched in selection.

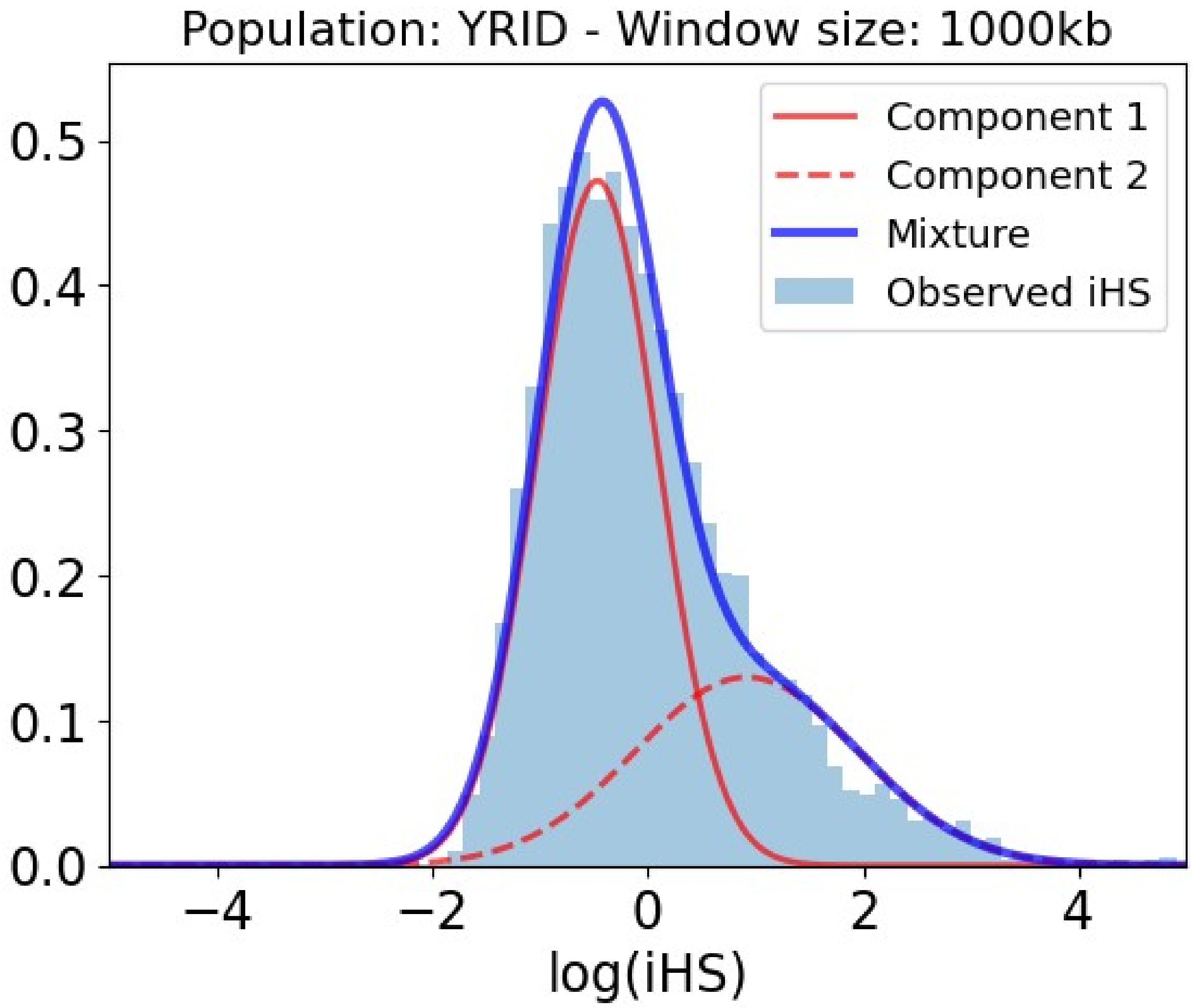


Table S2: Slopes and p-values of the association between iHS and genomic factors for the Yoruba population in 1000kb within the selection-enriched component.

| **Covariate** | **Slope** | **P-value** |
|---|---|---|
| Intercept | -1.420 | 0.000E+00 |
| Number iHS data points | -0.027 | 3.780E-01 |
| Regulatory density (ChIP-seq) | -0.222 | 7.334E-02 |
| Regulatory density in immune cells (ChIP-seq) | 0.072 | 4.520E-01 |
| Regulatory density in testis (ChIP-seq) | -0.765 | 0.000E+00 |
| Coding density | 0.225 | 1.321E-02 |
| Density of conserved elements | 0.100 | 2.620E-02 |
| Gene expression | -0.148 | 2.805E-02 |
| Gene length | -0.061 | 1.028E-01 |
| Gene number | -0.039 | 6.654E-01 |

| Covariate | Slope | P-value |
|---|---|---|
| Gene expression in immune cells | 0.271 | 9.964E-06 |
| Number PPIs | -0.076 | 2.607E-02 |
| Recombination rate | -2.390 | 0.000E+00 |
| Regulatory density (DNaseI) | 0.444 | 1.775E-07 |
| Gene expression in testis | 0.027 | 5.736E-01 |
| Distance to VIPs | -0.177 | 1.581E-06 |

#### *Yoruba 1000kb: Removing regulatory density according to DNAseI hypersensitivity and coding density.*

Figure S3: Mixture of Gaussian distributions fitting observed iHS (1000kb windows) for Yoruba. The figure shows the two Gaussian distributions, component 1 and 2, being the latter enriched in positive selection. In that component, iHS linearly depends on the genomic factors considered. Legend: Light blue = Observed iHS; Dark blue = Mixture model; Full red curve = Component 1 of the mixture model; Dashed red curve = Component 2 of the mixture model enriched in selection.

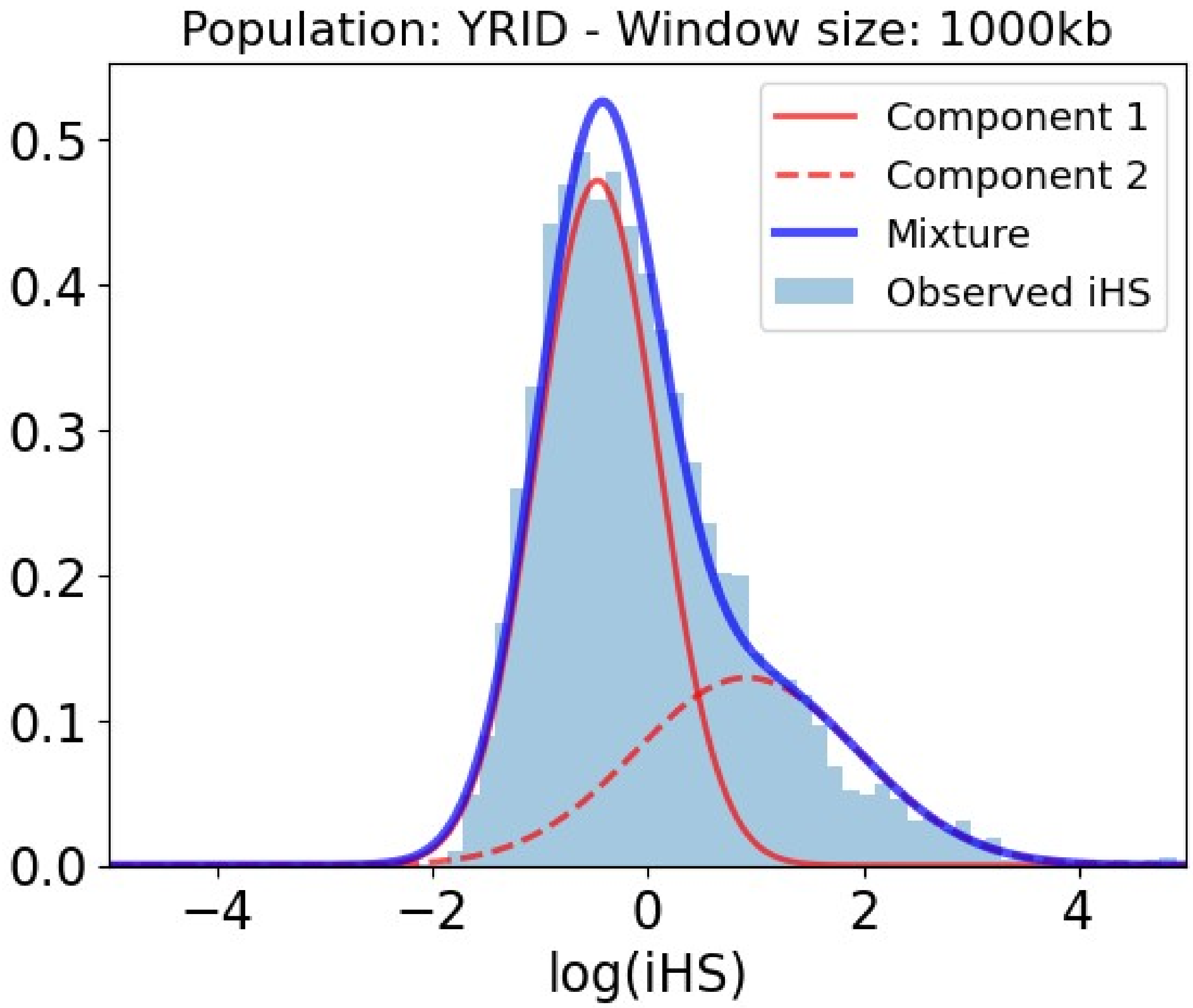


Table S3: Slopes and p-values of the association between iHS and genomic factors for the Yoruba population in 1000kb within the selection-enriched component.

| Covariate | Slope | P-value |
|---|---|---|
| Intercept | -1.433 | 0.000E+00 |
| Number iHS data points | -0.003 | 9.285E-01 |
| Regulatory density (ChIP-seq) | -0.175 | 1.029E-01 |
| Regulatory density in immune cells (ChIP-seq) | 0.037 | 6.950E-01 |
| Regulatory density in testis (ChIP-seq) | -0.783 | 0.000E+00 |
| Density of conserved elements | 0.183 | 2.482E-05 |
| Gene expression | -0.134 | 4.510E-02 |
| GC-content | 0.543 | 2.398E-14 |
| Gene length | -0.058 | 1.195E-01 |
| Gene number | 0.042 | 4.417E-01 |

| Covariate | Slope | P-value |
|---|---|---|
| Gene expression in immune cells | 0.264 | 1.699E-05 |
| Number PPIs | -0.073 | 3.572E-02 |
| Recombination rate | -2.451 | 0.000E+00 |
| Gene expression in testis | 0.006 | 9.036E-01 |
| Distance to VIPs | -0.169 | 4.627E-06 |

#### *Yoruba 1000kb: Removing regulatory density according to DNAseI sensitivity, coding density, regulatory density according to ChIP-seq experiments across multiple tissues and specifically in immune cells, gene number and gene length.*

Figure S4: Mixture of Gaussian distributions fitting observed iHS (1000kb windows) for Yoruba. The figure shows the two Gaussian distributions, component 1 and 2, being the latter enriched in positive selection. In that component, iHS linearly depends on the genomic factors considered. Legend: Light blue = Observed iHS; Dark blue = Mixture model; Full red curve = Component 1 of the mixture model; Dashed red curve = Component 2 of the mixture model enriched in selection.

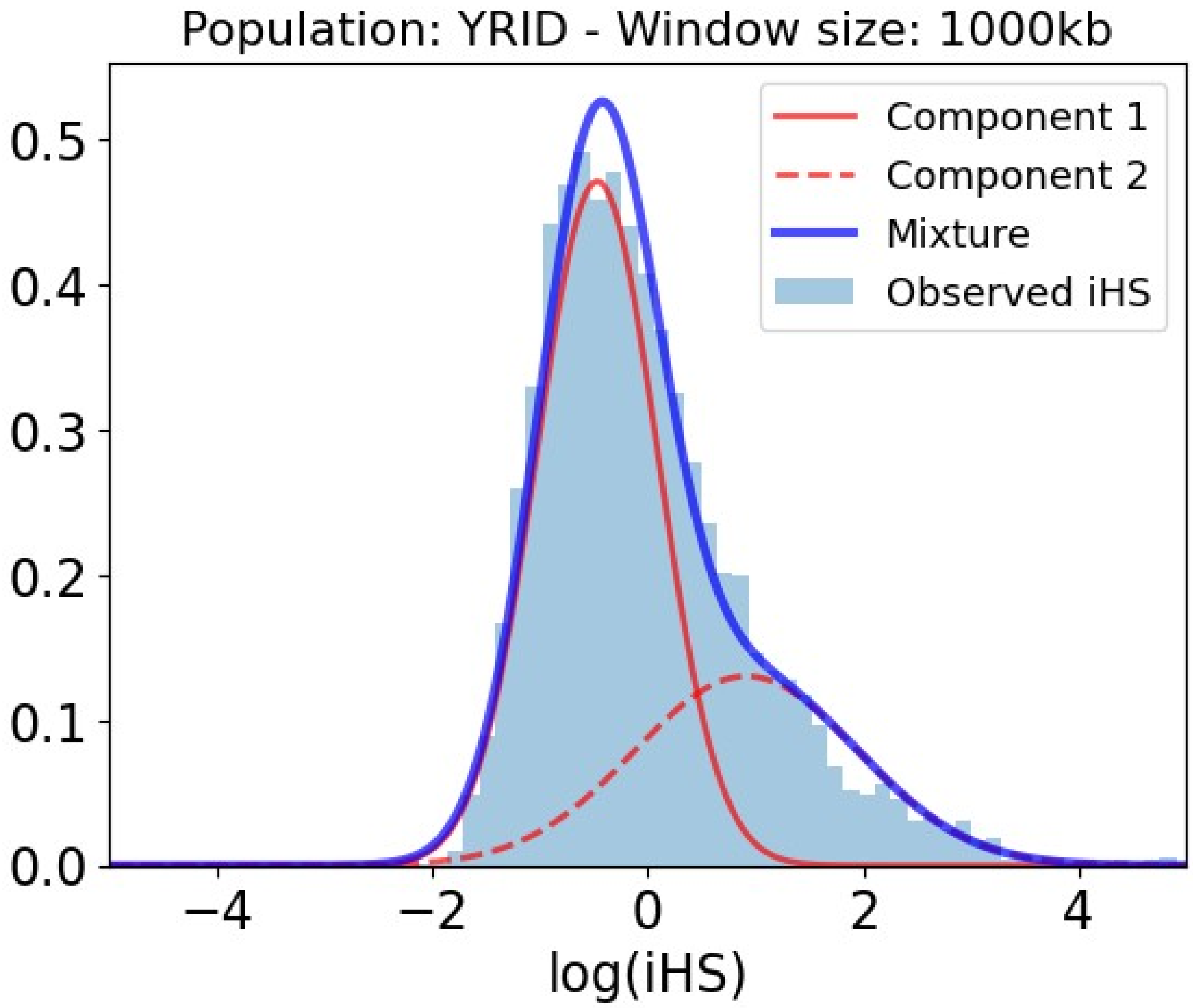


Table S4: Slopes and p-values of the association between iHS and genomic factors for the Yoruba population in 1000kb within the selection-enriched component.

| **Covariate** | **Slope** | **P-value** |
|---|---|---|
| Intercept | -1.412 | 0.000E+00 |
| Number iHS data points | -0.003 | 9.021E-01 |
| Regulatory density in testis (ChIP-seq) | -0.792 | 0.000E+00 |
| Density of conserved elements | 0.170 | 2.757E-05 |
| Gene expression | -0.154 | 1.781E-02 |
| GC-content | 0.492 | 0.000E+00 |
| Gene expression in immune cells | 0.274 | 3.358E-06 |
| Number PPIs | -0.080 | 1.953E-02 |
| Recombination rate | -2.466 | 0.000E+00 |
| Gene expression in testis | 0.002 | 9.685E-01 |

| Covariate | Slope | P-value |
|---|---|---|
| Distance to VIPs | -0.176 | 7.015E-07 |

##### *Yoruba 1000kb: Removing the density of conserved elements.*

Figure S5: Mixture of Gaussian distributions fitting observed iHS (1000kb windows) for Yoruba. The figure shows the two Gaussian distributions, component 1 and 2, being the latter enriched in positive selection. In that component, iHS linearly depends on the genomic factors considered. Legend: Light blue = Observed iHS; Dark blue = Mixture model; Full red curve = Component 1 of the mixture model; Dashed red curve = Component 2 of the mixture model enriched in selection.

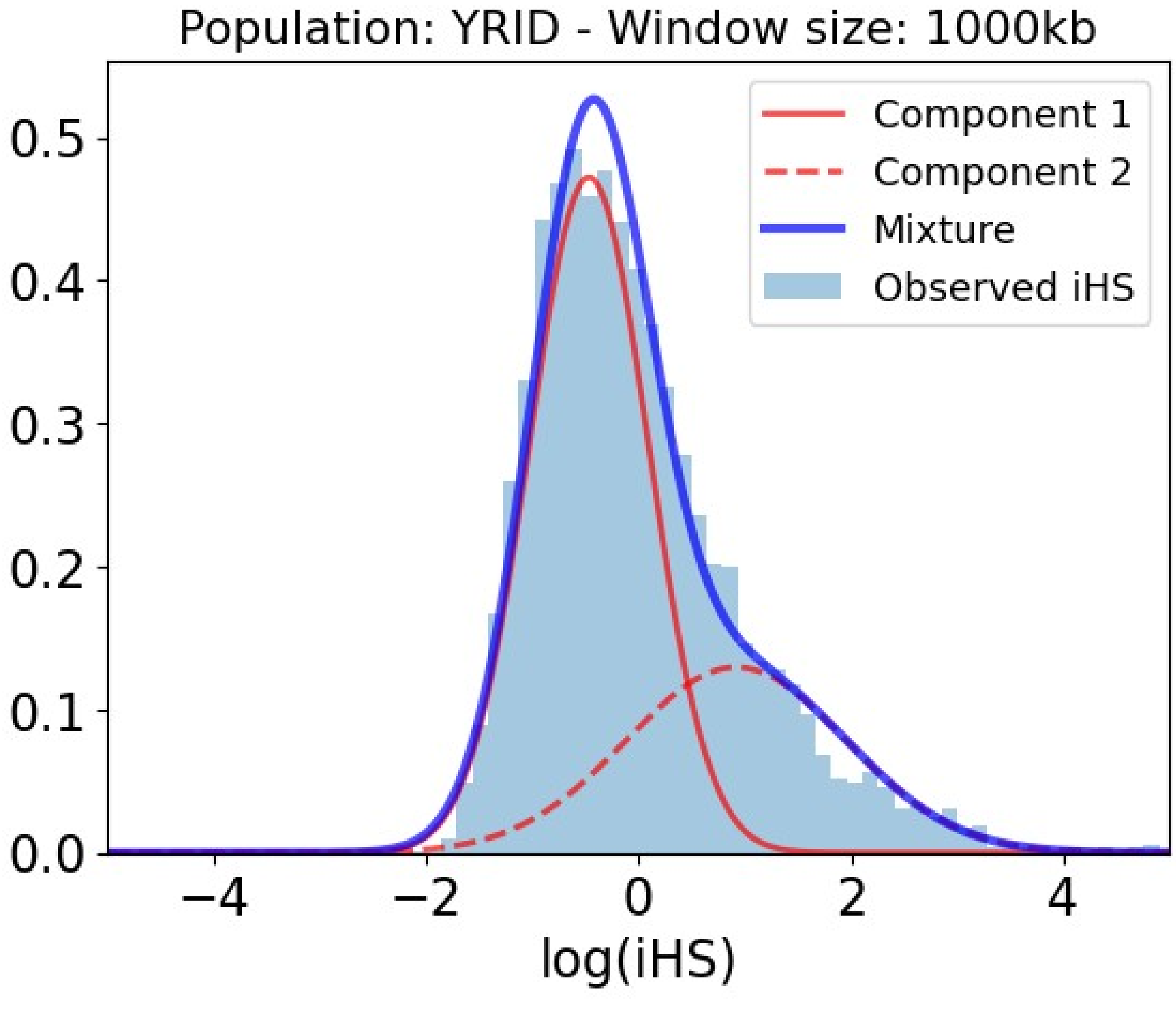


Table S5: Slopes and p-values of the association between iHS and genomic factors for the Yoruba population in 1000kb within the selection-enriched component.

| **Covariate** | **Slope** | **P-value** |
|---|---|---|
| Intercept | -1.411 | 0.000E+00 |
| Number iHS data points | -0.033 | 2.703E-01 |
| Regulatory density (ChIP-seq) | -0.135 | 2.741E-01 |
| Regulatory density in immune cells (ChIP-seq) | 0.095 | 3.160E-01 |
| Regulatory density in testis (ChIP-seq) | -0.763 | 0.000E+00 |
| Coding density | 0.259 | 2.347E-03 |
| Gene expression | -0.126 | 5.905E-02 |
| GC-content | 0.447 | 4.494E-05 |
| Gene length | -0.046 | 2.151E-01 |
| Gene number | -0.120 | 1.822E-01 |
| Gene expression in immune cells | 0.249 | 4.291E-05 |
| Number PPIs | -0.068 | 4.944E-02 |
| Recombination rate | -2.425 | 0.000E+00 |
| Regulatory density (DNaseI) | 0.023 | 8.815E-01 |
| Gene expression in testis | 0.010 | 8.351E-01 |
| Distance to VIPs | -0.174 | 2.795E-06 |

#### *Yoruba 1000kb: Removing the density of conserved elements and GC-content.*

Figure S6: Mixture of Gaussian distributions fitting observed iHS (1000kb windows) for Yoruba. The figure shows the two Gaussian distributions, component 1 and 2, being the latter enriched in positive selection. In that component, iHS linearly depends on the genomic factors considered. Legend: Light blue = Observed iHS; Dark blue = Mixture model; Full red curve = Component 1 of the mixture model; Dashed red curve = Component 2 of the mixture model enriched in selection.

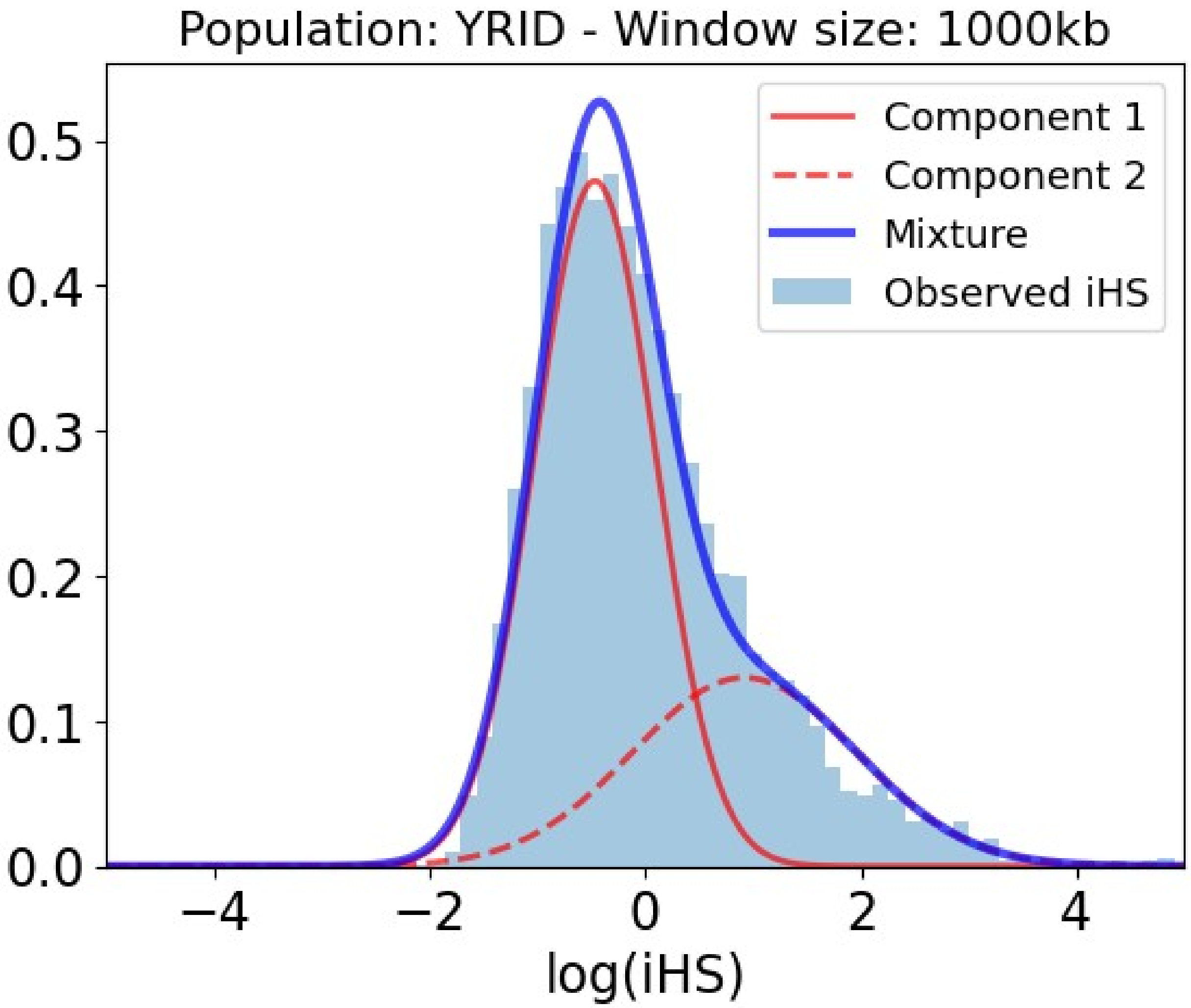


Table S6: Slopes and p-values of the association between iHS and genomic factors for the Yoruba population in 1000kb within the selection-enriched component.

| **Covariate** | **Slope** | **P-value** |
|---|---|---|
| Intercept | -1.409 | 0.000E+00 |
| Number iHS data points | -0.041 | 1.612E-01 |
| Regulatory density (ChIP-seq) | -0.202 | 9.854E-02 |
| Regulatory density in immune cells (ChIP-seq) | 0.099 | 2.894E-01 |
| Regulatory density in testis (ChIP-seq) | -0.749 | 0.000E+00 |
| Coding density | 0.287 | 7.763E-04 |
| Gene expression | -0.138 | 3.976E-02 |
| Gene length | -0.052 | 1.618E-01 |
| Gene number | -0.070 | 4.440E-01 |
| Gene expression in immune | 0.260 | 2.090E-05 |

| Covariate | Slope | P-value |
|---|---|---|
| cells | | |
| Number PPIs | -0.074 | 3.108E-02 |
| Recombination rate | -2.387 | 0.000E+00 |
| Regulatory density (DNaseI) | 0.427 | 4.012E-07 |
| Gene expression in testis | 0.029 | 5.363E-01 |
| Distance to VIPs | -0.180 | 1.293E-06 |

#### *Yoruba 1000kb: Removing no predictor.*

Figure S7: Mixture of Gaussian distributions fitting observed iHS (1000kb windows) for Yoruba. The figure shows the two Gaussian distributions, component 1 and 2, being the latter enriched in positive selection. In that component, iHS linearly depends on the genomic factors considered. Legend: Light blue = Observed iHS; Dark blue = Mixture model; Full red curve = Component 1 of the mixture model; Dashed red curve = Component 2 of the mixture model enriched in selection.

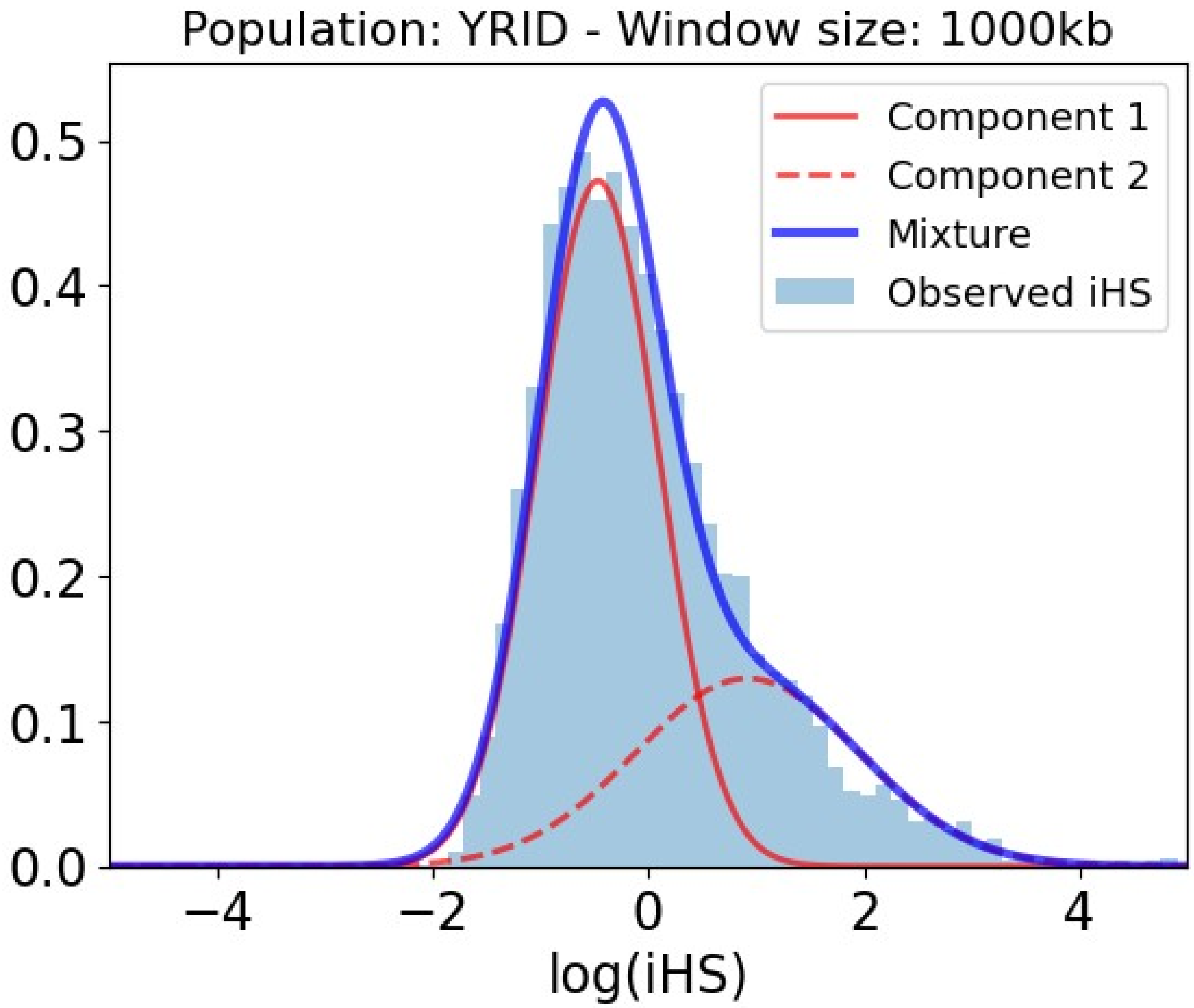


Table S7: Slopes and p-values of the association between iHS and genomic factors for the Yoruba population in 1000kb within the selection-enriched component.

| **Covariate** | **Slope** | **P-value** |
|---|---|---|
| Intercept | -1.426 | 0.000E+00 |
| Number iHS data points | -0.007 | 8.190E-01 |
| Regulatory density (ChIP-seq) | -0.147 | 2.372E-01 |
| Regulatory density in immune cells (ChIP-seq) | 0.047 | 6.230E-01 |
| Regulatory density in testis (ChIP-seq) | -0.795 | 0.000E+00 |
| Coding density | 0.159 | 8.411E-02 |
| Density of conserved elements | 0.159 | 6.397E-04 |
| Gene expression | -0.139 | 3.768E-02 |
| GC-content | 0.550 | 1.339E-06 |
| Gene length | -0.059 | 1.124E-01 |

| Covariate | Slope | P-value |
|---|---|---|
| Gene number | -0.086 | 3.458E-01 |
| Gene expression in immune cells | 0.265 | 1.512E-05 |
| Number PPIs | -0.070 | 4.252E-02 |
| Recombination rate | -2.436 | 0.000E+00 |
| Regulatory density (DNaseI) | -0.047 | 7.138E-01 |
| Gene expression in testis | 0.001 | 9.803E-01 |
| Distance to VIPs | -0.170 | 4.160E-06 |
