## Supplemental Results S4 for "Mixture Density Regression reveals frequent recent adaptation in the human genome"

### iHS, recombination rate and distance to regulatory elements

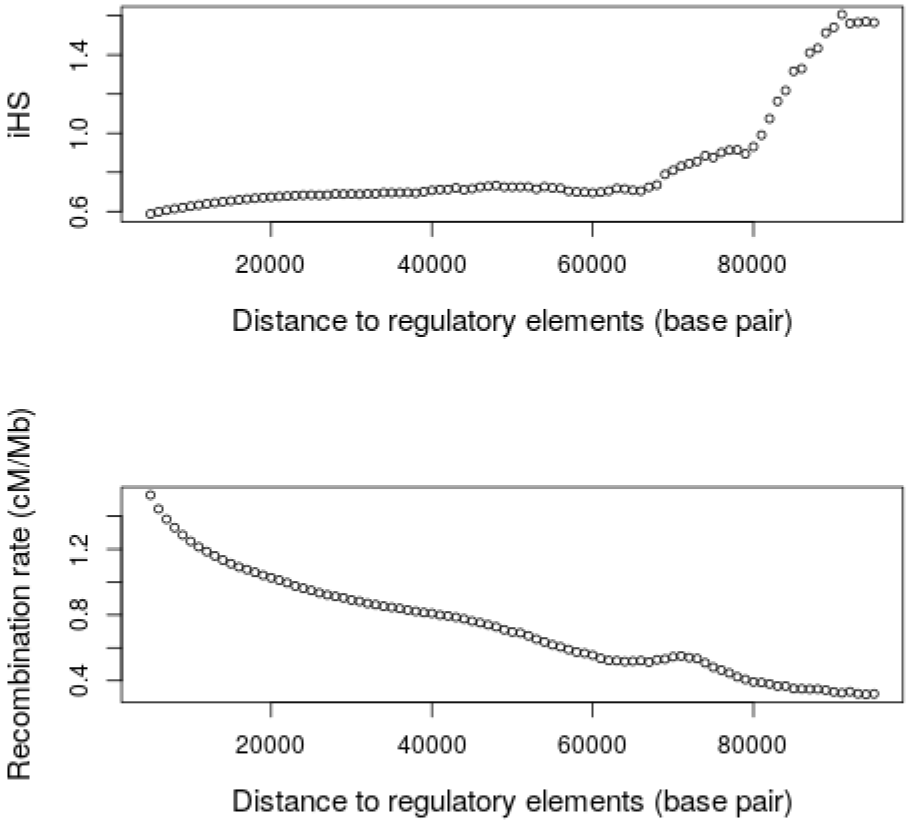


Figure S1: Association between distance to regulatory elements (ChIP-seq), iHS and recombination rate at a fine scale. For each of the studied SNPs in Yoruba, we calculated the distance to the center of the closest regulatory element. We also calculated the recombination rate in 100 kb windows centered around each SNP included in this study. We used the same approach than for the calculation of recombination rate in gene-centered windows (see Methods). We considered SNPs separated up to 100 kb from the closest regulatory element to focus on the fine scale patterns of selection and recombination around regulatory elements. We calculated a moving average using sliding windows of 10 kb with a separation of 1 kb. For example, we calculated the average iHS and recombination rate of all SNPs at distance to regulatory elements between 0 and 10 kb. In the next averages, we considered SNPs between 1 and 11 kb, 2 and 12 kb and so on.

The resulting averages are shown for iHS in the upper panel and recombination in the lower panel.

#### iHS, recombination rate and distance to regulatory elements in testis

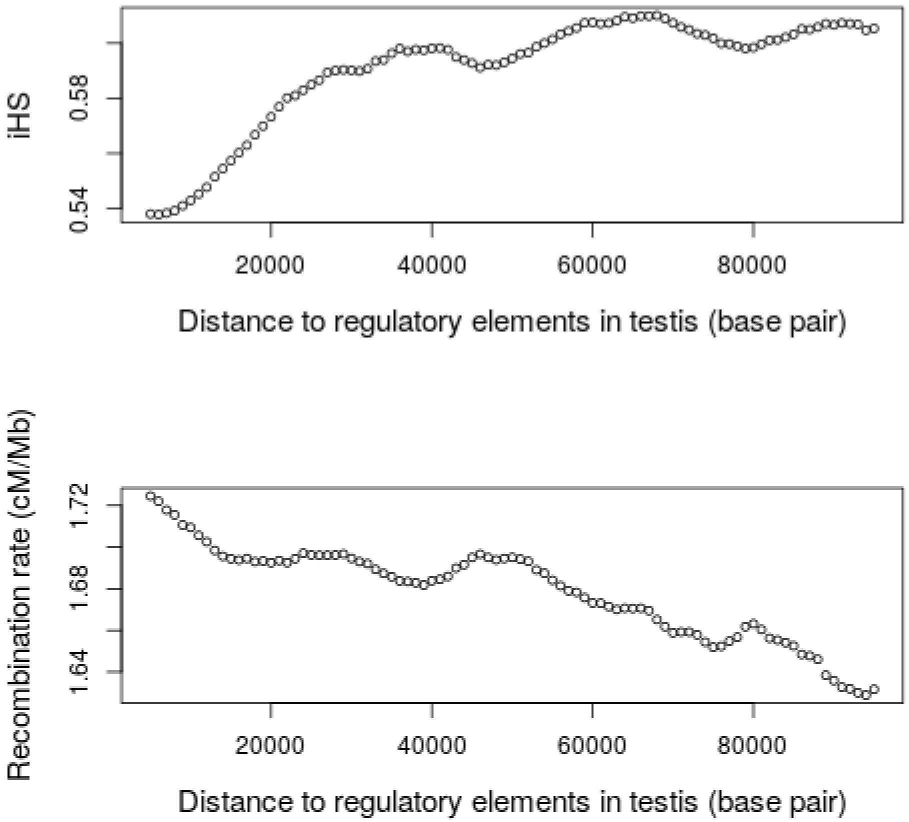


Figure S2: Association between distance to regulatory elements in testis (ChIP-seq), iHS and recombination rate at a fine scale. We used the same approach explained in Figure S1 but calculating the distance of each SNP to the center of the closest regulatory element in testis. The upper panel shows the association between the distance to regulatory elements in testis and iHS, while the lower panel show the association with recombination rate.

#### iHS, recombination rate and distance to regulatory elements in lymphocytes

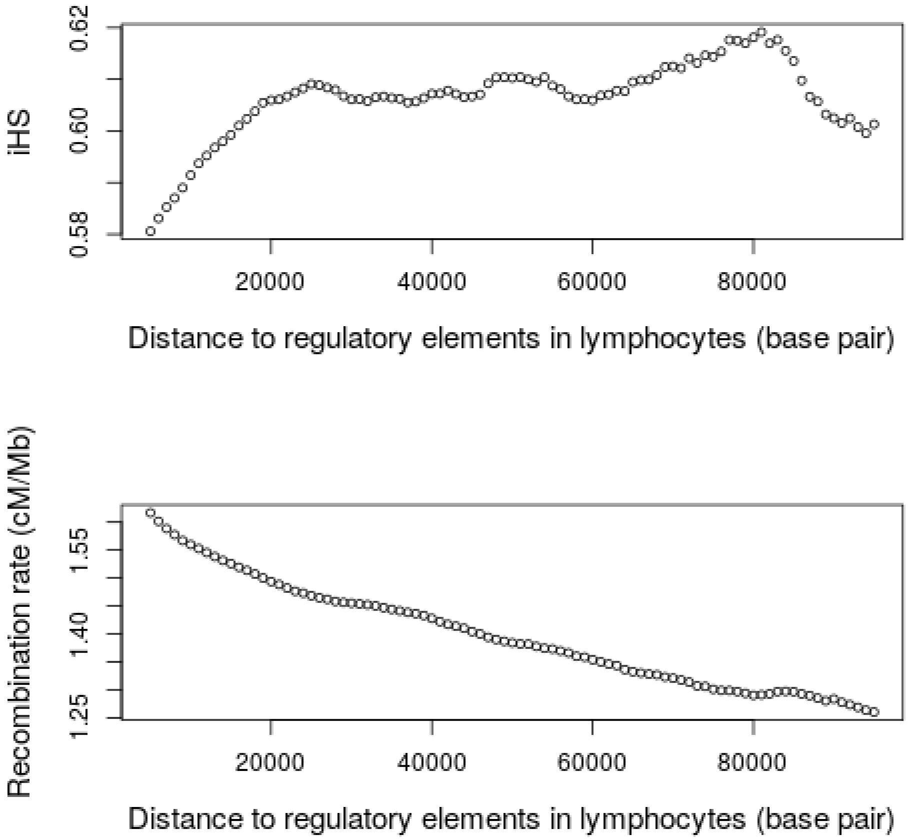


Figure S3: Association between distance to regulatory elements in lymphocytes (ChIP-seq), iHS and recombination rate at a fine scale. We used the same approach explained in Figure S1 but calculating the distance of each SNP to the center of the closest regulatory element in lymphocytes. The upper panel shows the association between the distance to regulatory elements in lymphocytes and iHS, while the lower panel show the association with recombination rate.

#### Recombination rate and the three regulatory densities

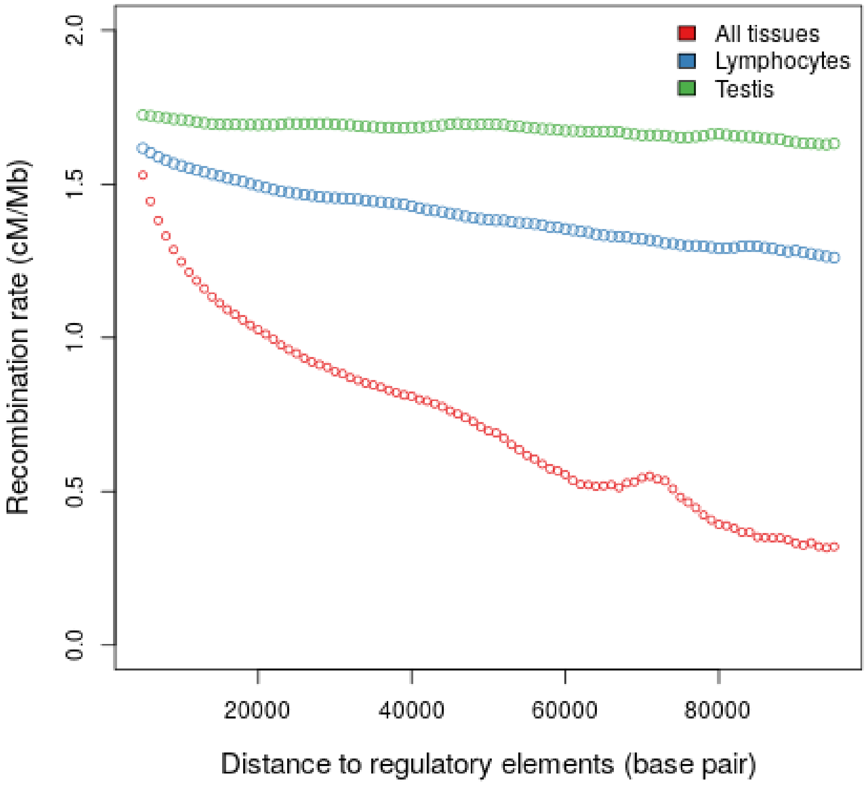


Figure S4: Association between recombination rate and the distance to regulatory elements in lymphocytes, testis and all tissues (ChIP-seq) at a fine scale. The plot shows the same average recombination rate previously calculated as a function of the distance to regulatory elements, but showing the three types of regulatory datasets at the same time.

#### iHS, recombination rate and distance to coding + regulatory elements

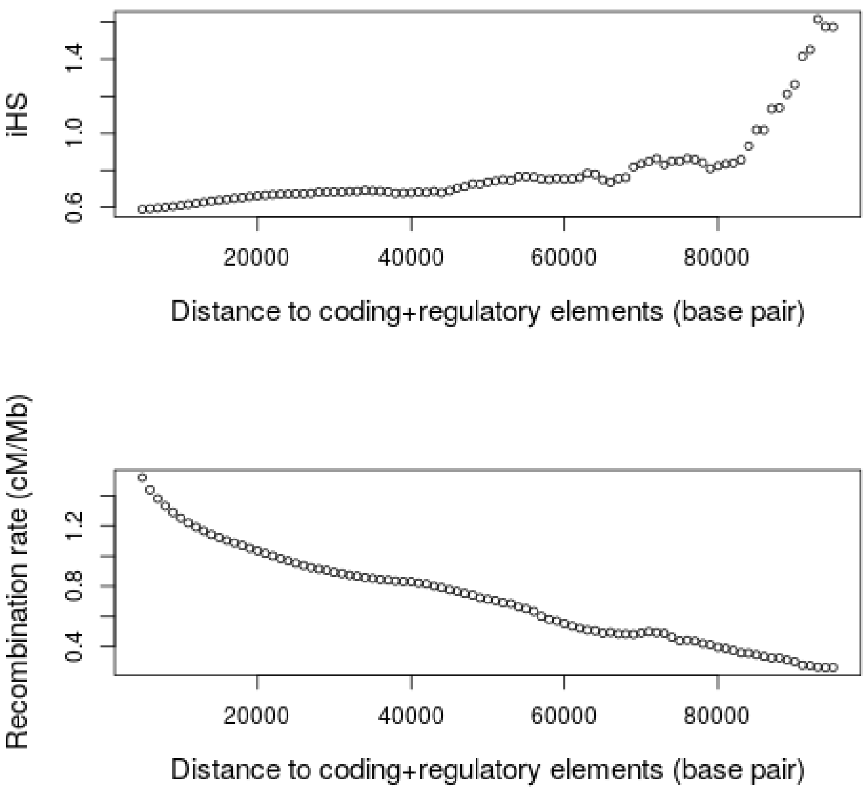


Figure S5: Association between distance to coding and regulatory elements (ChIP-seq), iHS and recombination rate at a fine scale. We used the same approach explained in Figure S1 but also considering the distance to center of coding elements. In other words, we calculated the distance between each SNP and the closest coding or regulatory element (whatever is closer). The upper panel shows the association between the distance to coding/regulatory elements and iHS, while the lower panel show the association with recombination rate.

#### Modeling results from fine scale-recombination analyses

Next sections include the results for analyses using variables about recombination around regulatory elements. We considered three sets of regulatory elements, that is, transcription factor binding sites across all tissues included in ChIP-seq experiments (UCSC Genome Browser), along with binding sites in lymphocytes and testis (see Methods for further details). In order to calculate the recombination around regulatory elements, we applied the following approach: For each set of regulatory elements, we created a 10 kb genomic window around the center of each element (5 kb at each side; regulatory window hereafter). We then fused those regulatory windows that were overlapped in at least 1 base. Next, recombination rate was calculated for the resulting regulatory windows. We followed the same approach than for gene windows, that is, calculating the ratio of genetic to physical distance between the ends of each regulatory window. In order to calculate the genetic position of each edge of the regulatory window, we again applied the same approach than in the case of gene windows. We searched for data about genetic position around each edge of the window up to 10 kb at each side, using then linear interpolation to estimate the genetic position of the corresponding edge (see Methods section for further details). Once the recombination rate around each regulatory window was estimated, we calculated the average of recombination of all regulatory windows completely overlapped with each gene window. In other words, we obtained a value of average recombination around regulatory elements for each gene window. We also counted the number of recombination data points, i.e., the number of regulatory windows for which recombination could be calculated in each gene window. In that way, we attempted to control for the potential noise caused by variability in the number of regulatory windows between gene windows.

We applied this approach separately for each regulatory variable, that is, considering the regulatory elements across all tissues, but also considering only regulatory elements in lymphocytes and testis in parallel calculations. Therefore, we obtained three different variables related to the recombination: recombination around regulatory elements in lymphocytes, testis and across all tissues. Each variable was included in the original model separately, being present only one recombination variable each time. We also run models that included one of these variables and the original recombination variable, that is, recombination rate across gene windows (window-wide scale hereafter). In that way, we attempted to control for the possibility that these new variables would not fully account for the influence of recombination at a wider scale. Similar results after including the original window-wide recombination variable would suggest that our results are not caused by a lack of control of recombination at a window-wide scale. Finally, we repeated the analyses considering only high recombination regions, that is, analyzing only genes with a recombination rate (at a window-wide level) equal or higher than the second tertile (1.552 cM/Mb). We further limited in this way the confounding effect caused by low-recombination regions, i.e., a mismatch between recombination at different scales: low at the window-wide level, but high at local level around regulatory elements. This mismatch should not exist in high recombination regions, where recombination is also high at the window-wide level. Only the Yoruba population and 1,000 kb windows were analyzed.

#### Yoruba

##### *Yoruba 1000kb: Recombination around lymphocytes regulatory elements without original recombination*

Figure S6: Mixture of Gaussian distributions fitting observed iHS (1000kb windows) for Yoruba. The figure shows the two Gaussian distributions, component 1 and 2, being the latter enriched in positive selection. In that component, iHS linearly depends on the genomic factors considered. Legend: Light blue = Observed iHS; Dark blue = Mixture model; Full red curve = Component 1 of the mixture model; Dashed red curve = Component 2 of the mixture model enriched in selection.

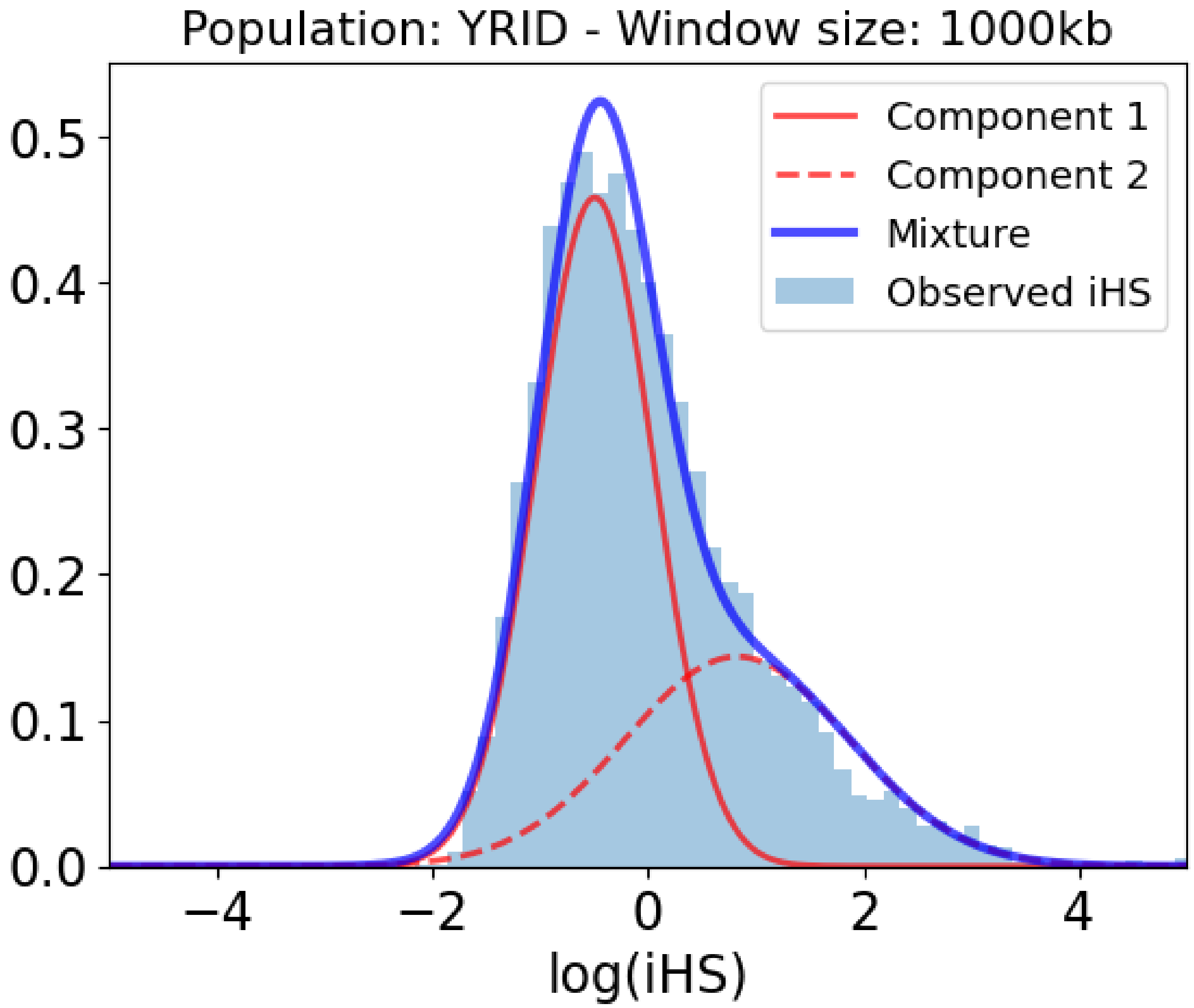


Table S1: Slopes and p-values of the association between iHS and genomic factors for the Yoruba population in 1000kb within the selection-enriched component.

| Covariate | Slope | P-value |
|---|---|---|
| Intercept | -0.724 | 0.000E+00 |
| Number iHS data points | -0.193 | 1.629E-10 |

| Covariate | Slope | P-value |
|---|---|---|
| Regulatory density (ChIP-seq) | -0.261 | 1.029E-02 |
| Regulatory density in immune cells (ChIP-seq) | 0.680 | 4.596E-14 |
| Recombination rate around regulatory elements in lymphocytes (ChIP-seq) | -0.451 | 0.000E+00 |
| Number of recombination rate data points around regulatory elements in lymphocytes (ChIP-seq) | -1.289 | 0.000E+00 |
| Regulatory density in testis (ChIP-seq) | -0.808 | 0.000E+00 |
| Coding density | 0.349 | 5.309E-04 |
| Density of conserved elements | 0.078 | 9.643E-02 |
| Gene expression | -0.114 | 8.713E-02 |
| GC-content | -0.118 | 2.687E-01 |
| Gene length | -0.100 | 7.104E-03 |
| Gene number | 0.056 | 5.203E-01 |
| Gene expression in immune cells | 0.273 | 6.407E-06 |
| Number PPIs | -0.108 | 1.736E-03 |
| Regulatory density (DNaseI) | 0.336 | 5.579E-03 |
| Gene expression in testis | 0.062 | 1.808E-01 |
| Distance to VIPs | -0.313 | 0.000E+00 |

#### *Yoruba 1000kb: Recombination around lymphocytes regulatory elements and original recombination*

Figure S7: Mixture of Gaussian distributions fitting observed iHS (1000kb windows) for Yoruba. The figure shows the two Gaussian distributions, component 1 and 2, being the latter enriched in positive selection. In that component, iHS linearly depends on the genomic factors considered. Legend: Light blue = Observed iHS; Dark blue = Mixture model; Full red curve = Component 1 of the mixture model; Dashed red curve = Component 2 of the mixture model enriched in selection.

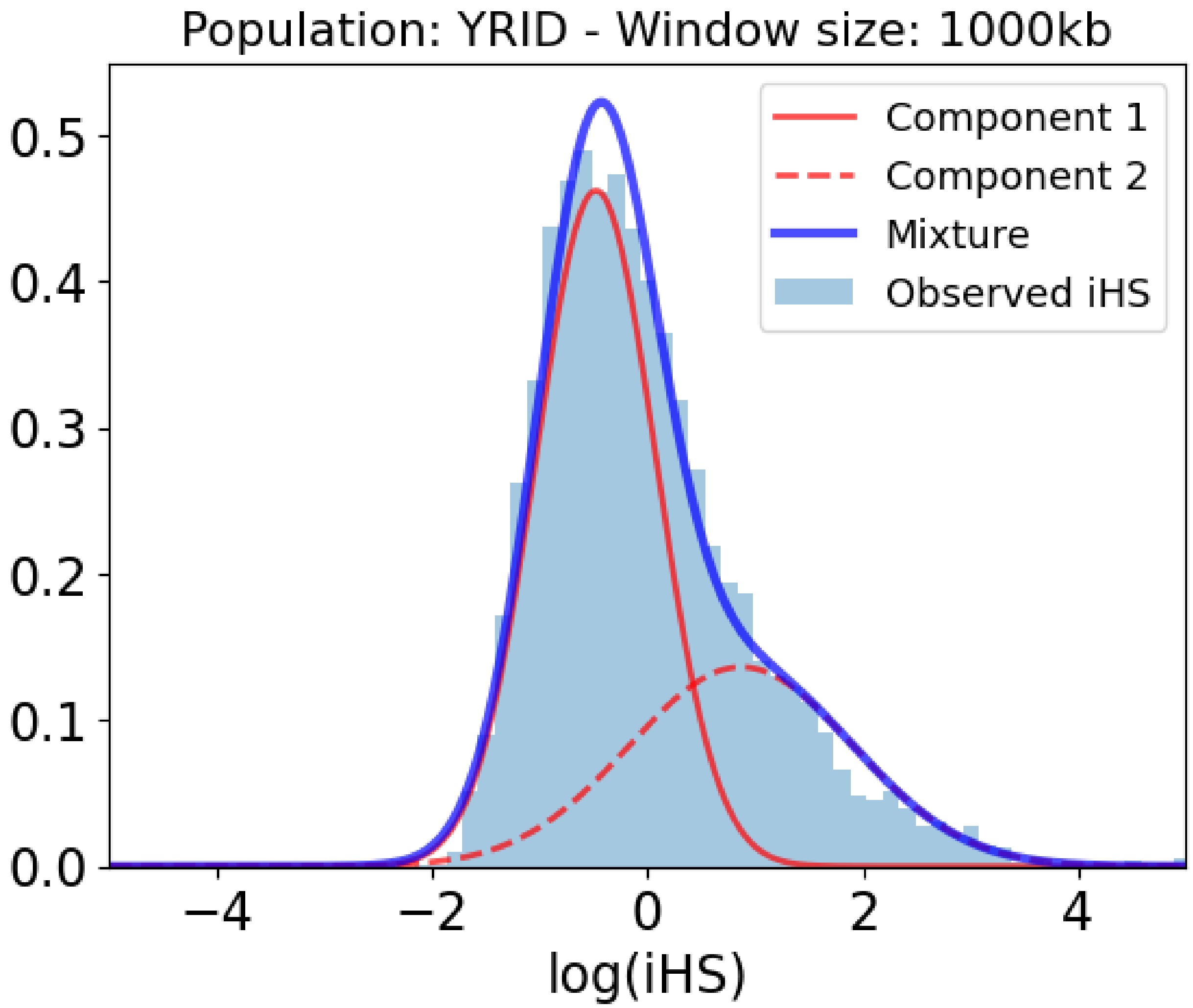


Table S2: Slopes and p-values of the association between iHS and genomic factors for the Yoruba population in 1000kb within the selection-enriched component.

| **Covariate** | **Slope** | **P-value** |
|---|---|---|
| Intercept | -1.179 | 0.000E+00 |
| Number iHS data points | 0.045 | 1.529E-01 |
| Regulatory density (ChIP-seq) | -0.073 | 5.309E-01 |
| Regulatory density in immune cells (ChIP-seq) | 0.275 | 5.717E-03 |
| Recombination rate around regulatory elements in lymphocytes (ChIP-seq) | 0.346 | 9.220E-10 |
| Number of recombination rate data points around regulatory elements in lymphocytes (ChIP-seq) | -0.757 | 0.000E+00 |
| Regulatory density in testis (ChIP-seq) | -0.909 | 0.000E+00 |
| Coding density | 0.144 | 1.789E-01 |

| Covariate | Slope | P-value |
|---|---|---|
| Density of conserved elements | 0.106 | 3.226E-02 |
| Gene expression | -0.086 | 2.223E-01 |
| GC-content | 0.390 | 1.036E-03 |
| Gene length | -0.085 | 2.672E-02 |
| Gene number | 0.020 | 8.623E-01 |
| Gene expression in immune cells | 0.219 | 6.323E-04 |
| Number PPIs | -0.088 | 1.773E-02 |
| Recombination rate | -2.110 | 0.000E+00 |
| Regulatory density (DNaseI) | 0.252 | 6.920E-02 |
| Gene expression in testis | 0.021 | 6.771E-01 |
| Distance to VIPs | -0.259 | 1.180E-10 |

##### *Yoruba 1000kb: Recombination around lymphocytes regulatory elements and original recombination in high recombination regions*

Figure S8: Mixture of Gaussian distributions fitting observed iHS (1000kb windows) for Yoruba. The figure shows the two Gaussian distributions, component 1 and 2, being the latter enriched in positive selection. In that component, iHS linearly depends on the genomic factors considered. Legend: Light blue = Observed iHS; Dark blue = Mixture model; Full red curve = Component 1 of the mixture model; Dashed red curve = Component 2 of the mixture model enriched in selection.

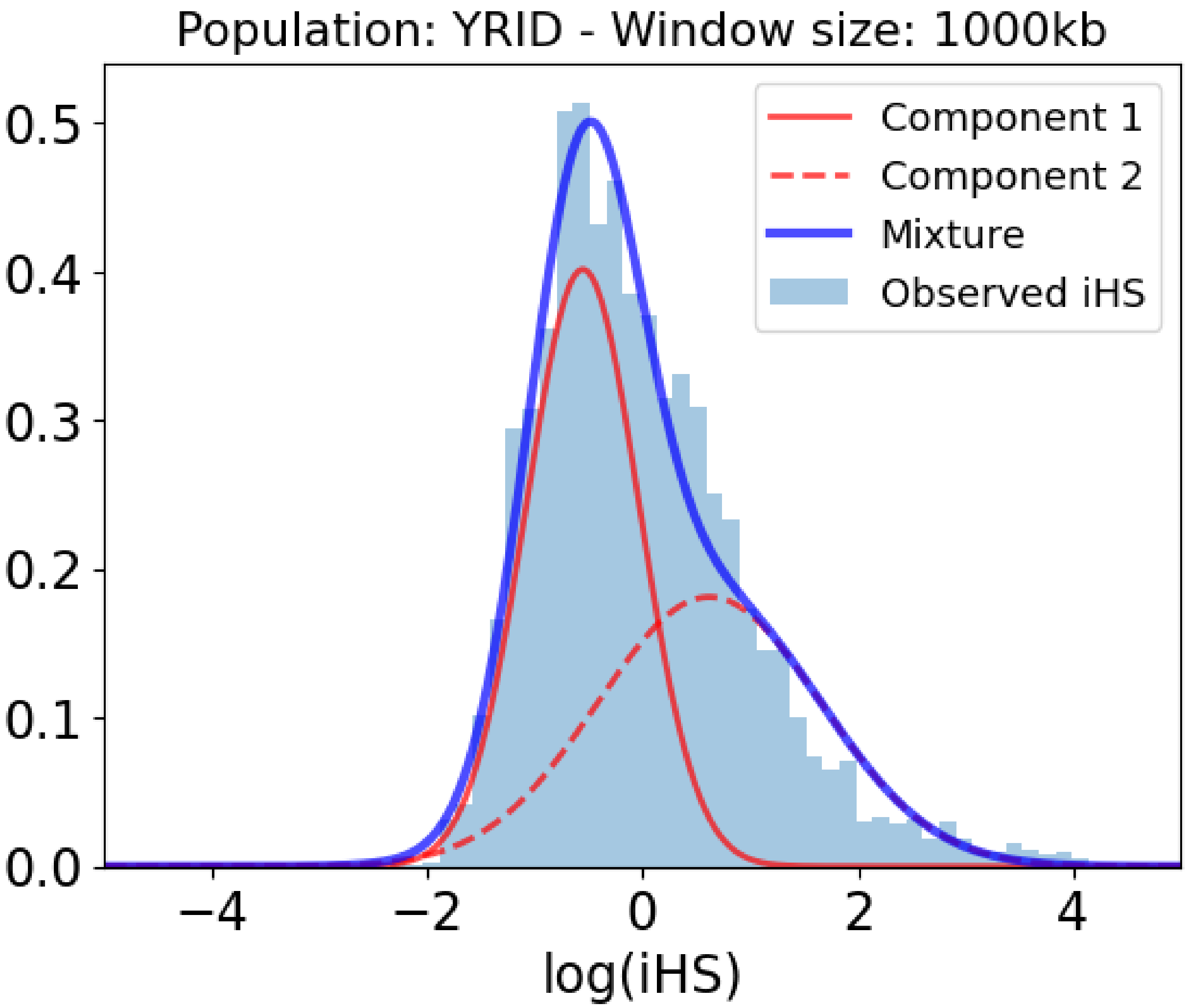


Table S3: Slopes and p-values of the association between iHS and genomic factors for the Yoruba population in 1000kb within the selection-enriched component.

| **Covariate** | **Slope** | **P-value** |
|---|---|---|
| Intercept | -0.145 | 4.730E-03 |
| Number iHS data points | -0.139 | 1.506E-02 |
| Regulatory density (ChIP-seq) | 0.496 | 9.345E-04 |
| Regulatory density in immune cells (ChIP-seq) | 0.846 | 1.267E-08 |
| Recombination rate around regulatory elements in lymphocytes (ChIP-seq) | 0.319 | 2.955E-06 |
| Number of recombination rate data points around regulatory elements in lymphocytes (ChIP-seq) | -0.890 | 0.000E+00 |
| Regulatory density in testis (ChIP-seq) | -0.483 | 1.498E-06 |
| Coding density | -0.092 | 5.957E-01 |

| Covariate | Slope | P-value |
|---|---|---|
| Density of conserved elements | -0.067 | 3.747E-01 |
| Gene expression | -0.286 | 7.175E-03 |
| GC-content | 0.546 | 1.056E-03 |
| Gene length | 0.100 | 1.016E-01 |
| Gene number | 0.090 | 5.836E-01 |
| Gene expression in immune cells | 0.221 | 2.202E-02 |
| Number PPIs | 0.025 | 6.538E-01 |
| Recombination rate | -0.859 | 0.000E+00 |
| Regulatory density (DNaseI) | -0.506 | 4.277E-03 |
| Gene expression in testis | 0.046 | 5.458E-01 |
| Distance to VIPs | -0.235 | 2.347E-04 |

#### *Yoruba 1000kb: Recombination around testis regulatory elements without original recombination*

Figure S9: Mixture of Gaussian distributions fitting observed iHS (1000kb windows) for Yoruba. The figure shows the two Gaussian distributions, component 1 and 2, being the latter enriched in positive selection. In that component, iHS linearly depends on the genomic factors considered. Legend: Light blue = Observed iHS; Dark blue = Mixture model; Full red curve = Component 1 of the mixture model; Dashed red curve = Component 2 of the mixture model enriched in selection.

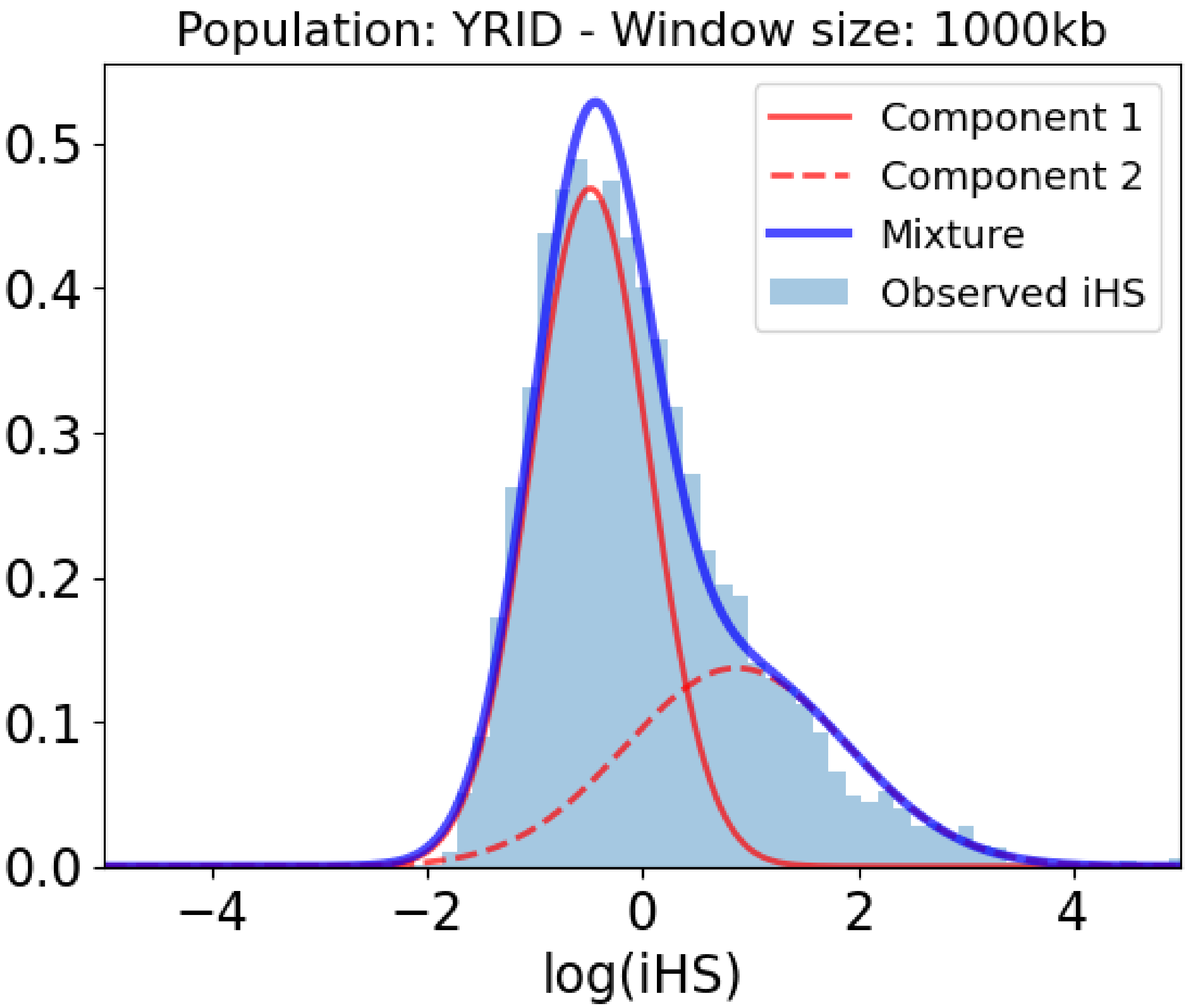


Table S4: Slopes and p-values of the association between iHS and genomic factors for the Yoruba population in 1000kb within the selection-enriched component.

| **Covariate** | **Slope** | **P-value** |
|---|---|---|
| Intercept | -0.759 | 0.000E+00 |
| Number iHS data points | -0.513 | 0.000E+00 |
| Regulatory density (ChIP-seq) | -0.273 | 2.761E-03 |
| Regulatory density in immune cells (ChIP-seq) | 0.348 | 1.221E-05 |
| Recombination rate around regulatory elements in testis (ChIP-seq) | -0.145 | 5.775E-07 |
| Number of recombination rate data points around regulatory elements in testis (ChIP-seq) | -1.123 | 0.000E+00 |
| Regulatory density in testis (ChIP-seq) | 0.063 | 3.198E-01 |
| Coding density | 0.389 | 6.015E-06 |
| Density of conserved elements | 0.194 | 2.905E-06 |

| Covariate | Slope | P-value |
|---|---|---|
| Gene expression | -0.056 | 3.396E-01 |
| GC-content | -0.206 | 2.687E-02 |
| Gene length | -0.075 | 2.275E-02 |
| Gene number | 0.008 | 9.198E-01 |
| Gene expression in immune cells | 0.149 | 5.850E-03 |
| Number PPIs | -0.087 | 4.562E-03 |
| Regulatory density (DNaseI) | 0.115 | 2.891E-01 |
| Gene expression in testis | 0.065 | 1.136E-01 |
| Distance to VIPs | -0.254 | 9.681E-14 |

#### *Yoruba 1000kb: Recombination around testis regulatory elements and original recombination*

Figure S10: Mixture of Gaussian distributions fitting observed iHS (1000kb windows) for Yoruba. The figure shows the two Gaussian distributions, component 1 and 2, being the latter enriched in positive selection. In that component, iHS linearly depends on the genomic factors considered. Legend: Light blue = Observed iHS; Dark blue = Mixture model; Full red curve = Component 1 of the mixture model; Dashed red curve = Component 2 of the mixture model enriched in selection.

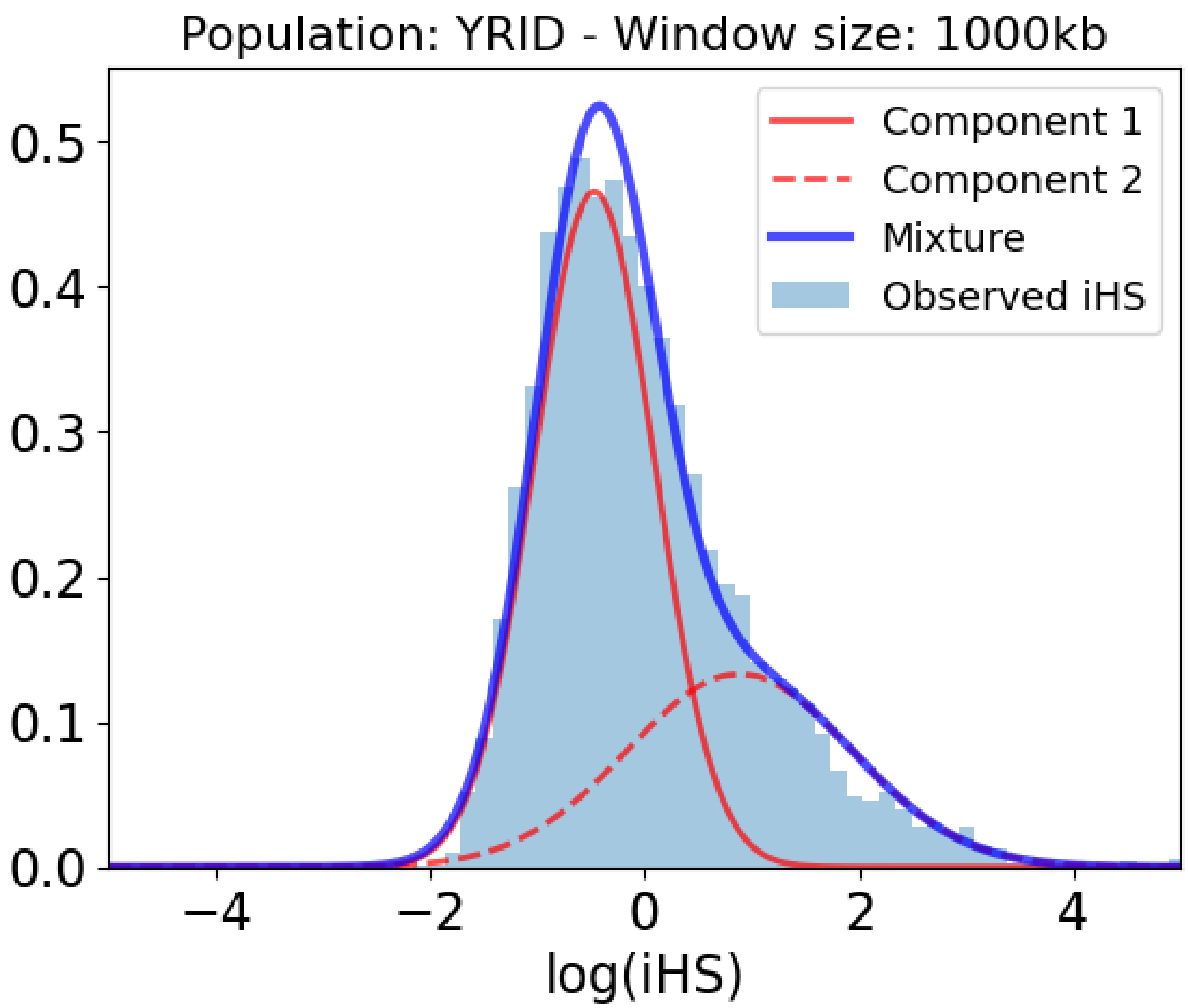


Table S5: Slopes and p-values of the association between iHS and genomic factors for the Yoruba population in 1000kb within the selection-enriched component.

| **Covariate** | **Slope** | **P-value** |
|---|---|---|
| Intercept | -1.276 | 0.000E+00 |
| Number iHS data points | 0.026 | 4.333E-01 |
| Regulatory density (ChIP-seq) | -0.188 | 1.065E-01 |
| Regulatory density in immune cells (ChIP-seq) | 0.132 | 1.644E-01 |
| Recombination rate around regulatory elements in testis (ChIP-seq) | 0.242 | 3.020E-08 |
| Number of recombination rate data points around regulatory elements in testis (ChIP-seq) | -0.497 | 1.040E-11 |
| Regulatory density in testis (ChIP-seq) | -0.550 | 1.548E-13 |
| Coding density | 0.195 | 4.638E-02 |
| Density of conserved elements | 0.179 | 2.236E-04 |

| Covariate | Slope | P-value |
|---|---|---|
| Gene expression | -0.092 | 1.810E-01 |
| GC-content | 0.402 | 4.419E-04 |
| Gene length | -0.067 | 7.885E-02 |
| Gene number | -0.071 | 4.578E-01 |
| Gene expression in immune cells | 0.178 | 4.423E-03 |
| Number PPIs | -0.071 | 4.914E-02 |
| Recombination rate | -2.266 | 0.000E+00 |
| Regulatory density (DNaseI) | 0.176 | 2.050E-01 |
| Gene expression in testis | 0.016 | 7.354E-01 |
| Distance to VIPs | -0.210 | 4.949E-08 |

#### *Yoruba 1000kb: Recombination around testis regulatory elements and original recombination in high recombination regions*

Figure S11: Mixture of Gaussian distributions fitting observed iHS (1000kb windows) for Yoruba. The figure shows the two Gaussian distributions, component 1 and 2, being the latter enriched in positive selection. In that component, iHS linearly depends on the genomic factors considered. Legend: Light blue = Observed iHS; Dark blue = Mixture model; Full red curve = Component 1 of the mixture model; Dashed red curve = Component 2 of the mixture model enriched in selection.

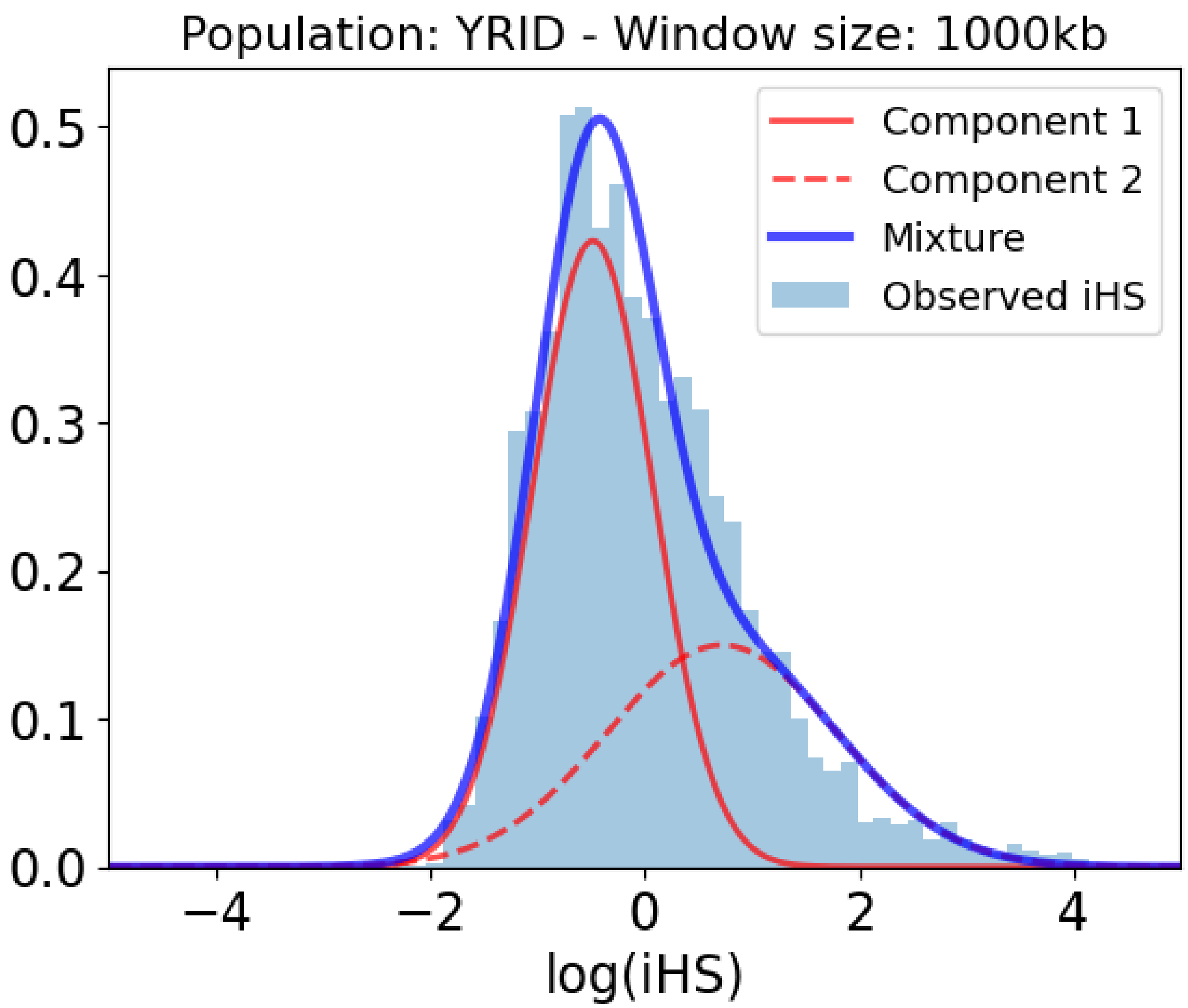


Table S6: Slopes and p-values of the association between iHS and genomic factors for the Yoruba population in 1000kb within the selection-enriched component.

| **Covariate** | **Slope** | **P-value** |
|---|---|---|
| Intercept | -0.483 | 0.000E+00 |
| Number iHS data points | -0.003 | 9.615E-01 |
| Regulatory density (ChIP-seq) | 0.427 | 2.304E-03 |
| Regulatory density in immune cells (ChIP-seq) | 0.366 | 5.361E-03 |
| Recombination rate around regulatory elements in testis (ChIP-seq) | 0.369 | 1.524E-10 |
| Number of recombination rate data points around regulatory elements in testis (ChIP-seq) | -1.246 | 0.000E+00 |
| Regulatory density in testis (ChIP-seq) | 0.585 | 4.983E-05 |
| Coding density | -0.295 | 9.542E-02 |
| Density of conserved elements | 0.038 | 6.080E-01 |

| Covariate | Slope | P-value |
|---|---|---|
| Gene expression | -0.208 | 4.748E-02 |
| GC-content | 0.689 | 2.563E-05 |
| Gene length | 0.071 | 2.858E-01 |
| Gene number | 0.178 | 2.776E-01 |
| Gene expression in immune cells | 0.098 | 3.054E-01 |
| Number PPIs | 0.018 | 7.504E-01 |
| Recombination rate | -0.854 | 0.000E+00 |
| Regulatory density (DNaseI) | -0.548 | 1.262E-03 |
| Gene expression in testis | 0.046 | 5.309E-01 |
| Distance to VIPs | -0.261 | 1.734E-04 |

##### *Yoruba 1000kb: Recombination around regulatory elements without original recombination*

Figure S12: Mixture of Gaussian distributions fitting observed iHS (1000kb windows) for Yoruba. The figure shows the two Gaussian distributions, component 1 and 2, being the latter enriched in positive selection. In that component, iHS linearly depends on the genomic factors considered. Legend: Light blue = Observed iHS; Dark blue = Mixture model; Full red curve = Component 1 of the mixture model; Dashed red curve = Component 2 of the mixture model enriched in selection.

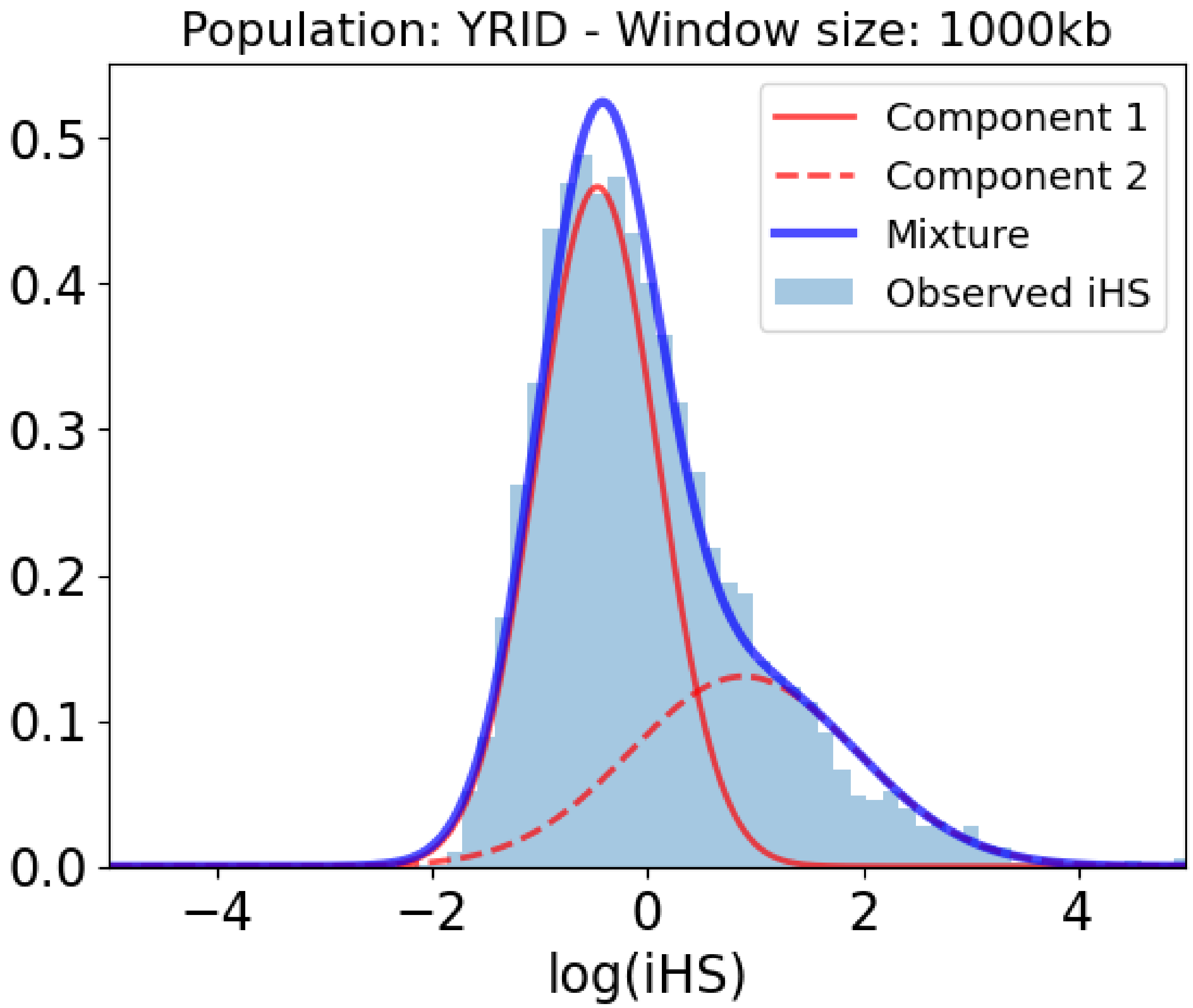


Table S7: Slopes and p-values of the association between iHS and genomic factors for the Yoruba population in 1000kb within the selection-enriched component.

| Covariate | Slope | P-value |
|---|---|---|
| Intercept | -1.078 | 0.000E+00 |
| Number iHS data points | 0.044 | 1.602E-01 |
| Regulatory density (ChIP-seq) | -1.955 | 0.000E+00 |
| Regulatory density in immune cells (ChIP-seq) | 0.487 | 1.563E-07 |
| Recombination rate around regulatory elements (ChIP-seq) | -0.690 | 0.000E+00 |
| Number of recombination rate data points around regulatory elements (ChIP-seq) | -1.835 | 0.000E+00 |
| Regulatory density in testis (ChIP-seq) | -0.659 | 0.000E+00 |
| Coding density | -0.002 | 9.713E-01 |
| Density of conserved elements | 0.382 | 3.519E-14 |
| Gene expression | 0.038 | 5.961E-01 |

| Covariate | Slope | P-value |
| --- | --- | --- |
| GC-content | 0.238 | 5.356E-02 |
| Gene length | -0.049 | 2.358E-01 |
| Gene number | 0.212 | 3.837E-02 |
| Gene expression in immune cells | 0.052 | 4.408E-01 |
| Number PPIs | -0.054 | 1.461E-01 |
| Regulatory density (DNaseI) | -0.242 | 7.496E-02 |
| Gene expression in testis | 0.002 | 9.728E-01 |
| Distance to VIPs | -0.238 | 8.612E-08 |

#### *Yoruba 1000kb: Recombination around regulatory elements and original recombination*

Figure S13: Mixture of Gaussian distributions fitting observed iHS (1000kb windows) for Yoruba. The figure shows the two Gaussian distributions, component 1 and 2, being the latter enriched in positive selection. In that component, iHS linearly depends on the genomic factors considered. Legend: Light blue = Observed iHS; Dark blue = Mixture model; Full red curve = Component 1 of the mixture model; Dashed red curve = Component 2 of the mixture model enriched in selection.

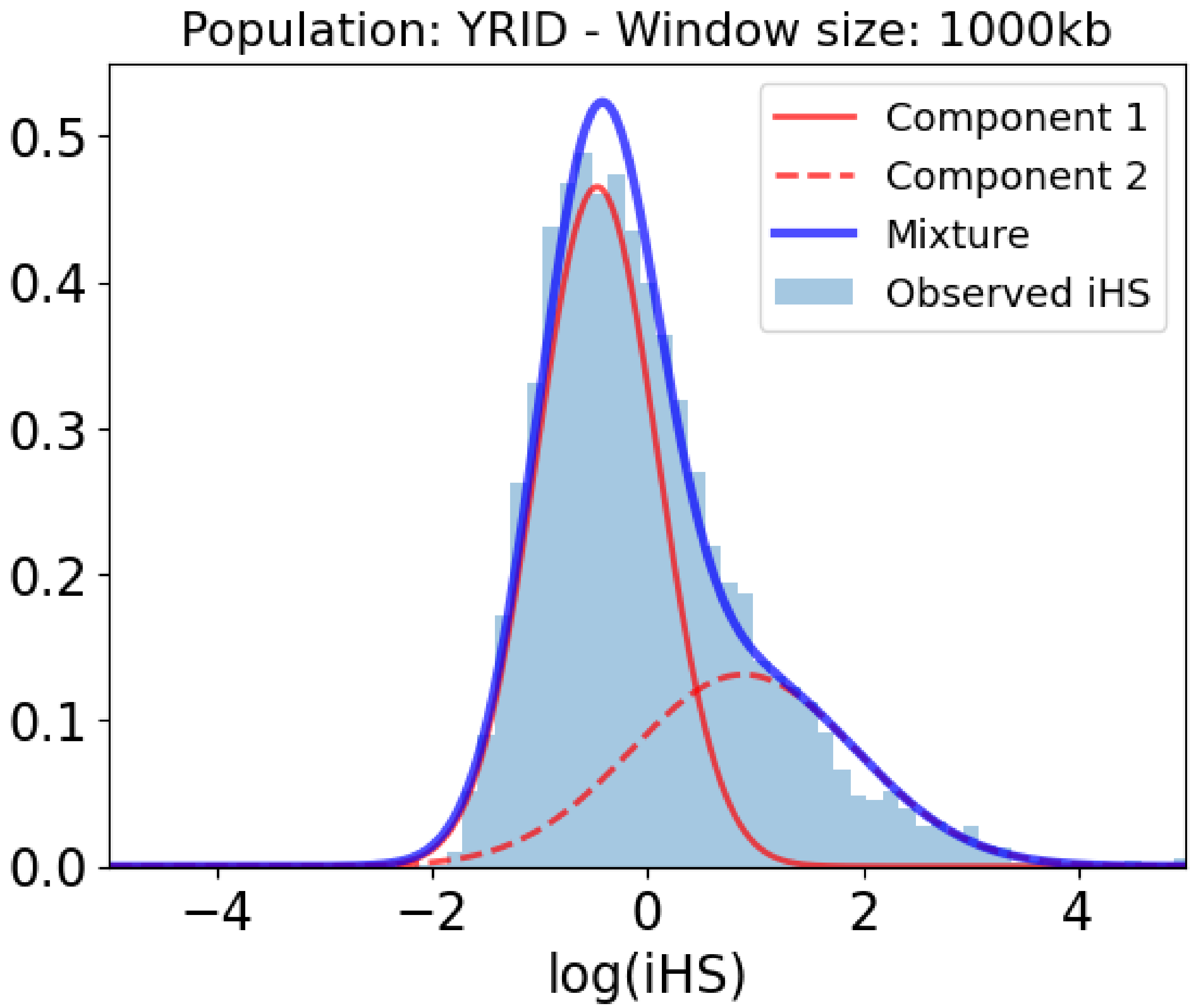


Table S8: Slopes and p-values of the association between iHS and genomic factors for the Yoruba population in 1000kb within the selection-enriched component.

| **Covariate** | **Slope** | **P-value** |
|---|---|---|
| Intercept | -1.253 | 0.000E+00 |
| Number iHS data points | 0.161 | 1.396E-06 |
| Regulatory density (ChIP-seq) | -1.322 | 0.000E+00 |
| Regulatory density in immune cells (ChIP-seq) | 0.188 | 5.430E-02 |
| Recombination rate around regulatory elements (ChIP-seq) | 0.044 | 5.366E-01 |
| Number of recombination rate data points around regulatory elements (ChIP-seq) | -1.435 | 0.000E+00 |
| Regulatory density in testis (ChIP-seq) | -0.801 | 0.000E+00 |
| Coding density | -0.038 | 6.941E-01 |
| Density of conserved elements | 0.340 | 1.331E-10 |
| Gene expression | 0.006 | 9.245E-01 |

| Covariate | Slope | P-value |
|---|---|---|
| GC-content | 0.305 | 1.703E-02 |
| Gene length | -0.053 | 2.168E-01 |
| Gene number | 0.173 | 9.452E-02 |
| Gene expression in immune cells | 0.068 | 3.229E-01 |
| Number PPIs | -0.056 | 1.532E-01 |
| Recombination rate | -1.515 | 0.000E+00 |
| Regulatory density (DNaseI) | 0.007 | 9.572E-01 |
| Gene expression in testis | -0.014 | 7.941E-01 |
| Distance to VIPs | -0.226 | 4.450E-07 |

#### *Yoruba 1000kb: Recombination around testis regulatory elements and original recombination in high recombination regions*

Figure S14: Mixture of Gaussian distributions fitting observed iHS (1000kb windows) for Yoruba. The figure shows the two Gaussian distributions, component 1 and 2, being the latter enriched in positive selection. In that component, iHS linearly depends on the genomic factors considered. Legend: Light blue = Observed iHS; Dark blue = Mixture model; Full red curve = Component 1 of the mixture model; Dashed red curve = Component 2 of the mixture model enriched in selection.

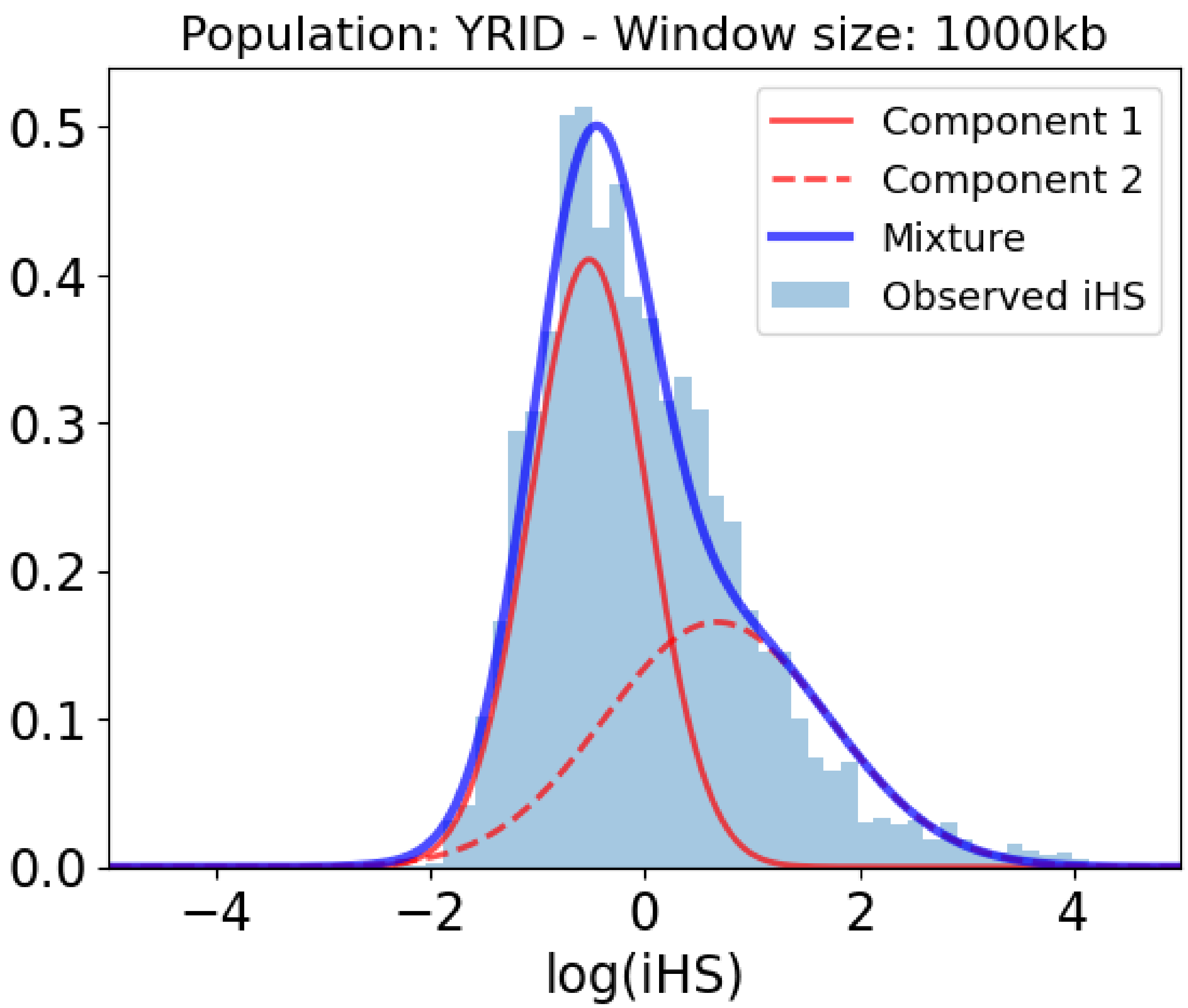


Table S9: Slopes and p-values of the association between iHS and genomic factors for the Yoruba population in 1000kb within the selection-enriched component.

| **Covariate** | **Slope** | **P-value** |
|---|---|---|
| Intercept | -0.337 | 3.131E-11 |
| Number iHS data points | -0.045 | 4.329E-01 |
| Regulatory density (ChIP-seq) | -0.392 | 9.898E-03 |
| Regulatory density in immune cells (ChIP-seq) | 0.417 | 1.759E-03 |
| Recombination rate around regulatory elements (ChIP-seq) | 0.178 | 1.230E-02 |
| Number of recombination rate data points around regulatory elements (ChIP-seq) | -0.921 | 0.000E+00 |
| Regulatory density in testis (ChIP-seq) | -0.344 | 3.891E-04 |
| Coding density | 0.053 | 7.621E-01 |
| Density of conserved elements | -0.050 | 4.960E-01 |
| Gene expression | -0.246 | 1.793E-02 |

| Covariate | Slope | P-value |
|---|---|---|
| GC-content | 0.819 | 1.352E-06 |
| Gene length | 0.138 | 3.260E-02 |
| Gene number | 0.004 | 9.817E-01 |
| Gene expression in immune cells | 0.102 | 2.749E-01 |
| Number PPIs | 0.038 | 4.942E-01 |
| Recombination rate | -0.735 | 0.000E+00 |
| Regulatory density (DNaseI) | -0.878 | 1.660E-05 |
| Gene expression in testis | 0.068 | 3.611E-01 |
| Distance to VIPs | -0.134 | 4.261E-02 |
