## Supplemental Results S5 for "Mixture Density Regression reveals frequent recent adaptation in the human genome"

### SLiM simulations results

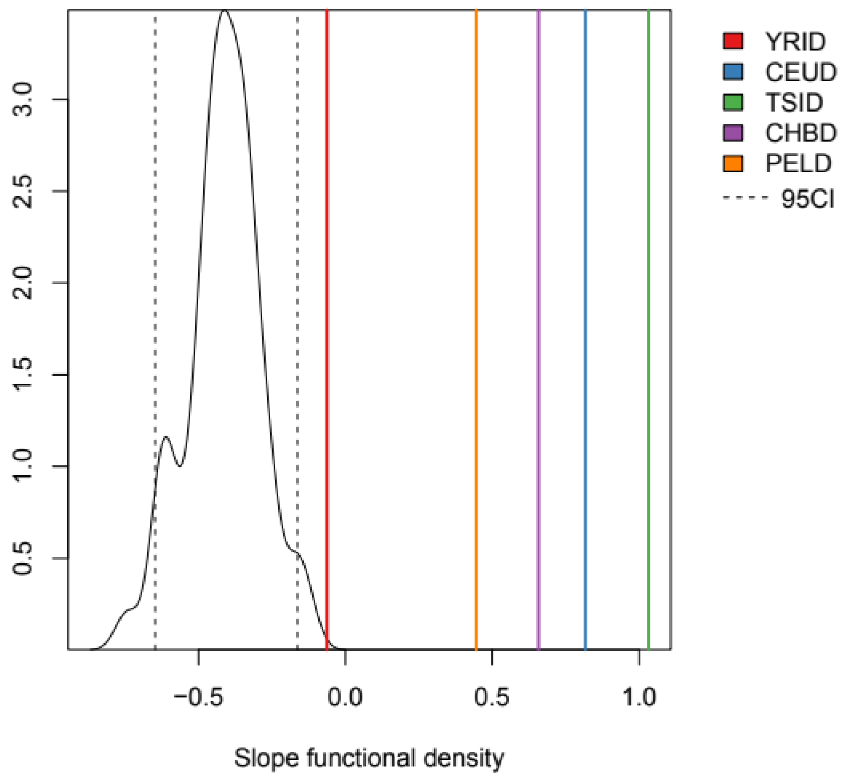


Figure S1: Distribution of the slope of functional (coding + regulatory) density across the 100 neutral simulations. The 95% confidence interval of this slope is shown with dashed lines. Moreover, the slope of functional density observed in the actual populations is represented with different vertical lines. These slopes are calculated using the original set of predictors in each population but summing coding and regulatory density into a single predictor. Abbreviations: YRID = Yoruba; CEUD = Utah residents; TSID = Toscani; CHBD = Han Chinse; PELD = Peruvians.

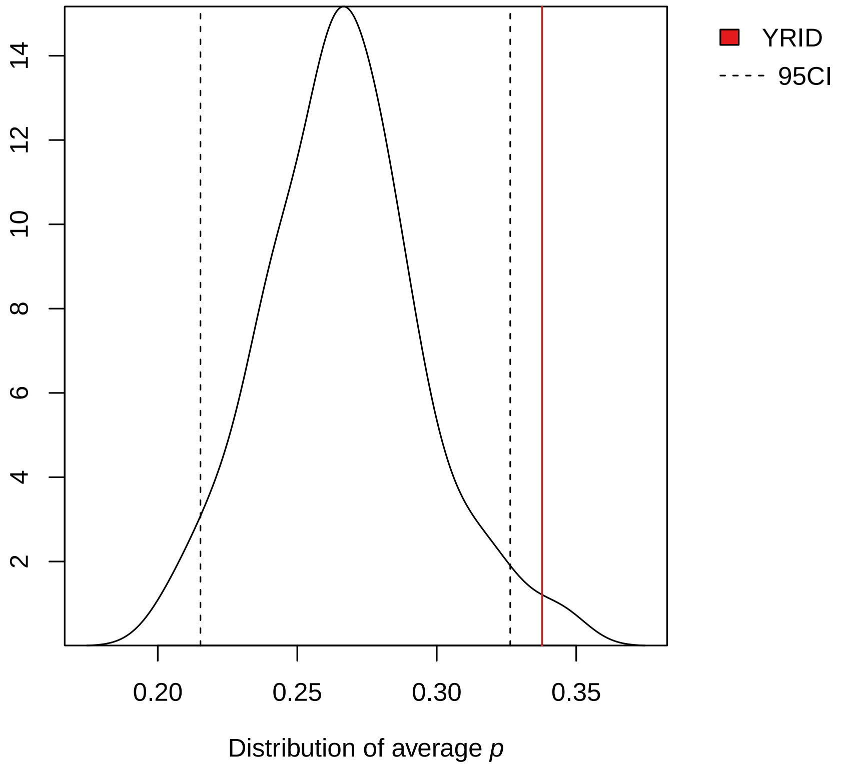


Figure S2: Distribution of average *p*. The parameter *p* can be regarded as the accumulative effect of all genomic factors on the probability of the second component of iHS (see Methods). In each neutral simulation, we calculated the average of *p* across all gene windows, obtaining a total of 100 values. We have used this value as a metric for the magnitude of the second component of iHS. The 95% confidence interval of this parameter is shown with dashed lines. Moreover, the value observed for this parameter in Yoruba is represented with a red solid line. Abbreviations: YRID = Yoruba.
